# Bioinformatics Analysis of next generation sequencing data for Risk Prediction in Patients with Type 1 diabetes mellitus

**DOI:** 10.1101/2022.03.17.484749

**Authors:** Basavaraj Vastrad, Chanabasayya Vastrad

## Abstract

Type 1 diabetes mellitus (T1DM) comprise the most common forms of autoimmune disease. The aim of this investigation was to apply a bioinformatics approach to reveal related pathways or genes involved in the development of T1DM. The next generation sequencing (NGS) dataset GSE182870 was downloaded from the gene expression omnibus (GEO) database. Differentially expressed gene (DEG) analysis was performed using DESeq2. The g:Profiler was utilized to analyze the functional enrichment, gene ontology (GO) and REACTOME pathway of the differentially expressed genes. Protein-protein interaction (PPI) network, modules, miRNA- hub gene regulatory network, and TF-hub gene regulatory network were conducted via comprehensive target prediction and network analyses. Finally, hub genes were validated by using receiver operating characteristic curve analysis. A total of 860 DEGs were screened out from NGS dataset, among which 477 genes were up regulated and 383 genes were down regulated. GO enrichment analysis indicated that up regulated genes were mainly involved in cellular metabolic process, intracellular anatomical structure and catalytic activity, and the down regulated genes were significantly enriched in cellular nitrogen compound biosynthetic process, protein containing complex and heterocyclic compound binding. REACTOME pathway enrichment analysis showed that the up regulated genes were mainly enriched in metabolism of carbohydrates and the down regulated genes were significantly enriched in metabolism of RNA. A PPI network, modules, miRNA-hub gene regulatory network and TF - hub gene regulatory network were constructed, and hub genes (MAPK14, RHOC, MAD2L1, TAF1, TRAF2, HSP90AA1, TP53, HSP90AB1, UBA52 and RACK1) were identified. This study provides further insights into the underlying pathogenesis of T1DM.

## Introduction

Type 1 diabetes mellitus (T1DM) is a auto immune endocrine and metabolic disease in children and adolescents, there is expected to increase by 50% in 20 years and reach 55 million by 2030 in the developed countries [1]. It is characterized by autoimmune destruction of insulin-producing beta cells results in absolute or insufficient insulin secretion [2]. Accumulating studies showed that T1DM develops as a manifestation of systemic illnesses that are linked to hypertension [3], cognitive diseases [4], obesity [5], renal diseases [6] and cardiovascular diseases [7]. The occurrence and development of T1DM are correlated with multiple factors includes genetic [8] and environmental factors [9]. Due to the lack of effective treatments, T1DM remains an incurable disease for the vast majority of patients [10]. Consequently, it needs more effort to clarify the molecular mechanism underlying T1DM advancement and progression, holding promise for finding potential drug targets and diagnostic biomarkers of T1DM.

Previous investigation identified aspects of the molecular mechanism of T1DM advancement. T1DM can be related to an altered expression of genes include transporter associated with antigen processing 1 (TAP1) [11], interleukin 21 (IL21) [12], protein tyrosine phosphatase non-receptor type 2 (PTPN2) [13], interleukin 10 (IL10) [14] and interleukin-1alpha (IL-1A) [15]. Several studies have recently suggested signaling pathways that may be responsible for the development of T1DM, including T cell receptor signaling pathway [16], NLRP3 and NLRP1 inflammasomes signaling pathway [17], Keap1/Nrf2 signaling pathway [18], mTORC signaling pathway [19] and type I IFN/STAT signaling pathway [20]. Despite the increasing insights into several aspects of molecular mechanism, a comprehensive overview of the integrated biological landscape underlying T1DM is currently missing.

Gene expression analysis based on next generation sequencing (NGS) technology is an extensively used, high-throughput and powerful investigation method, which can simultaneously detect expression change of thousands of genes. NGS investigations have found genes which played a key role in T1DM initiation and development and could be assessed as potential molecular targets and diagnostic markers [21]. In the process of this research on T1DM, based on this technology, we can detect and explore the gene expression of T1DM at the molecular level.

Bioinformatics analysis offers an ideal way to screen NGS dataset to comprehensively understand the molecular mechanisms underlying T1DM. In this investigation, we used a bioinformatics analysis approach to detect key genes and potential new biomarkers involved in T1DM. Molecular mechanisms underlying T1DM were also explored to search for possible new treatment targets for T1DM.

## Materials and methods

### Data resources

The NGS dataset of GSE182870 [22] were obtained from the National Centre of Biotechnology Information (NCBI) Gene Expression Omnibus database (GEO, https://www.ncbi.nlm.nih.gov/geo/) [23]. GSE182870, which comprises a total of 780 samples, including 600 T1DM samples, and 180 normal control samples, was based on the platform of the GPL18573 Illumina NextSeq 500 (Homo sapiens).

### Identification of DEGs

Differentially expressed genes (DEGs) between T1DM samples and normal control samples were identified by analyzing NGS data with DESeq2 package of R software [24]. The Benjamini and Hochberg (BH) method was accomplished to adjust P value to reduce the false discover rate [25]. According to the standard, we used FC > 3.835 as the screening criterion for up regulation of DEGs, FC < < 0 for down regulation of DEGs and adjust P value was < 0.05. Volcano plot and heat L map were generated using ggplot2 and gplot in R Bioconductor.

### GO and pathway enrichment analyses of DEGs

Gene Ontology (GO) enrichment (http://www.geneontology.org) [26] analysis was used for the examination of functional roles of gene sets, while REACTOME pathway (https://reactome.org/) [27] enrichment analyses were used to classify the pathways in which such genes might function. GO term is a used to express the features of genes and gene products, including three parts: biological process (BP), cellular component (CC), and molecular function (MF). In this investigation, to clarify the biological functions of DEGs, based on the g:Profiler (http://biit.cs.ut.ee/gprofiler/) [28], we performed enrichment analysis of DEGs in GO and REACTOME. The enriched GO terms and pathways were captured to analyze the DEGs at the functional level with the setting p < 0.05.

### Construction of the protein-protein interaction (PPI) and module analysis

DEGs were submitted to Search Tool for the Retrieval of Interacting Genes (STRING, https://string-db.org/) [29] to investigate interactions among the proteins encoded by the DEGs. Then, Cytoscape software version 3.9.1 (http://www.cytoscape.org/) [30] was used for construction of PPI network. The PPI network for hub genes was computed with the node degree [31] , betweenness [32], stress [33] and closeness [34] methods and Cytoscape plug-in Network Analyzer. PEWCC1 [35] was used to identify the representative modules using Cytoscape software.

### miRNA-hub gene regulatory network construction

The miRNA-hub gene was downloaded from the miRNet database (https://www.mirnet.ca/) [36]. The expressed miRNA in T1DM, which could regulate hub genes in T1DM, were identified as key miRNAs. The expressed miRNAs and hub genes were used to construct the miRNA-hub gene regulatoury network using Cytoscape software [30].

### TF-hub gene regulatory network construction

The TF-hub gene was downloaded from the NetworkAnalyst database (https://www.networkanalyst.ca/) [37]. The expressed TF in T1DM, which could regulate hub genes in T1DM, were identified as key TFs. The expressed TFs and hub genes were used to construct the TF-hub gene regulatory network using Cytoscape software [30].

### Receiver operating characteristic curve (ROC) analysis

Genes in the PPI network, modules, miRNA-hub gene network regulatory and TF-hub gene regulatory network were selected as candidate hub genes. We performed a ROC analysis and calculate area under curve (AUC) with mRNA expression value in GSE182870 to investigate the diagnostic value of hub genes in T1DM using “pROC” R package [38]. AUC value reflects the sensitivity and specificity of a hub gene in distinguishing T1DM from health. Here, we believed that hub genes with AUC greater than 0.8 can be used as biomarkers for the diagnosis of T1DM.

## Results

### Identification of DEGs

A total of 860 DEGs between T1DM and normal control were finally screened out according to the above criteria, including 477 up regulated genes and 383 down regulated genes (Table 1). The DEGs were visualized by volcano plot and heat map, as shown in Fig. 1 and Fig. 2.

**Fig. 1.**
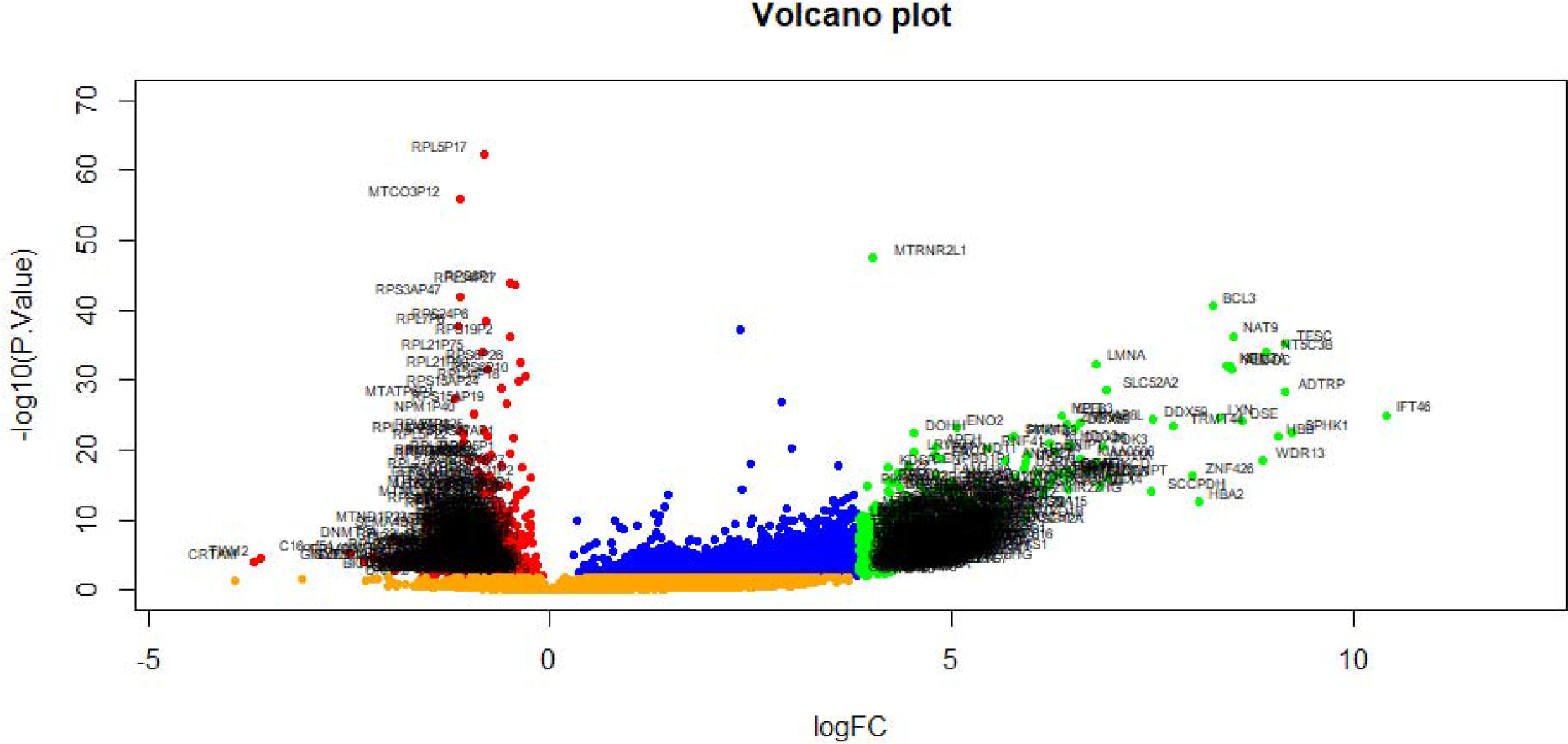
Volcano plot of differentially expressed genes. Genes with a significant change of more than two-fold were selected. Green dot represented up regulated significant genes and red dot represented down regulated significant genes.

**Fig. 2.**
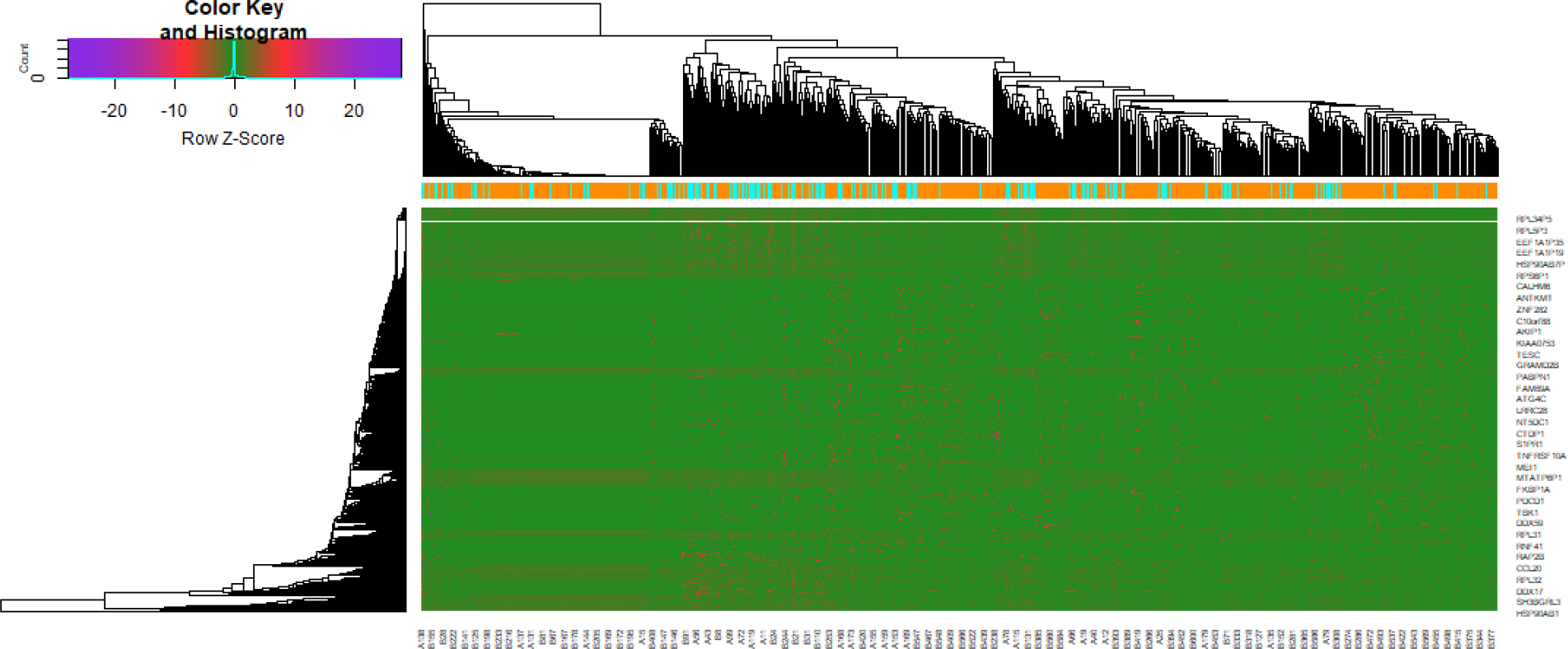
Heat map of differentially expressed genes. Legend on the top left indicate log fold change of genes. (A1 – A37 = normal control samples; B1 – B47 = HF samples)

**Table 1.**
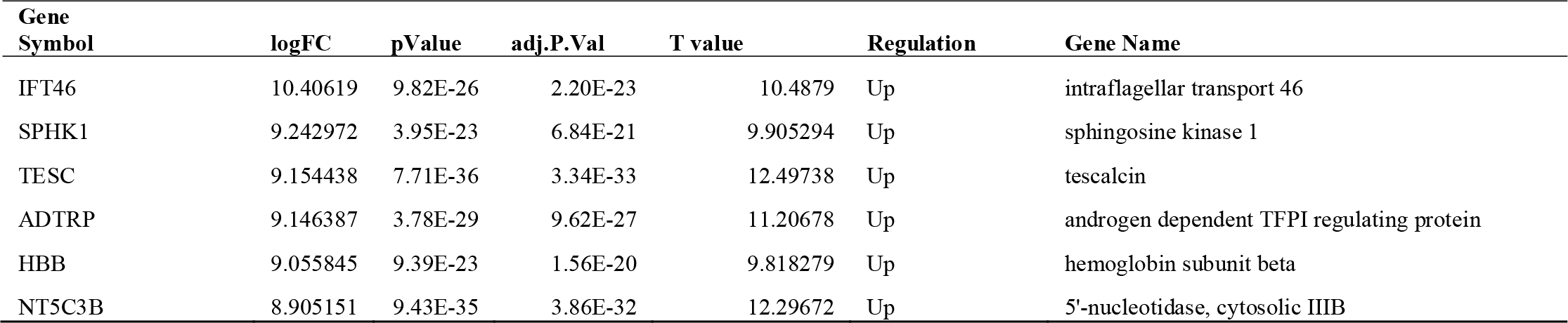
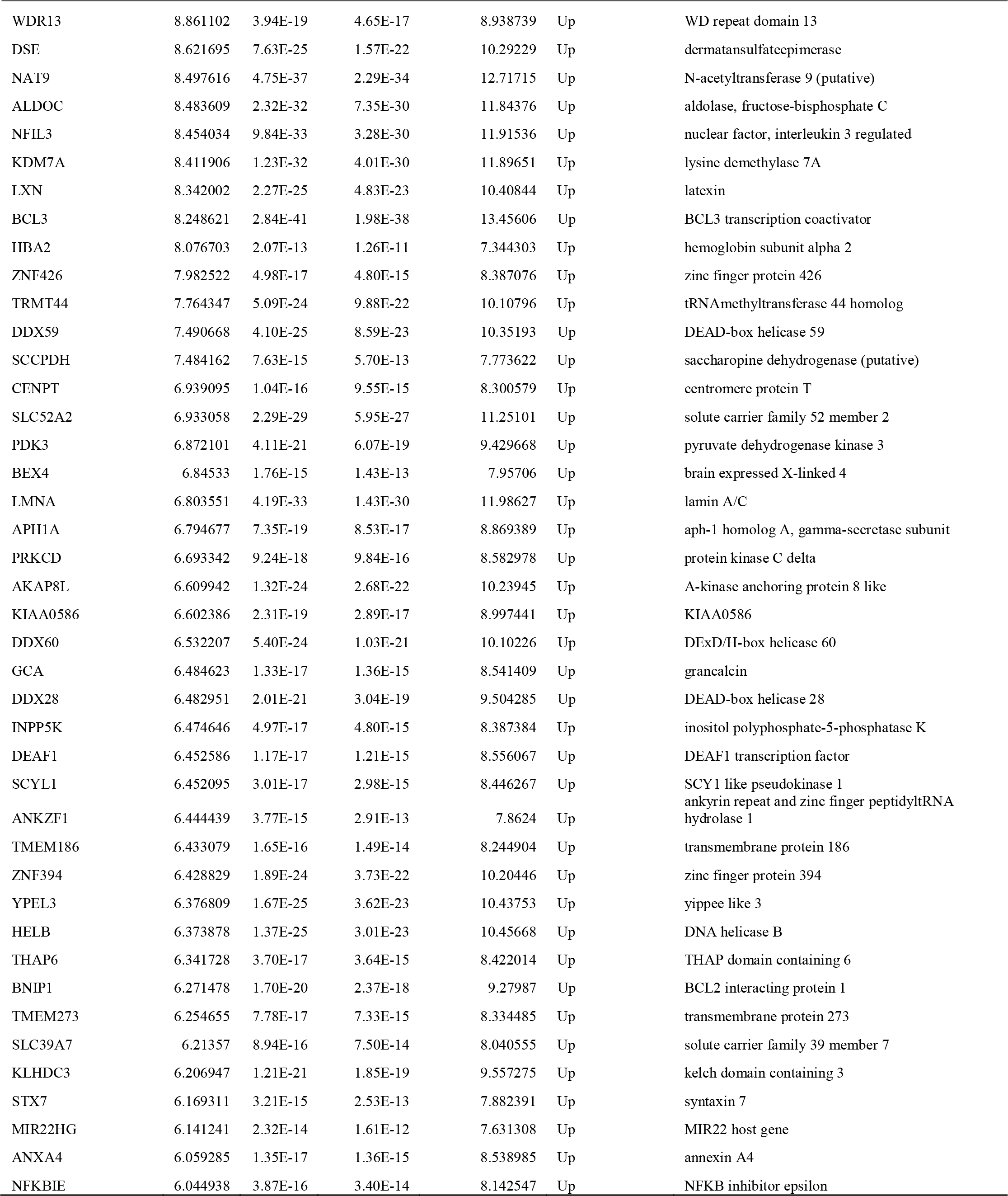
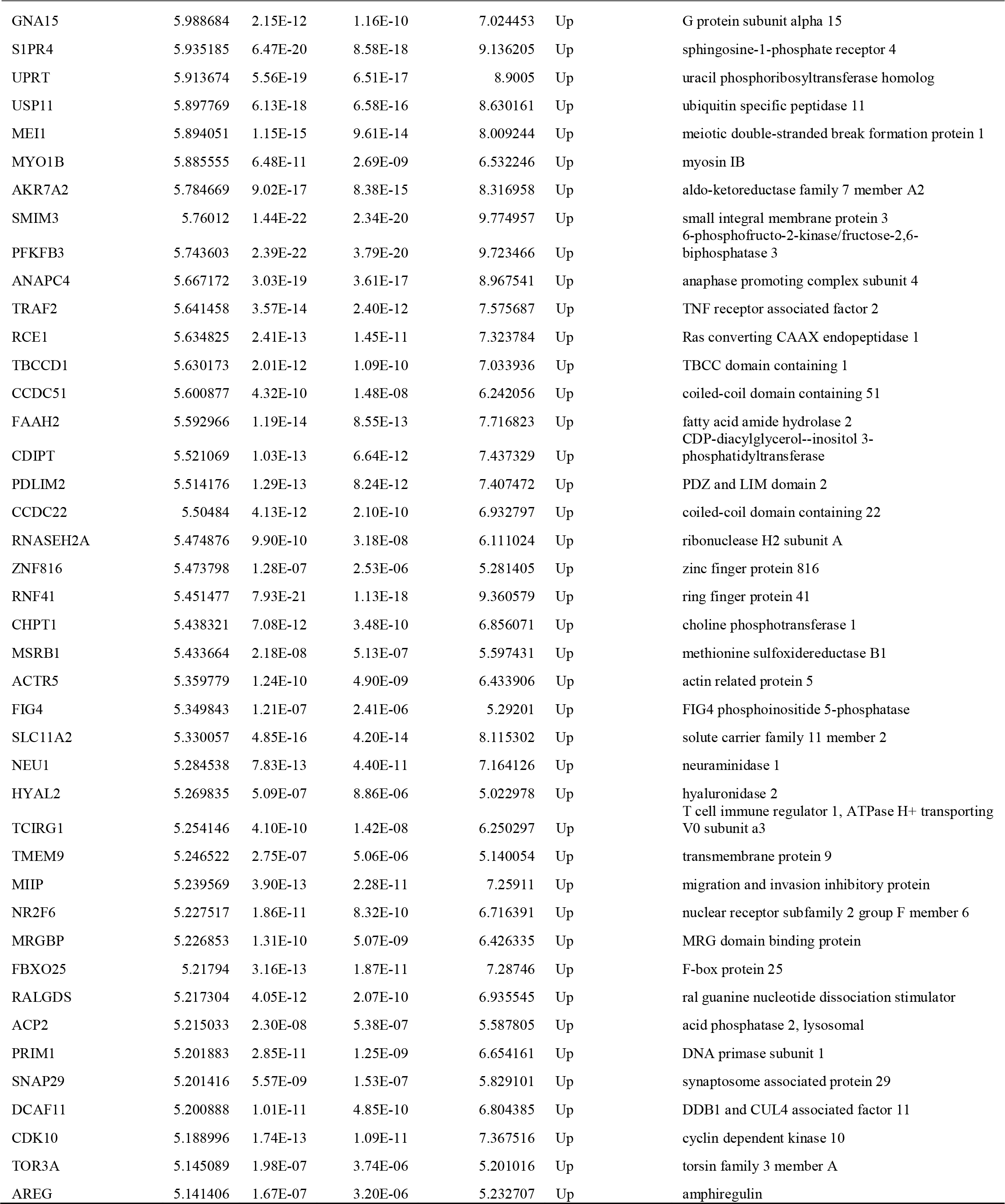
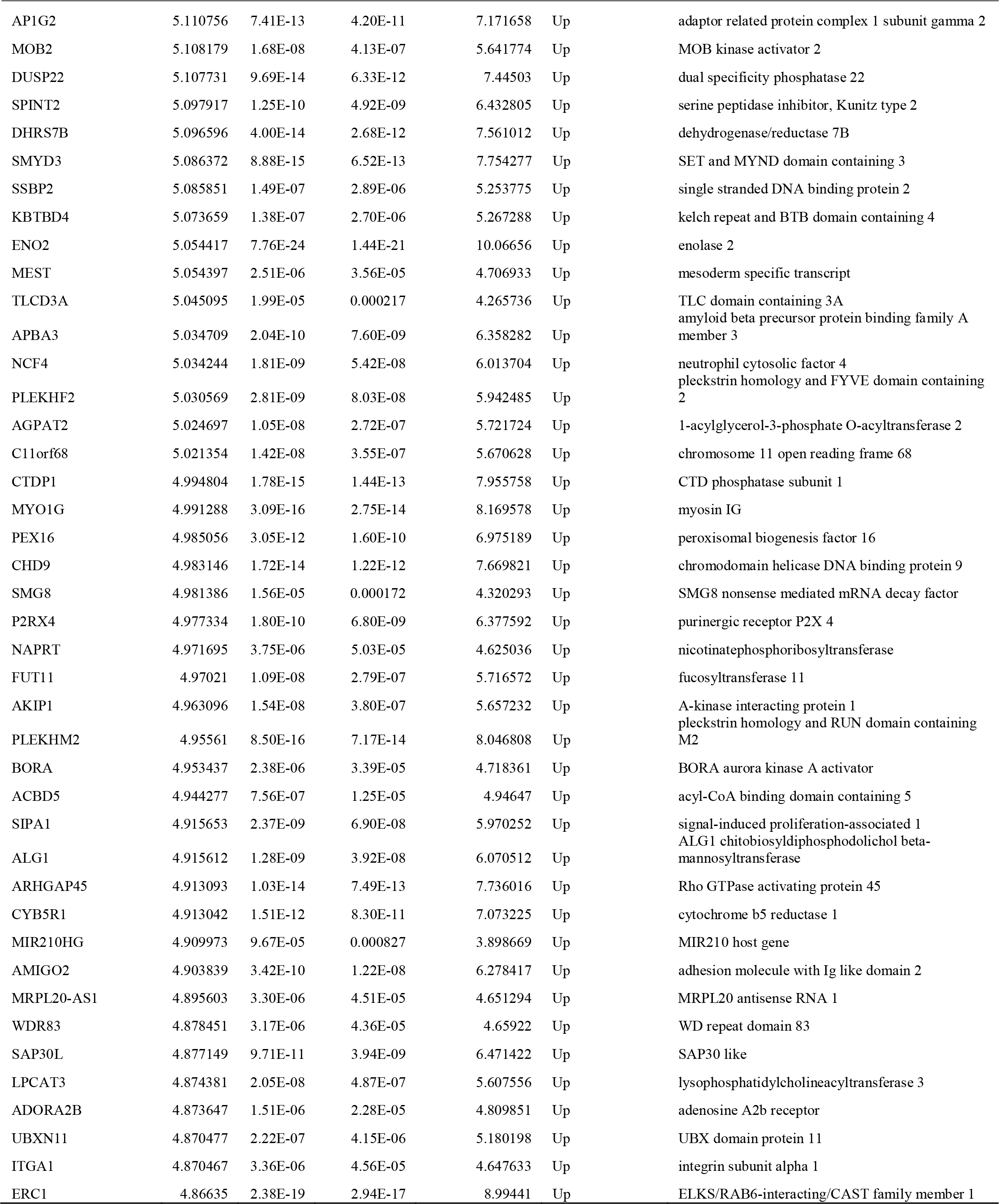
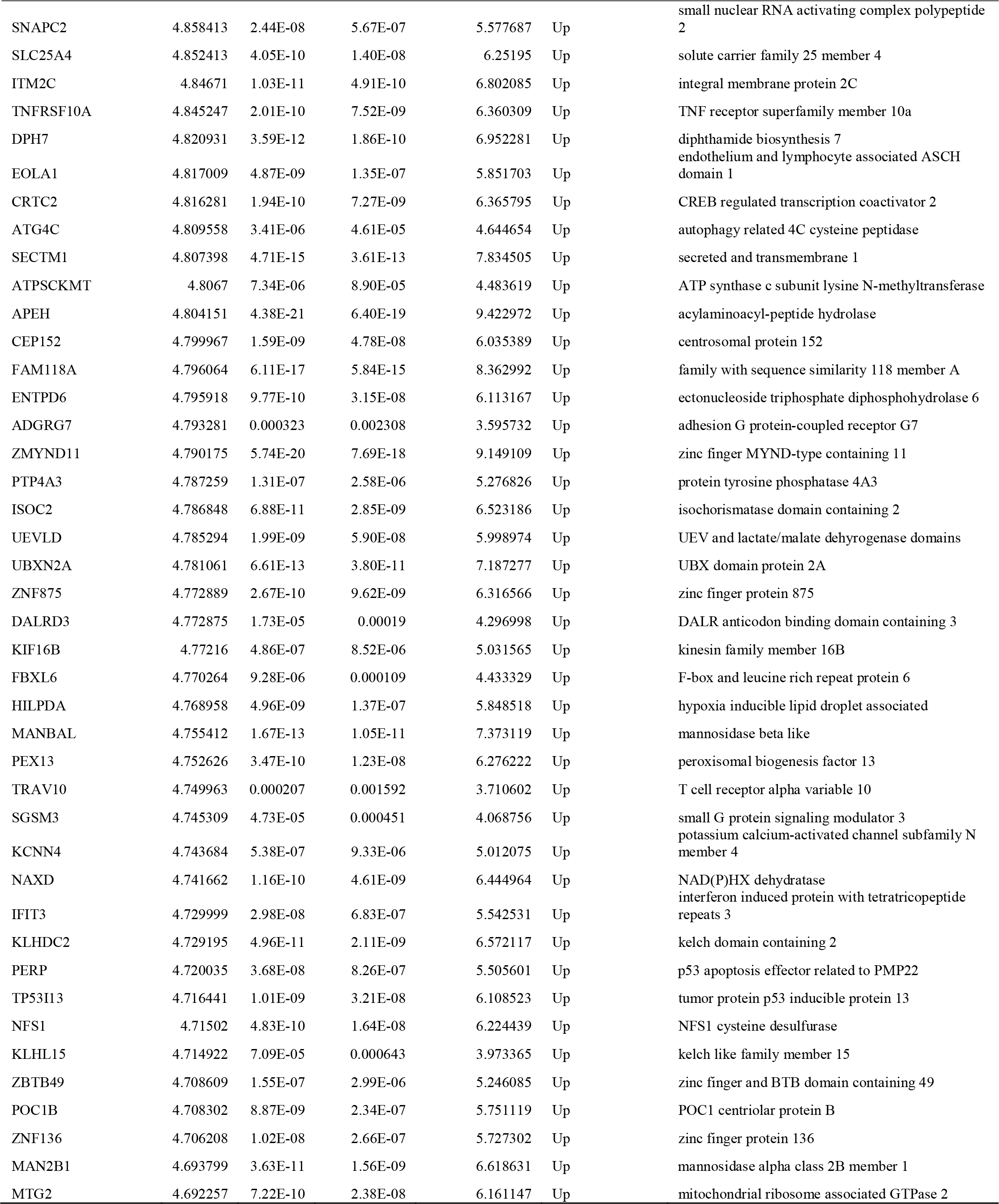
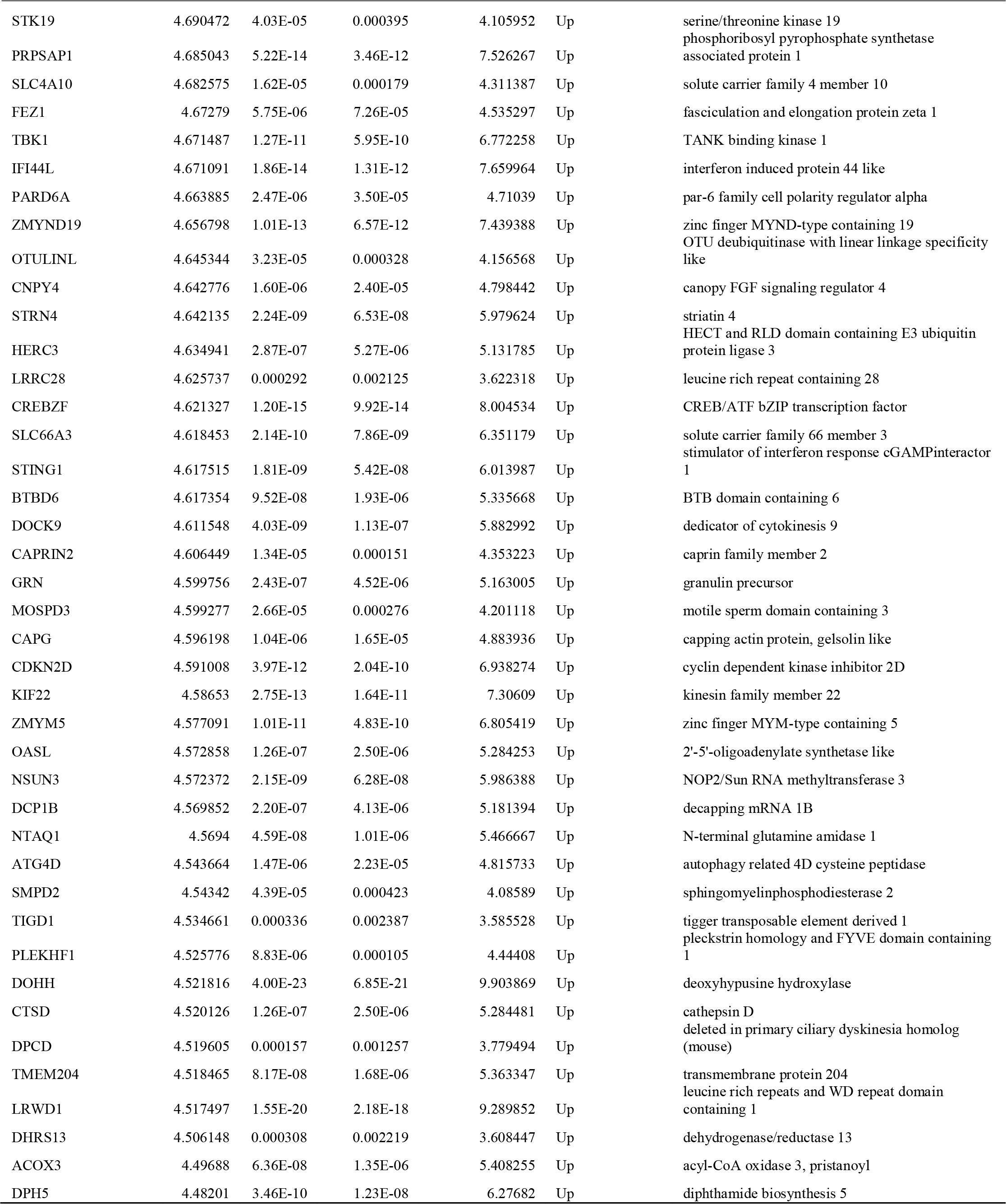
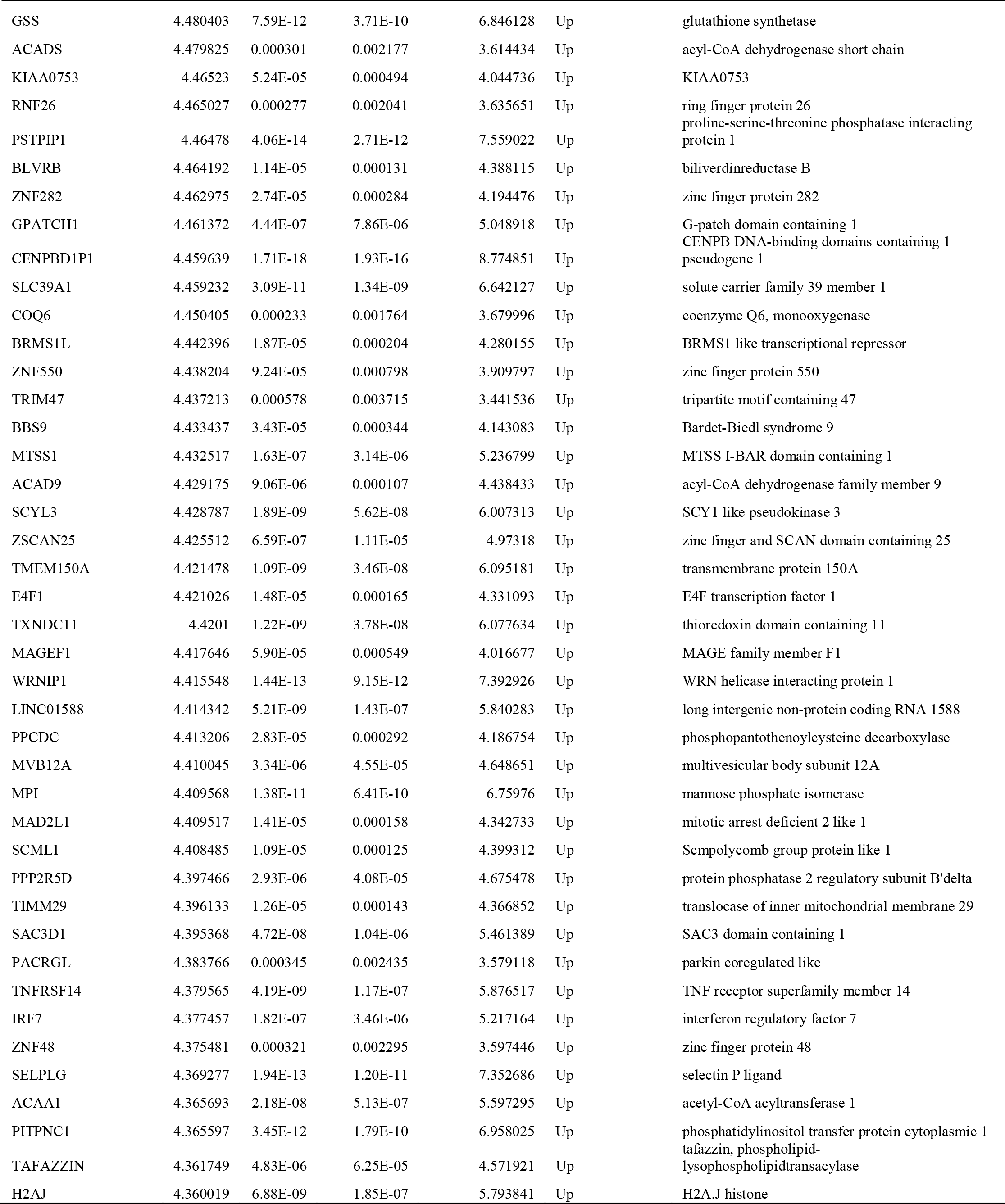
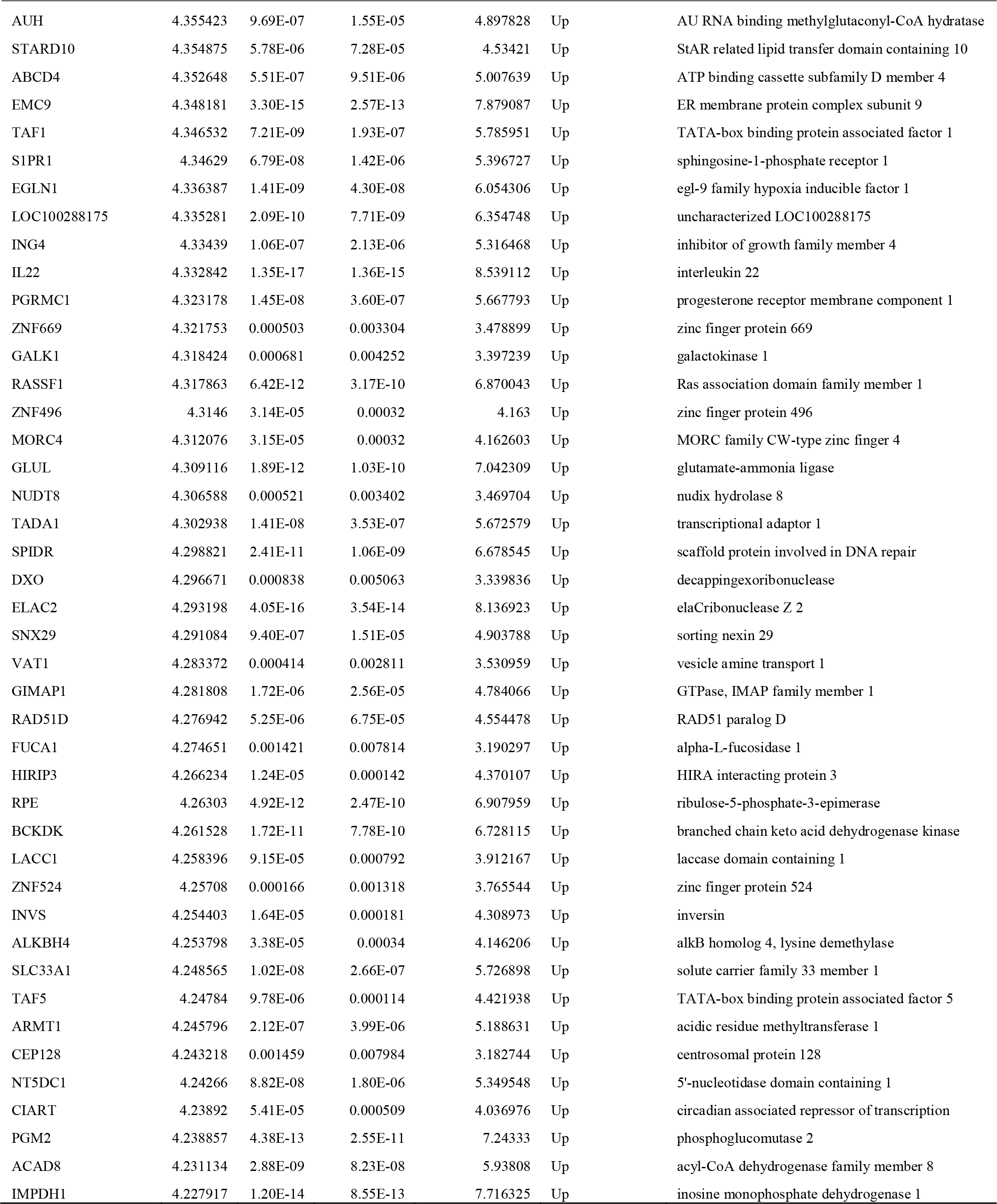
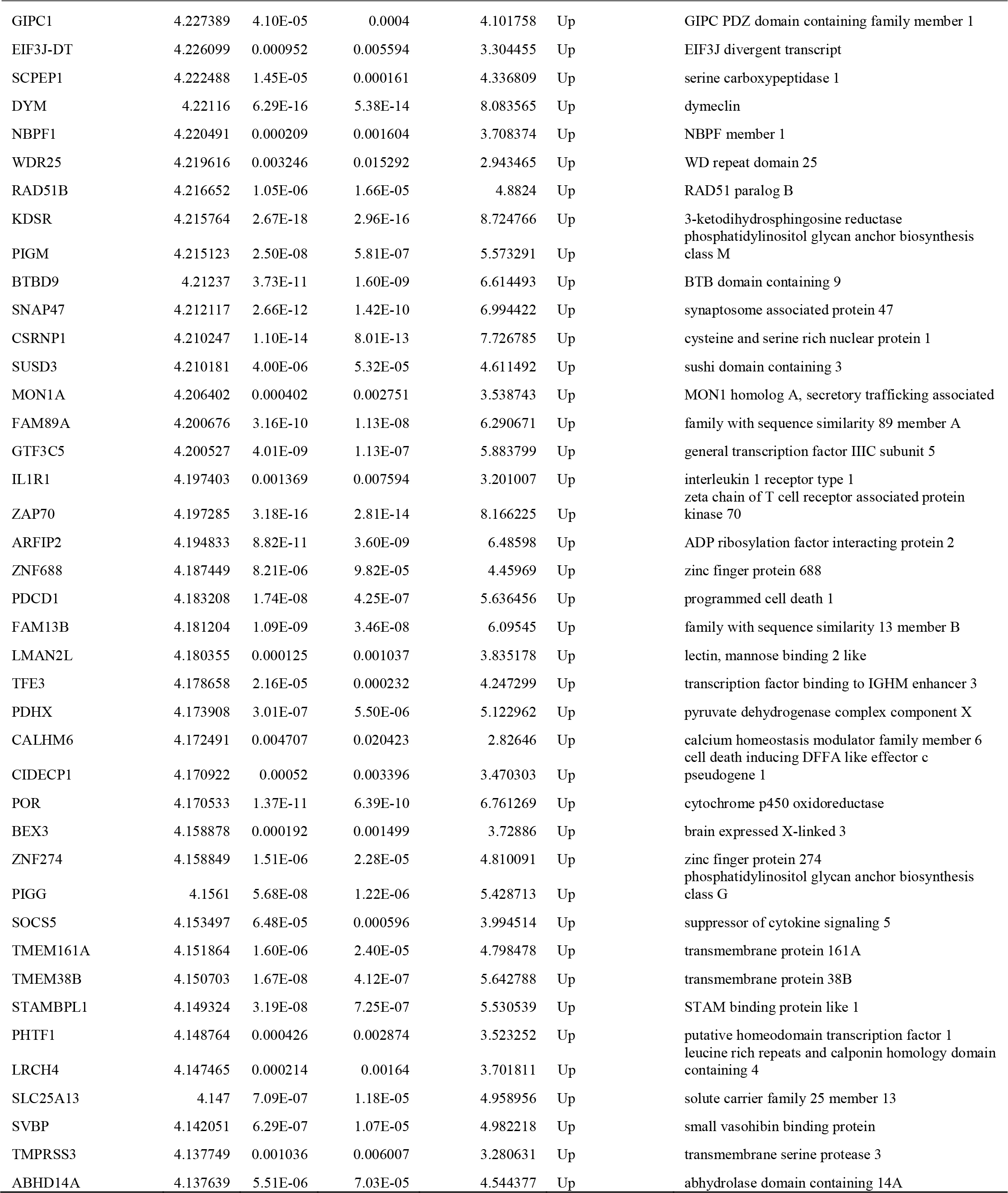
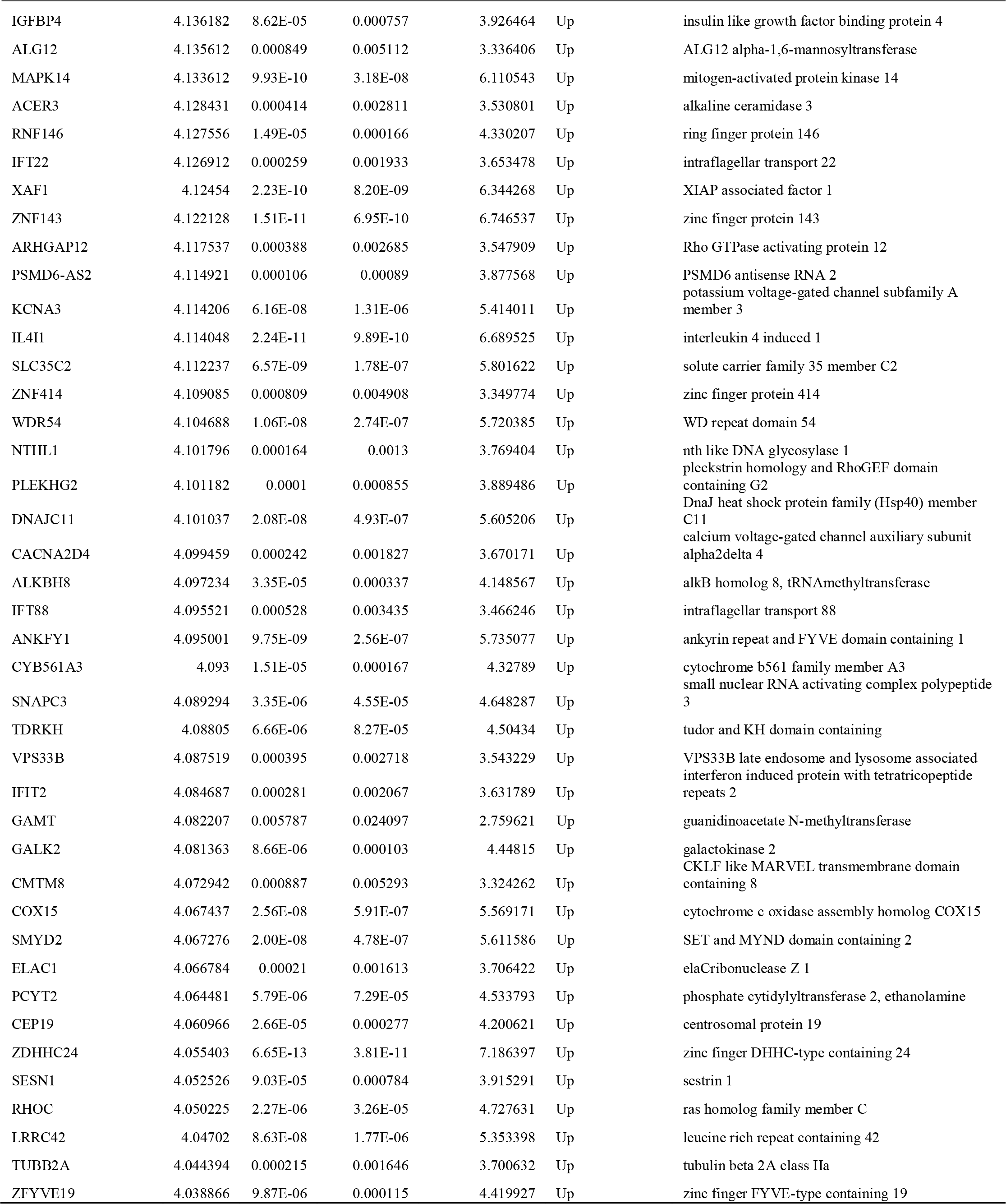
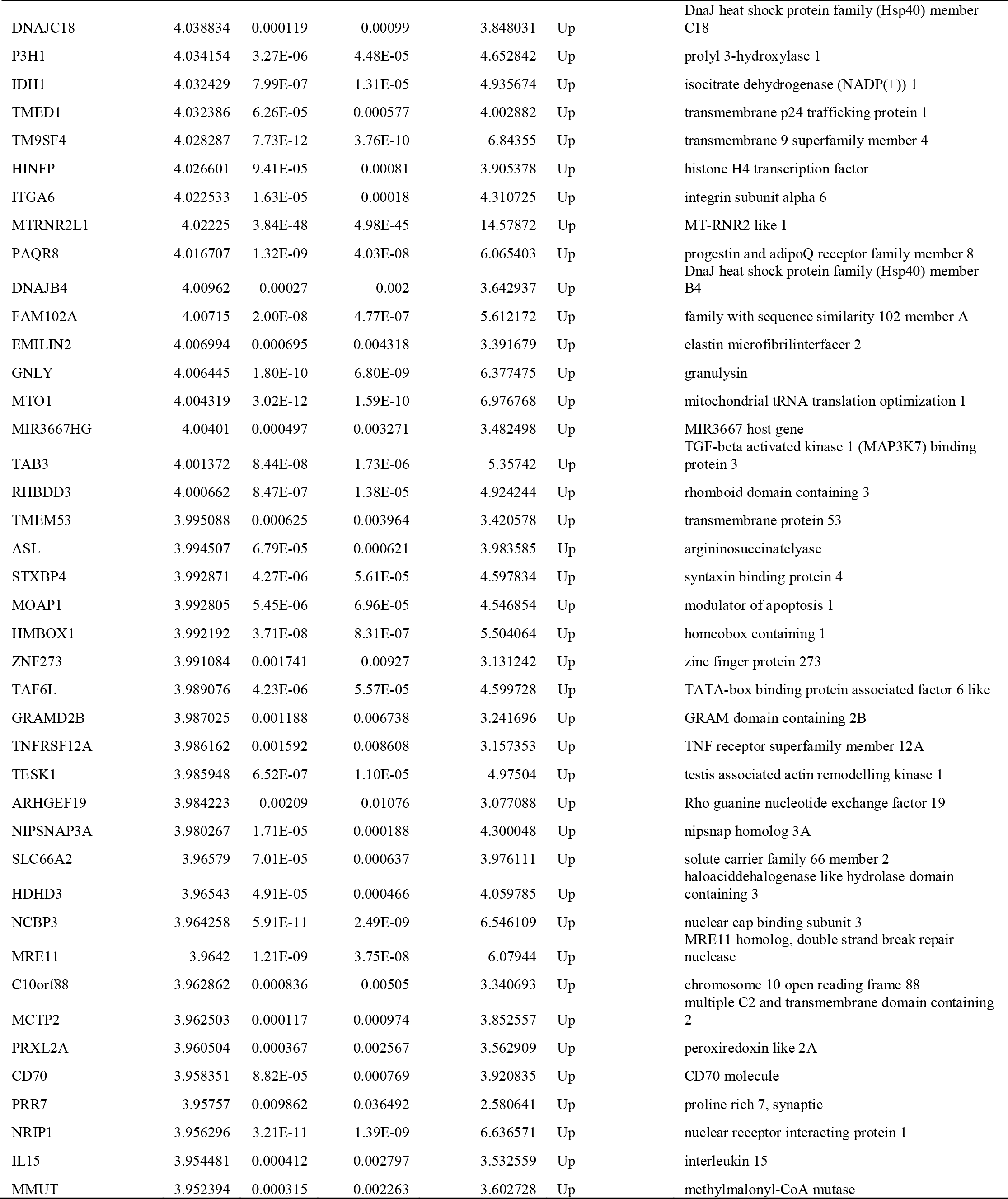
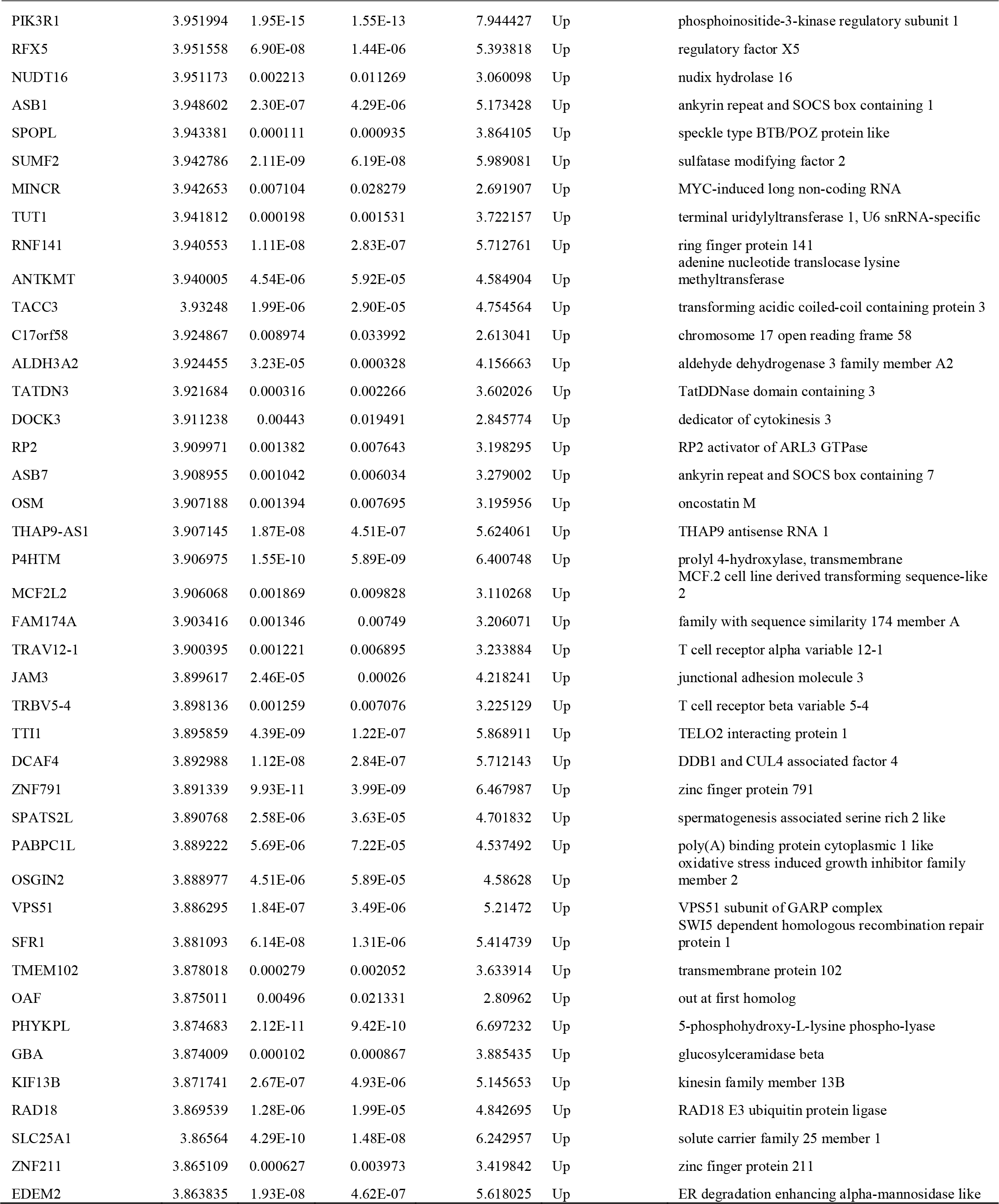
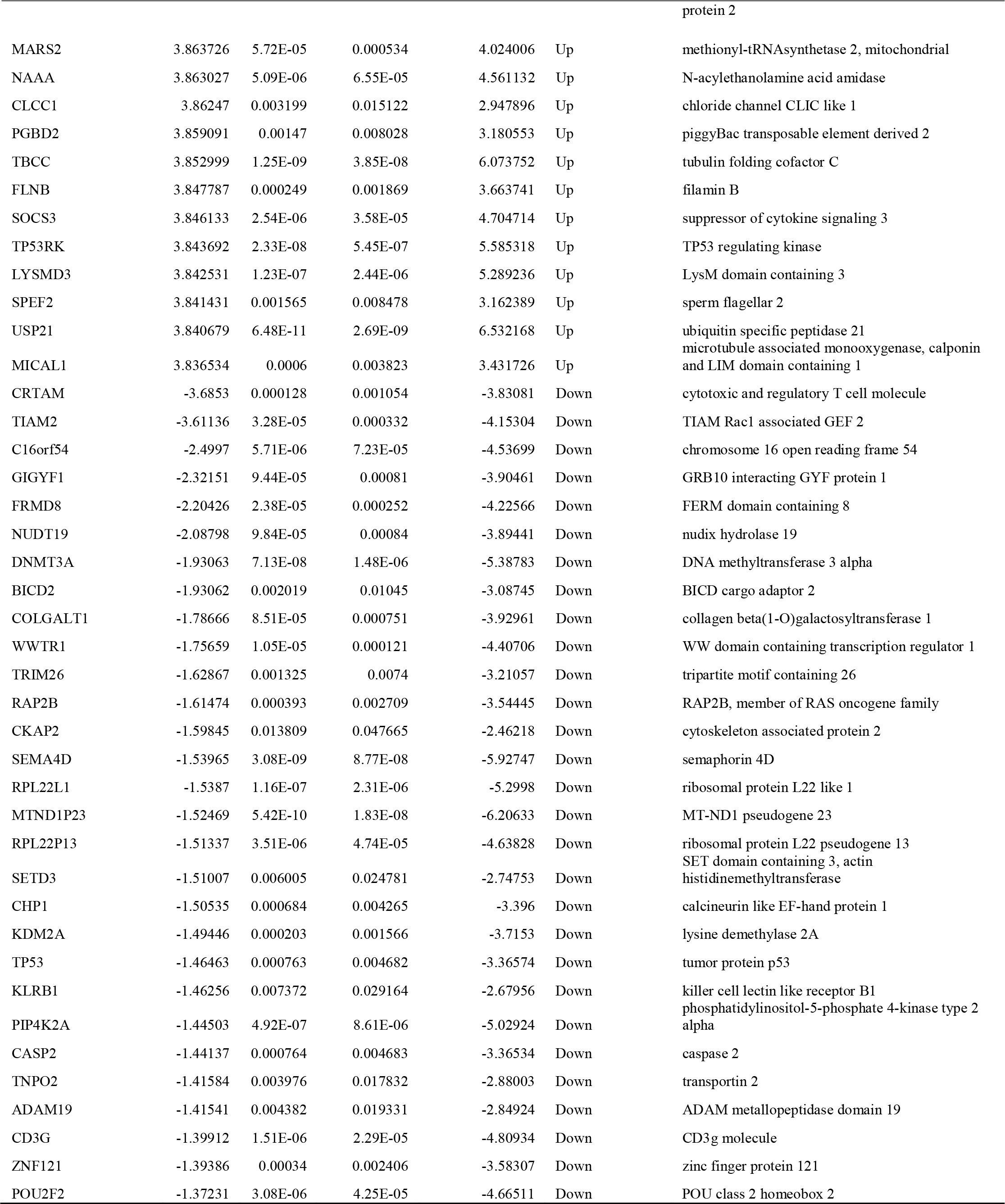
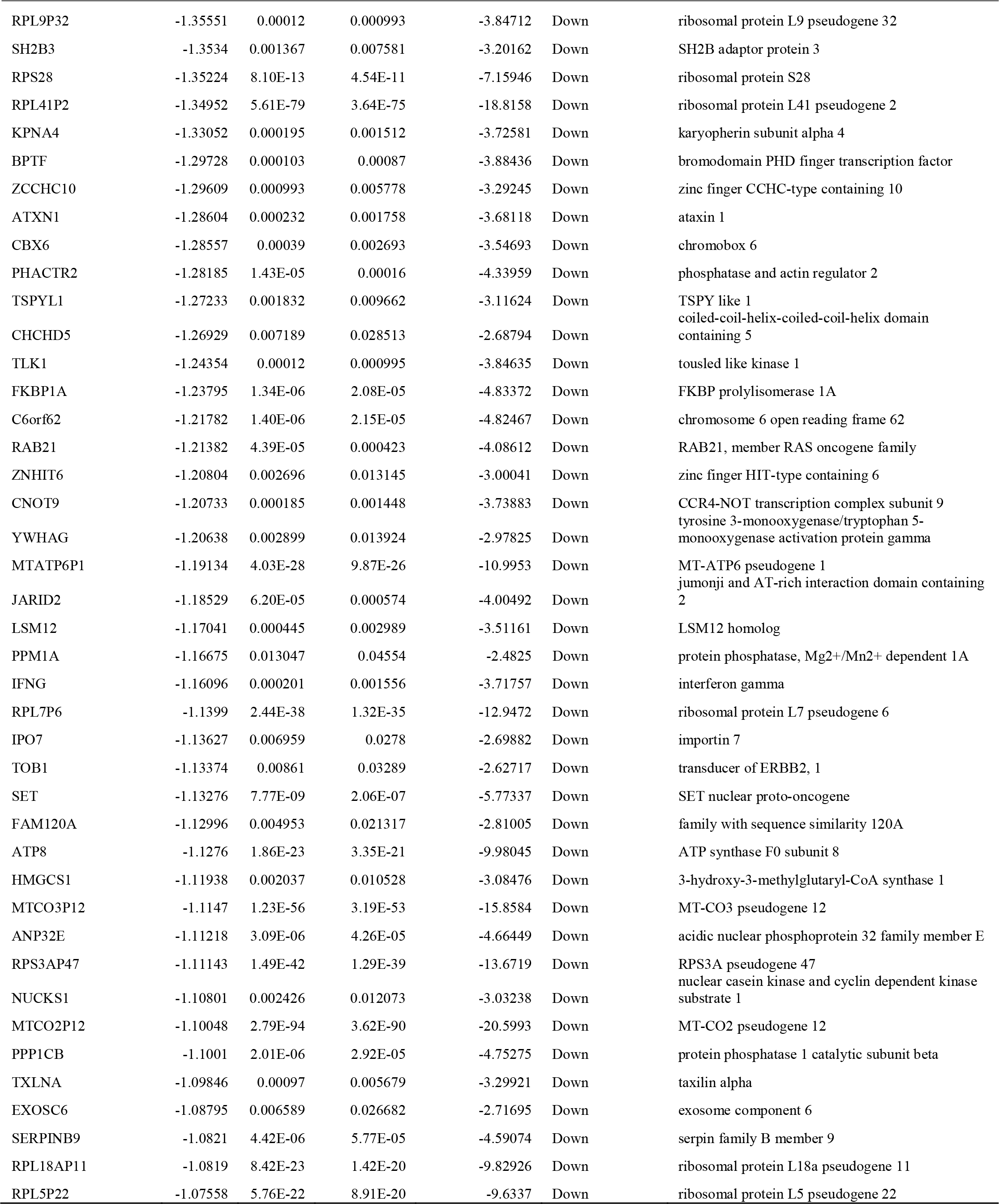
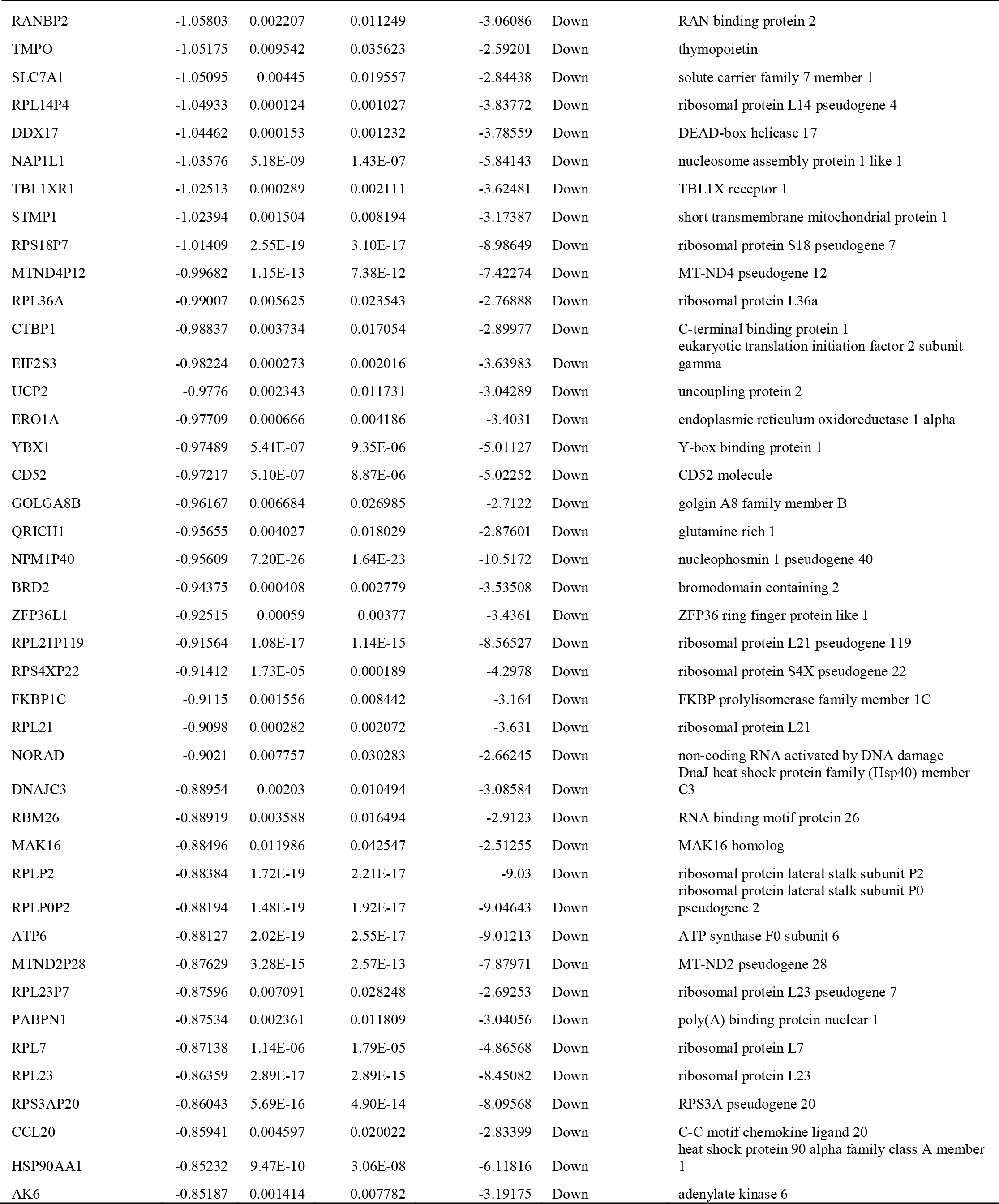
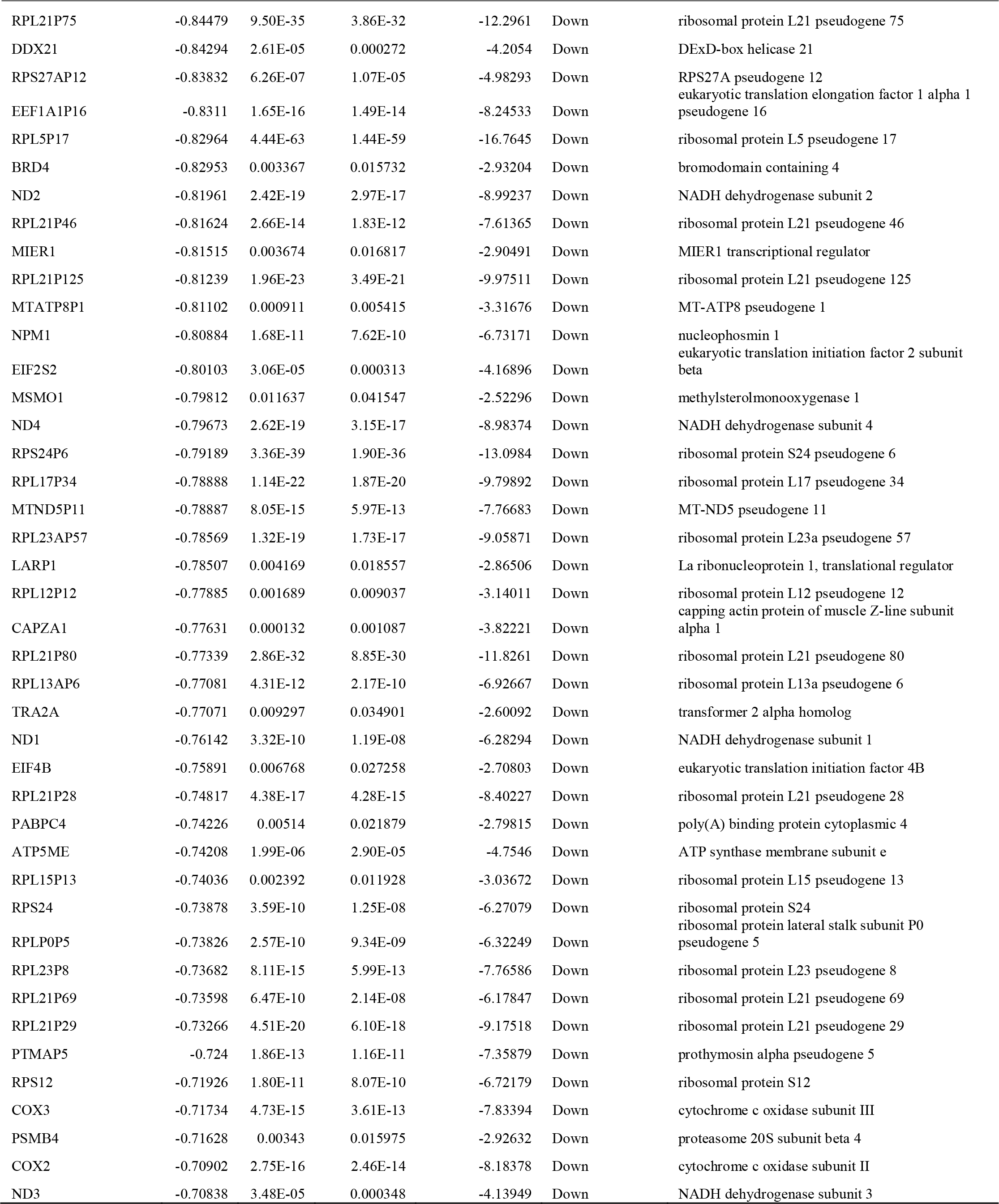
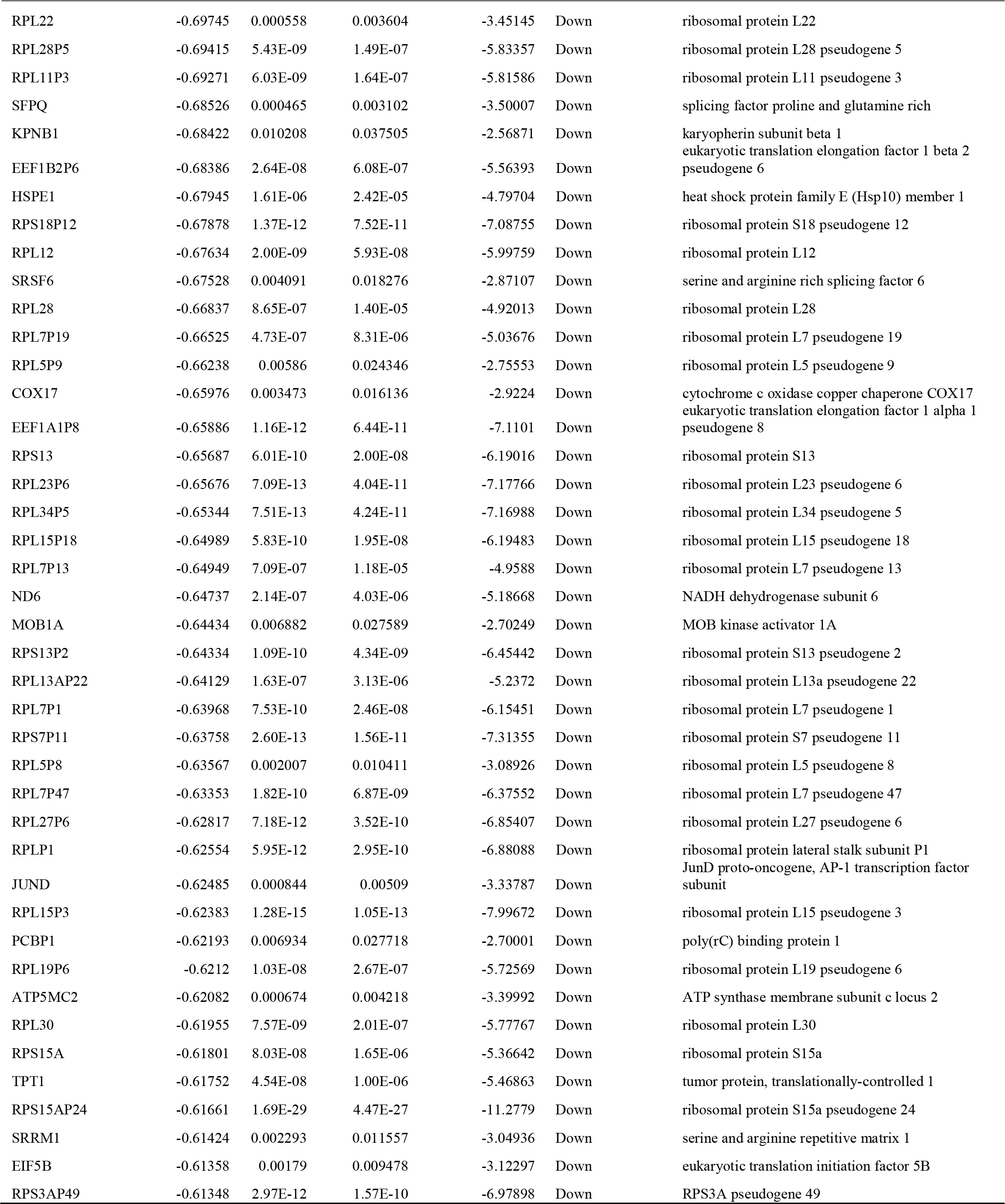
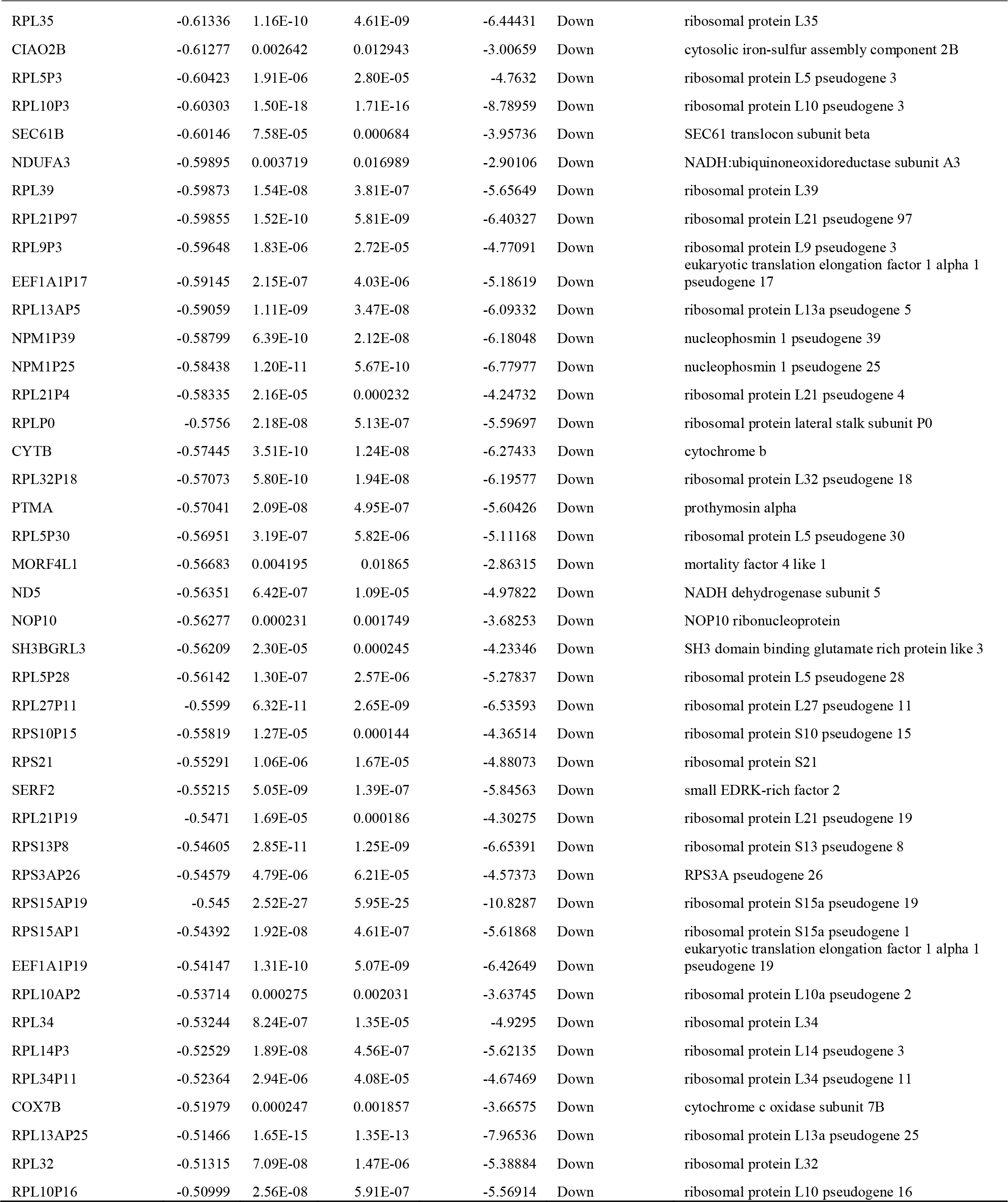
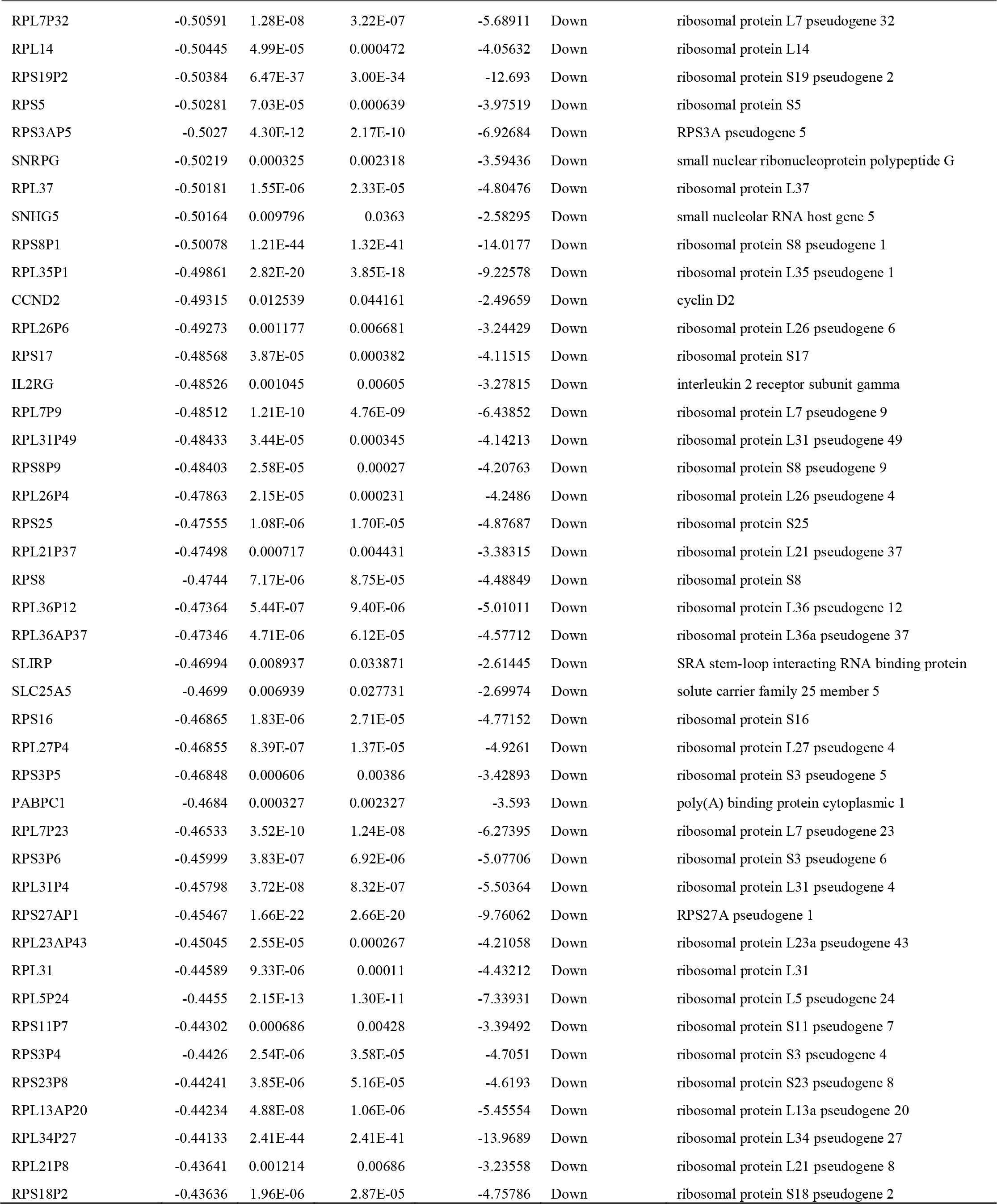
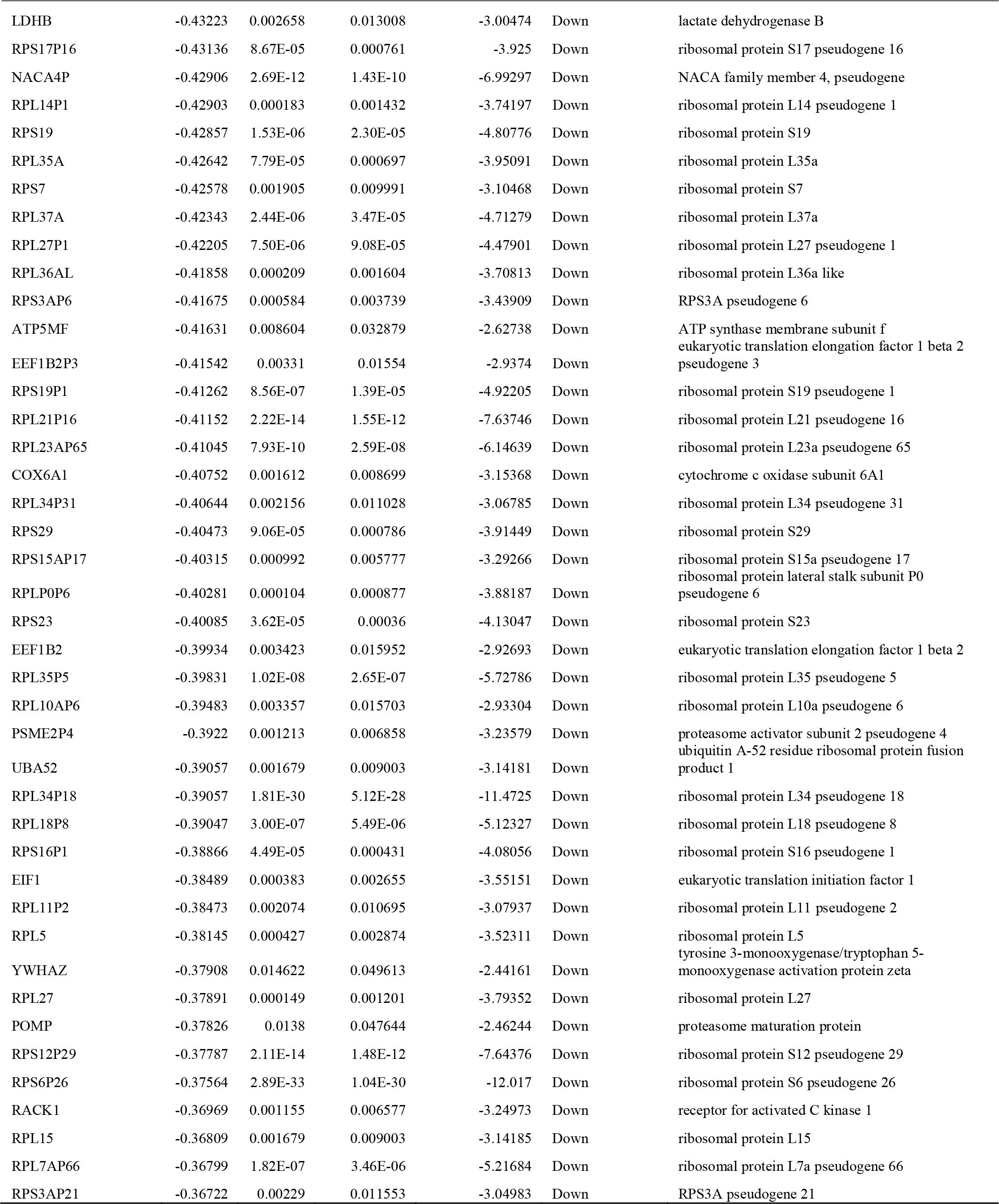
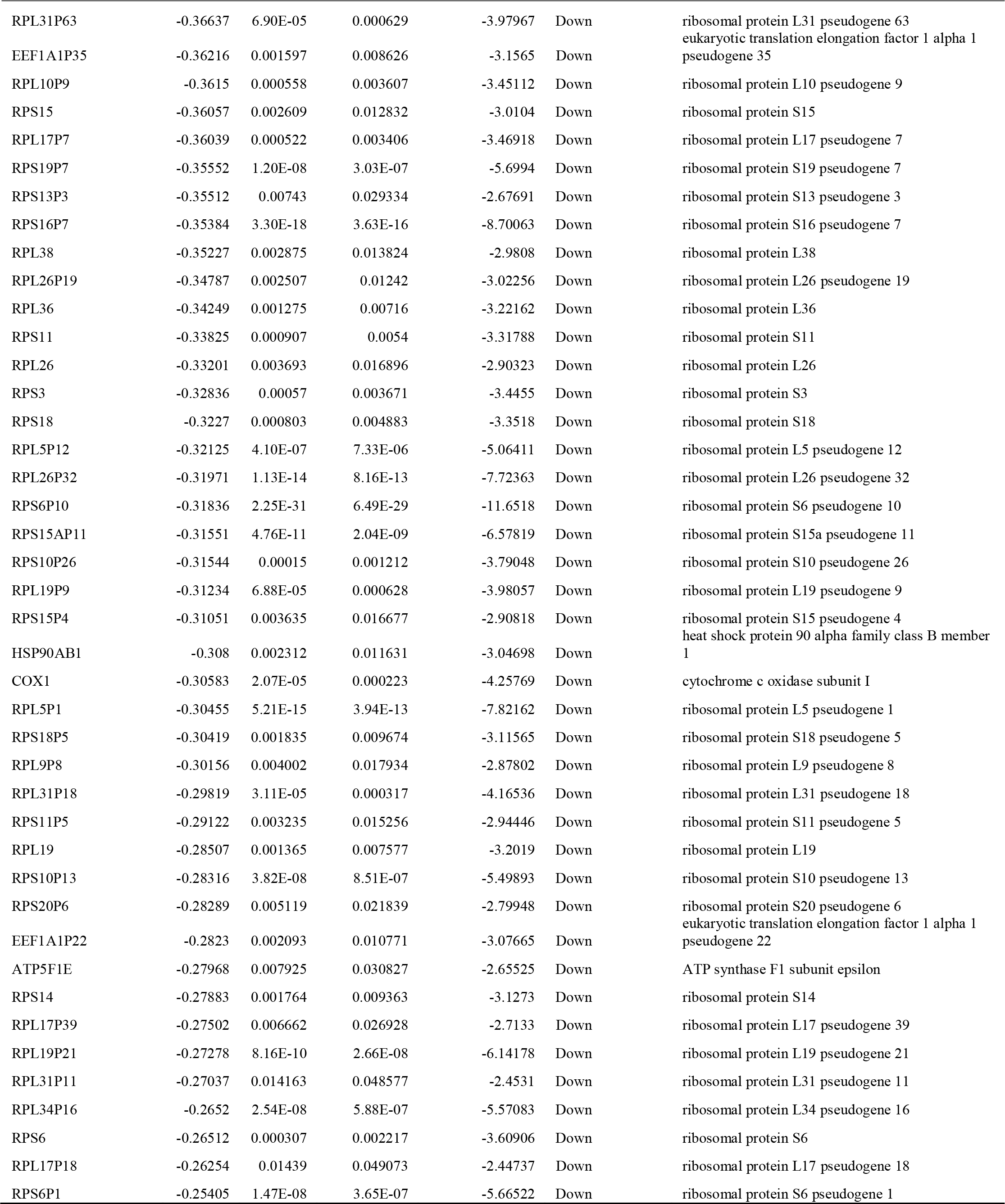
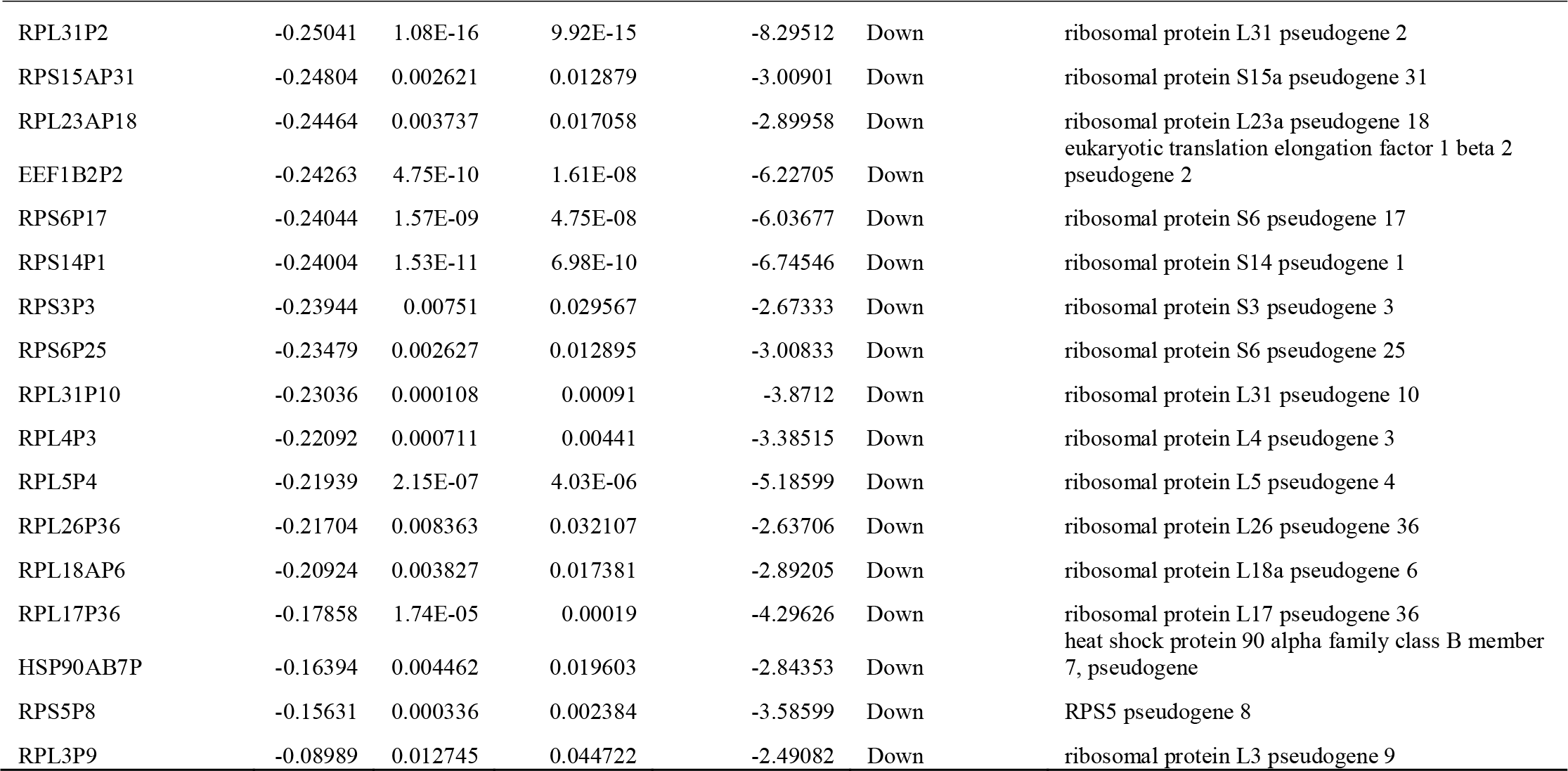
The statistical metrics for key differentially expressed genes (DEGs)

### GO and pathway enrichment analyses of DEGs

All 860 DEGs were analyzed by g:Profiler software and the results of GO enrichment analysis indicated that 1) for BP, up regulated genes were enriched in cellular metabolic process and metabolic process and down regulated genes enriched in cellular nitrogen compound biosynthetic process and gene expression; 2) for CC up genes were enriched in intracellular anatomical structure and organelle and down regulated genes enriched in protein-containing complex and cytoplasm; 3) for MF up genes were enriched in catalytic activity and ion binding and down regulated genes enriched in heterocyclic compound binding and protein binding (Table 2). As shown in Table 3, REACTOME pathway enrichment analysis found significantly enriched pathways. Up regulated genes and down regulated genes enriched in metabolism of carbohydrates, metabolism, metabolism of RNA and metabolism of proteins.

**Table 2.**
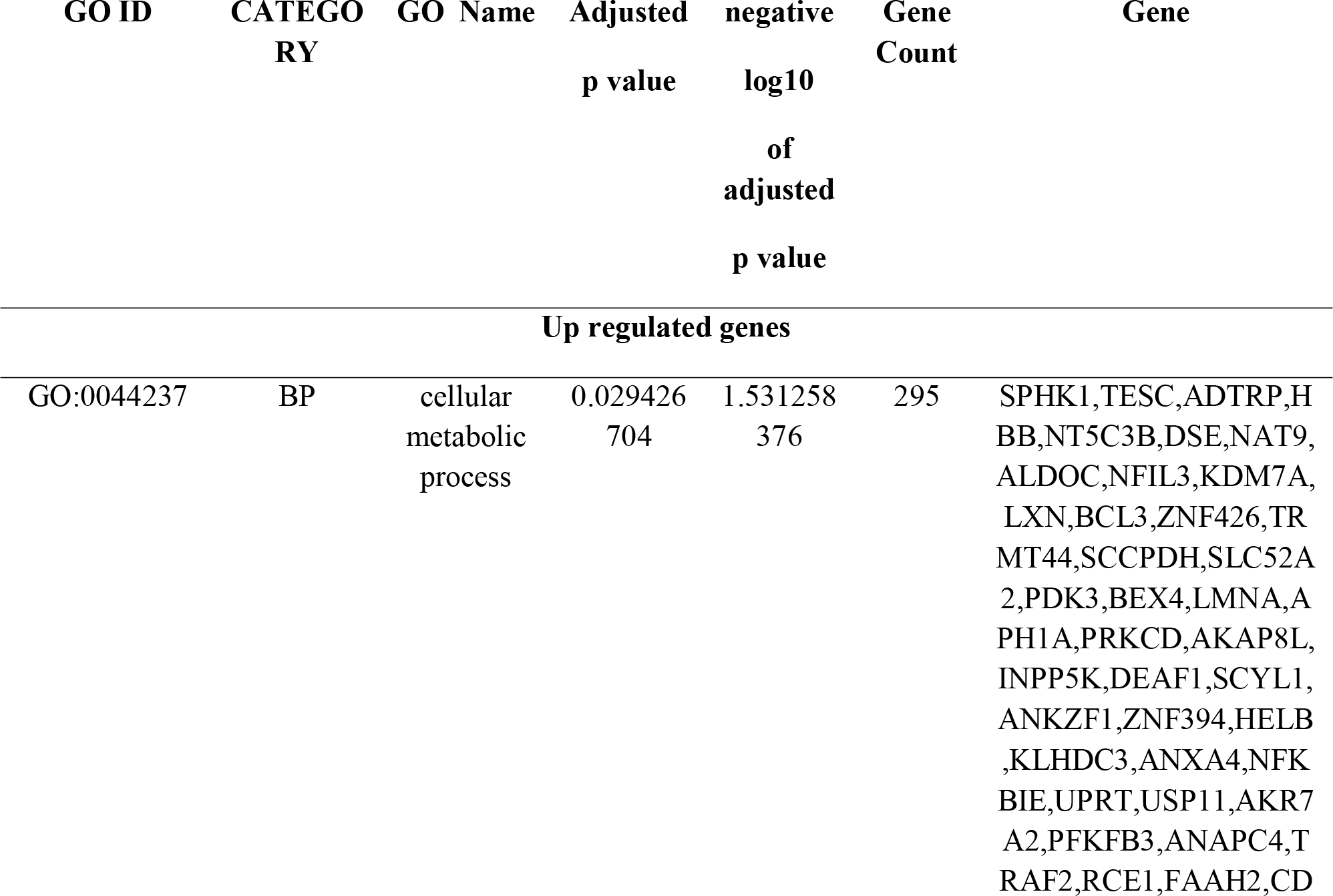
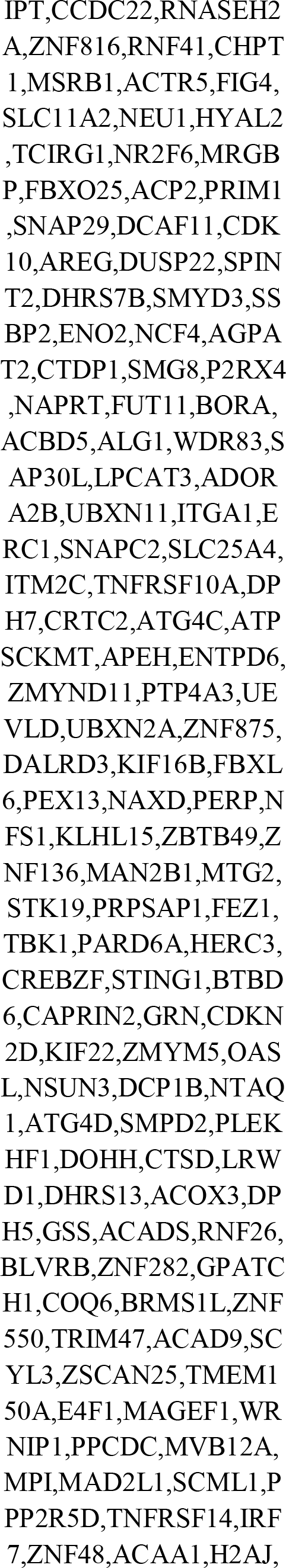
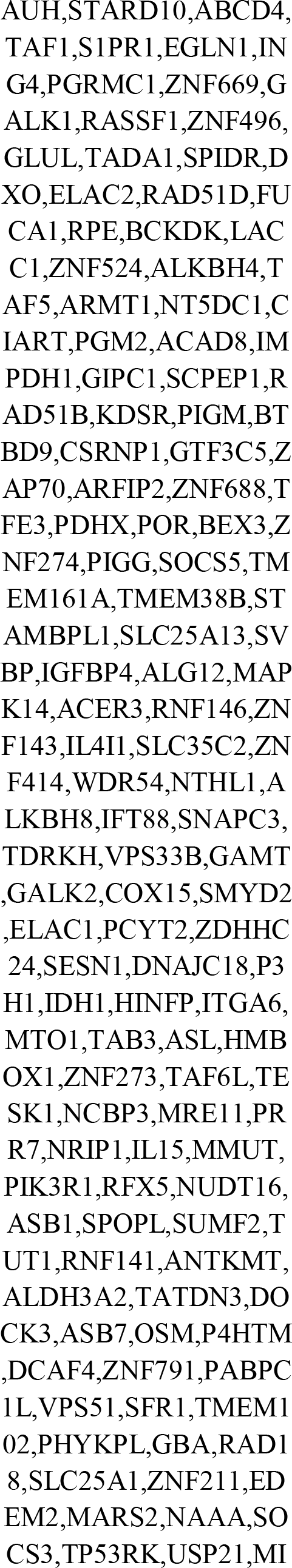
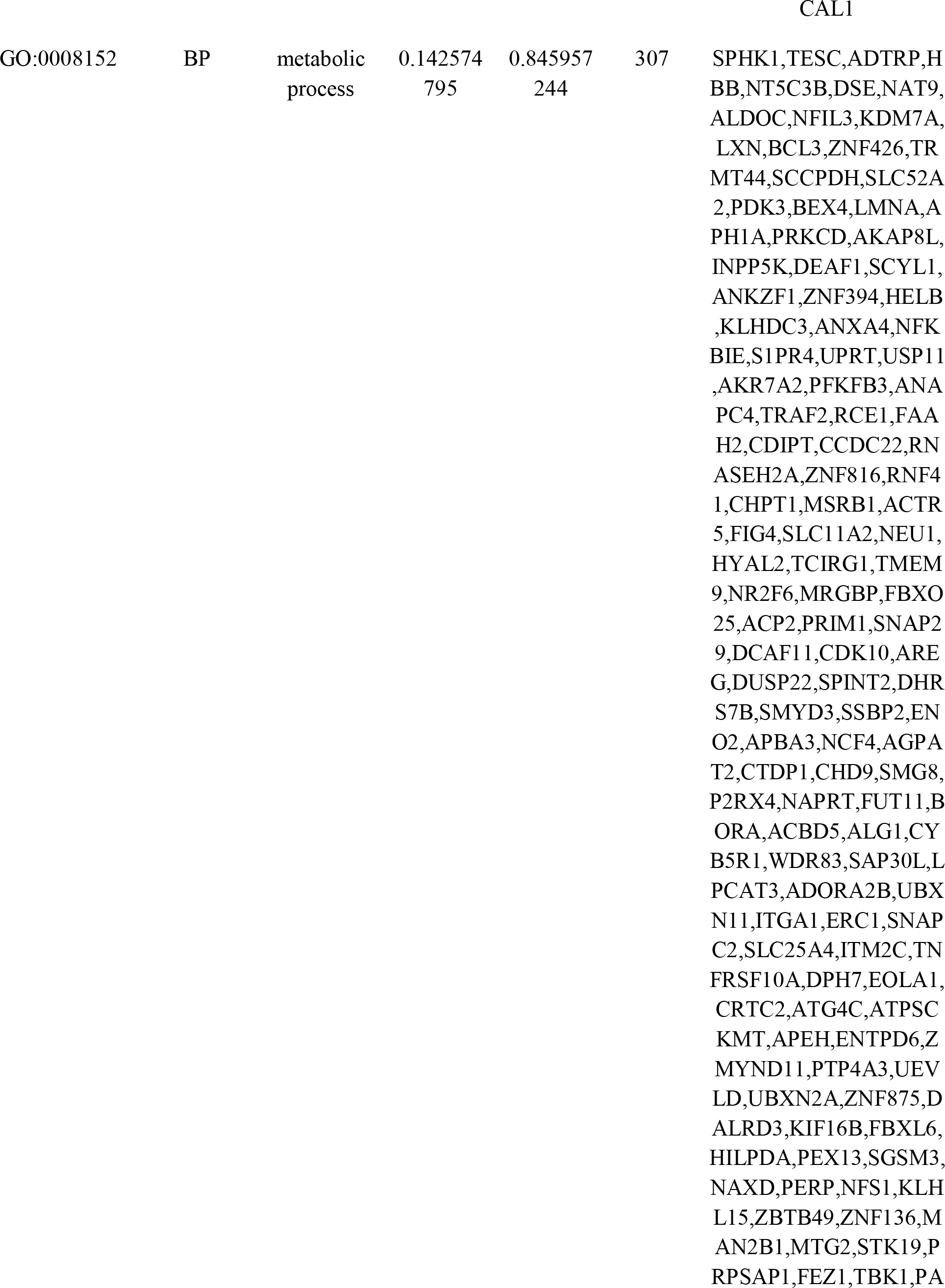
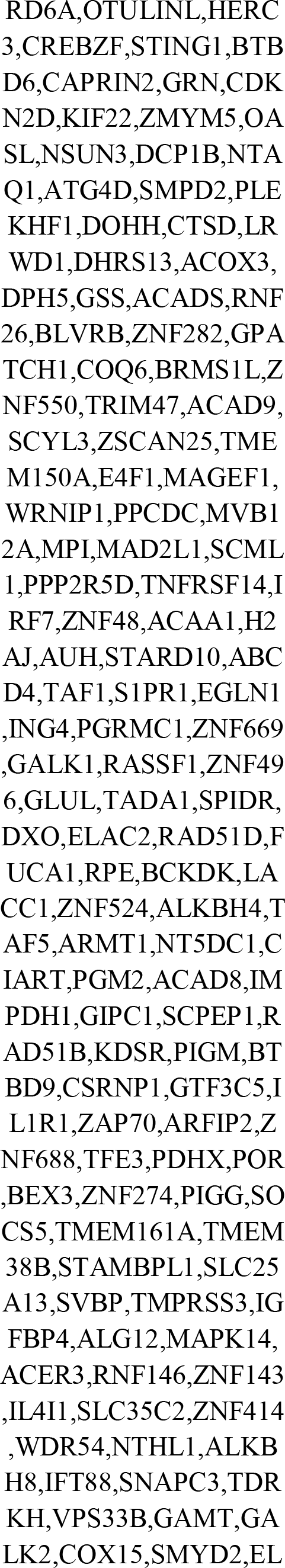
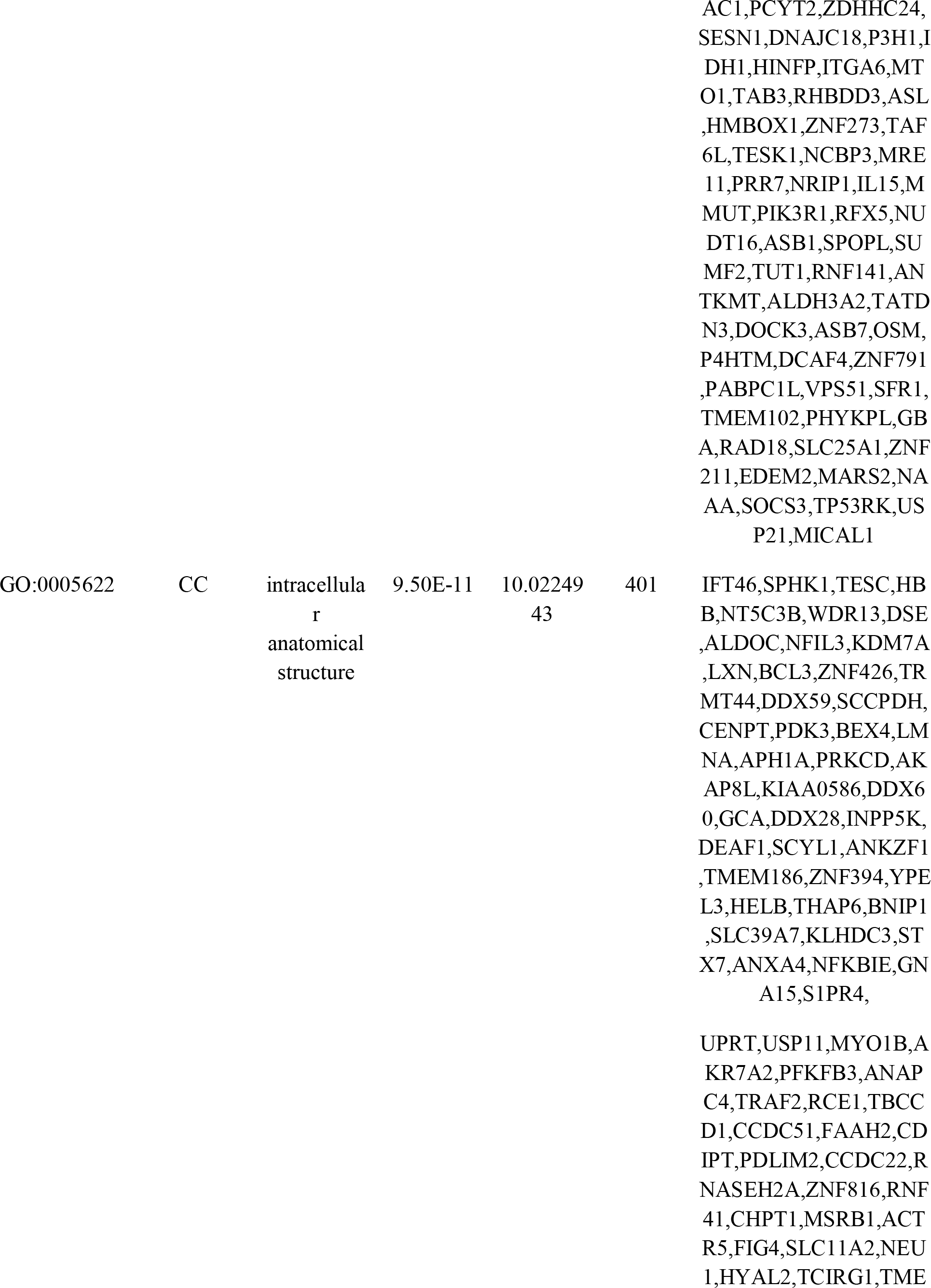
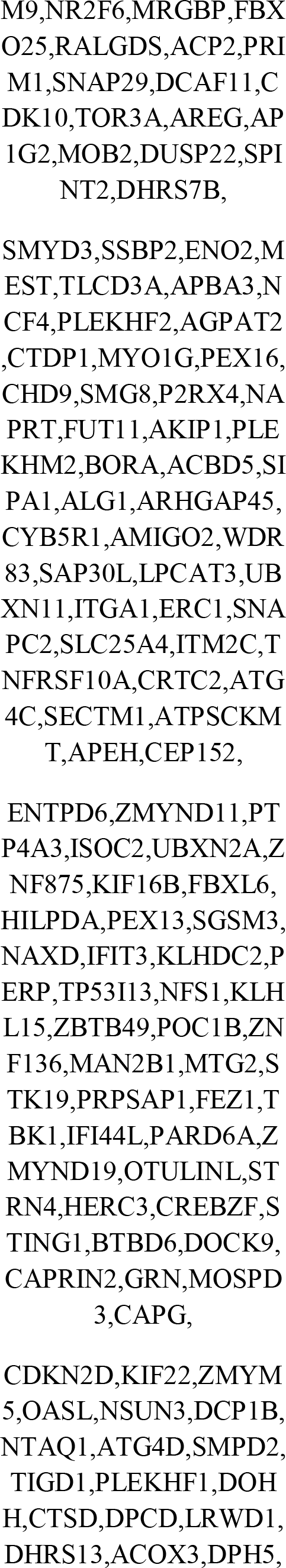
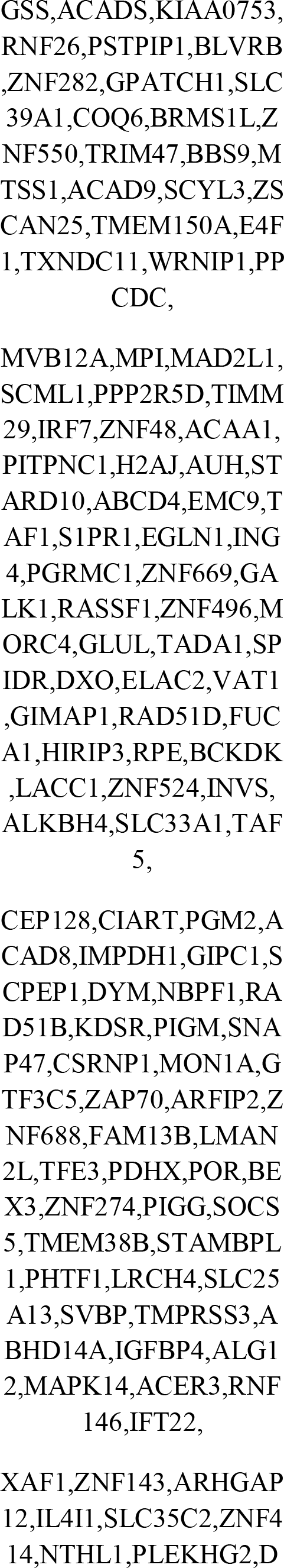
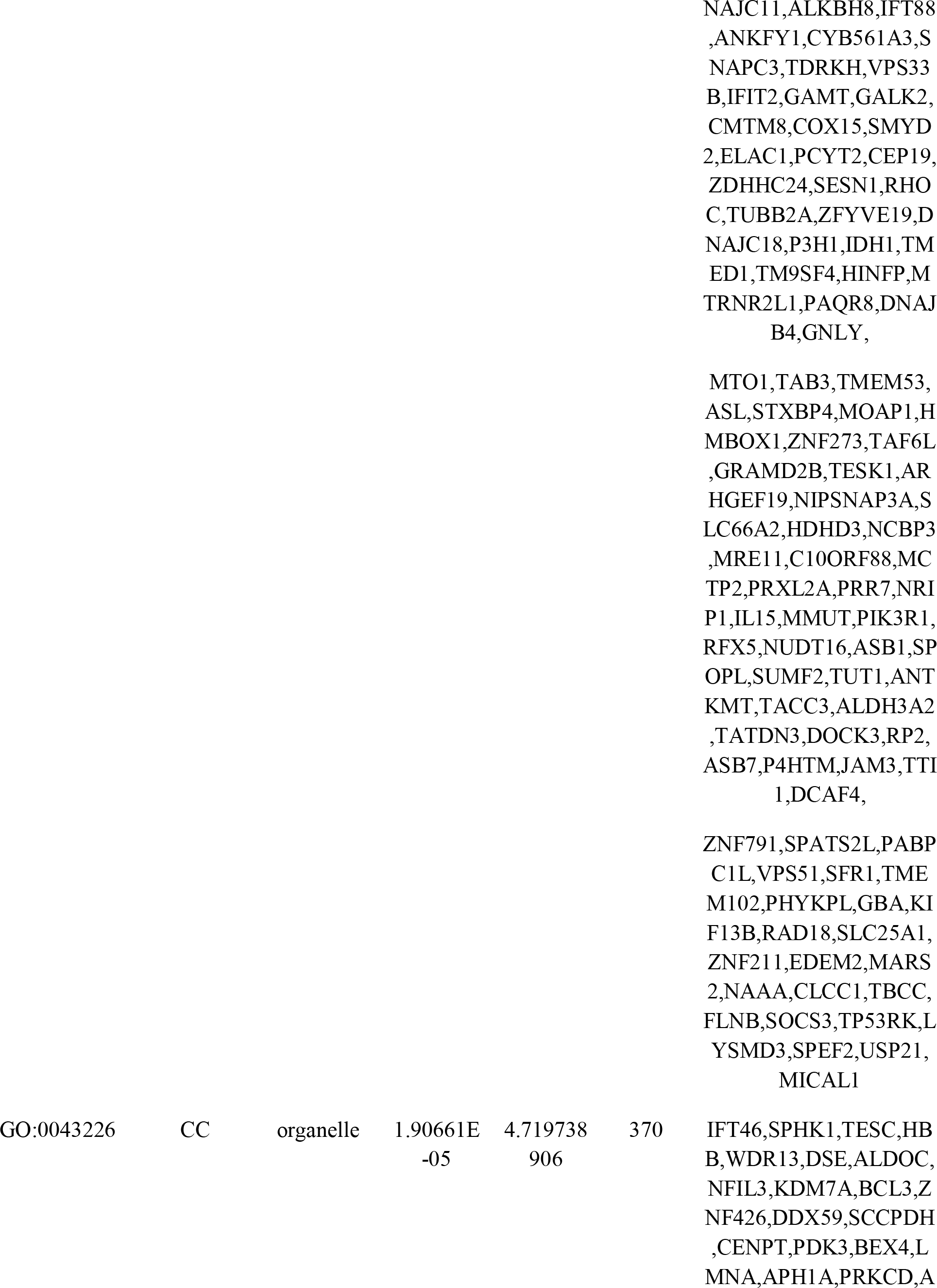
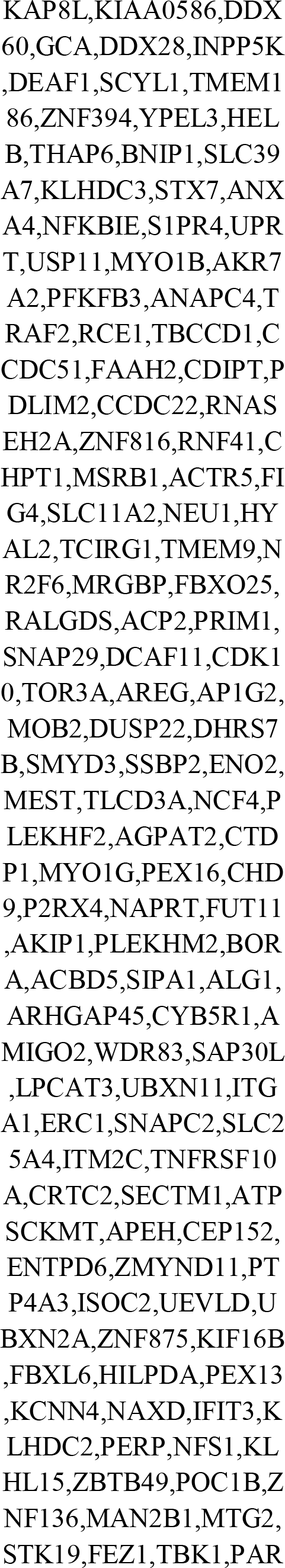
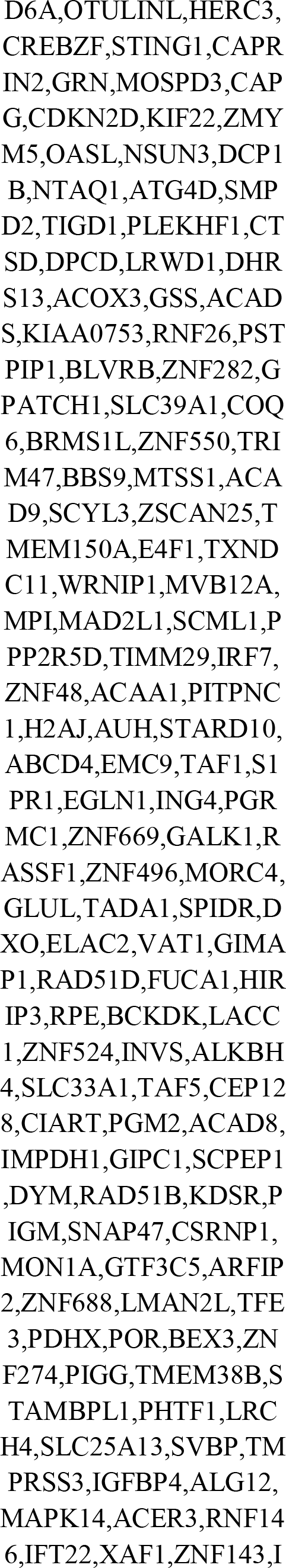
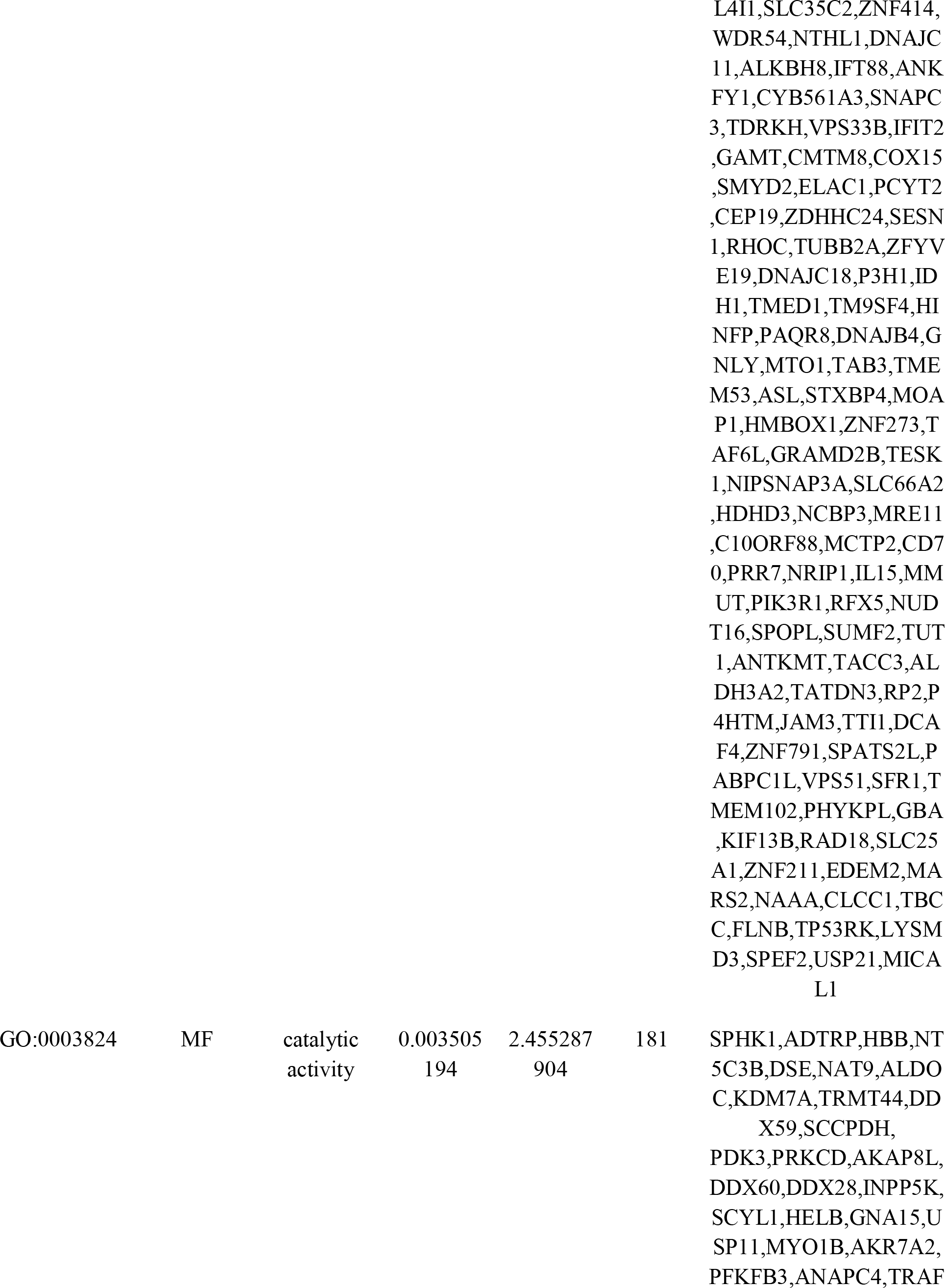
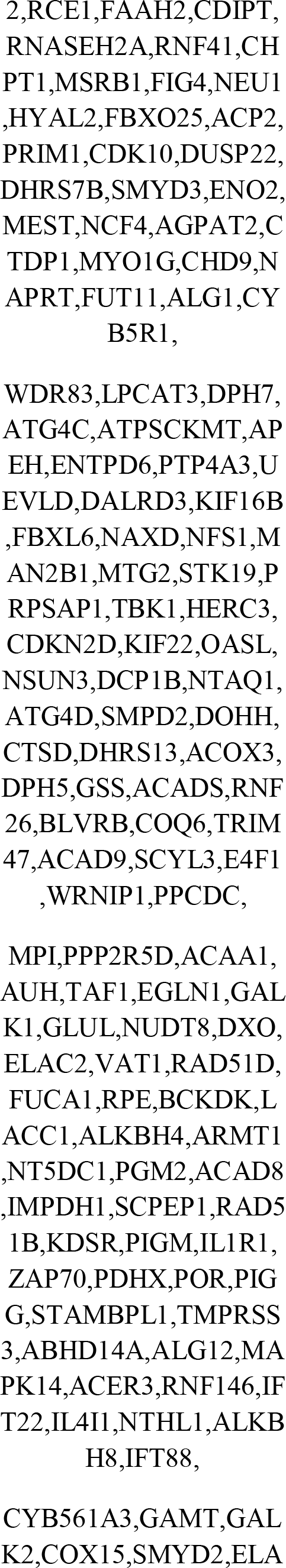
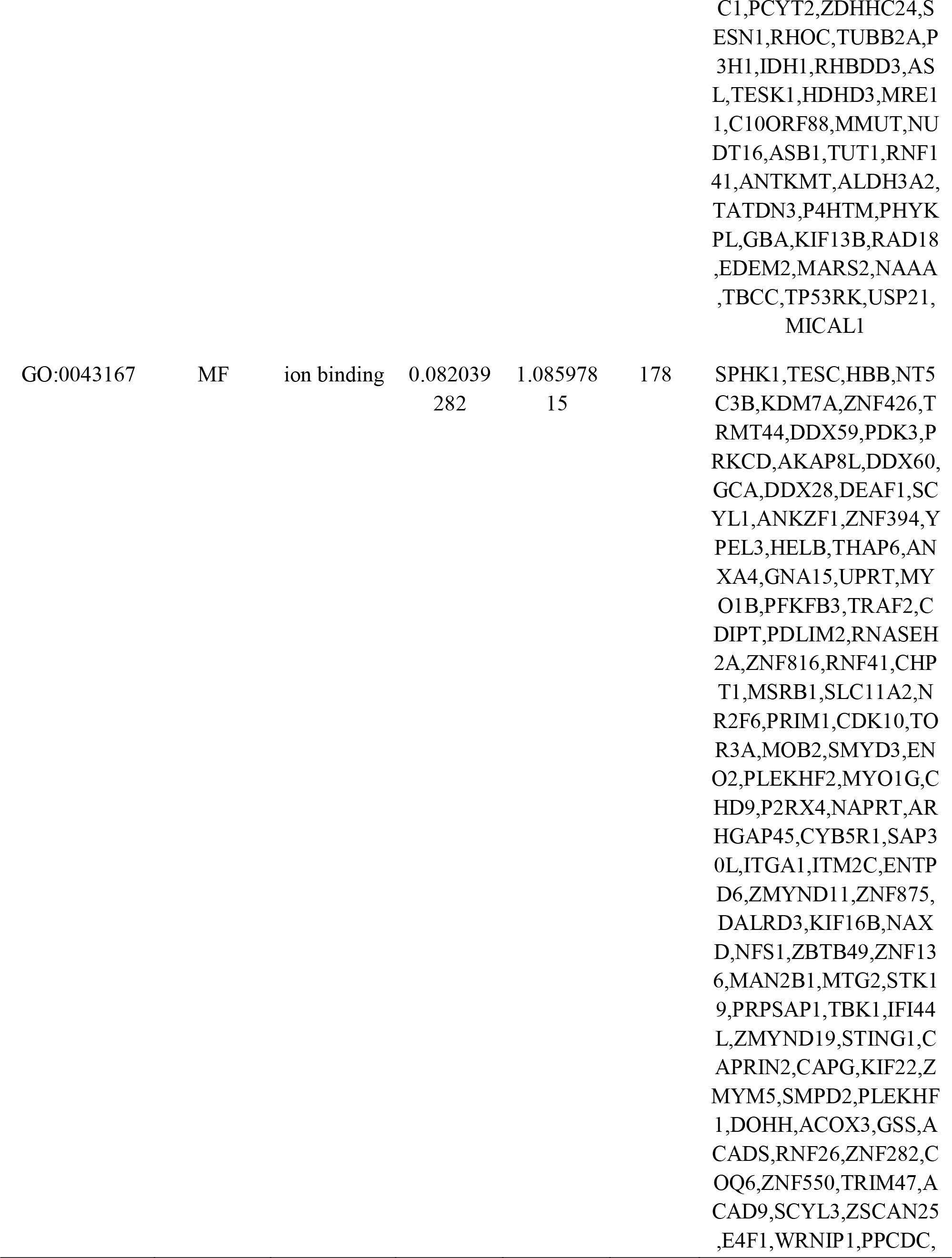
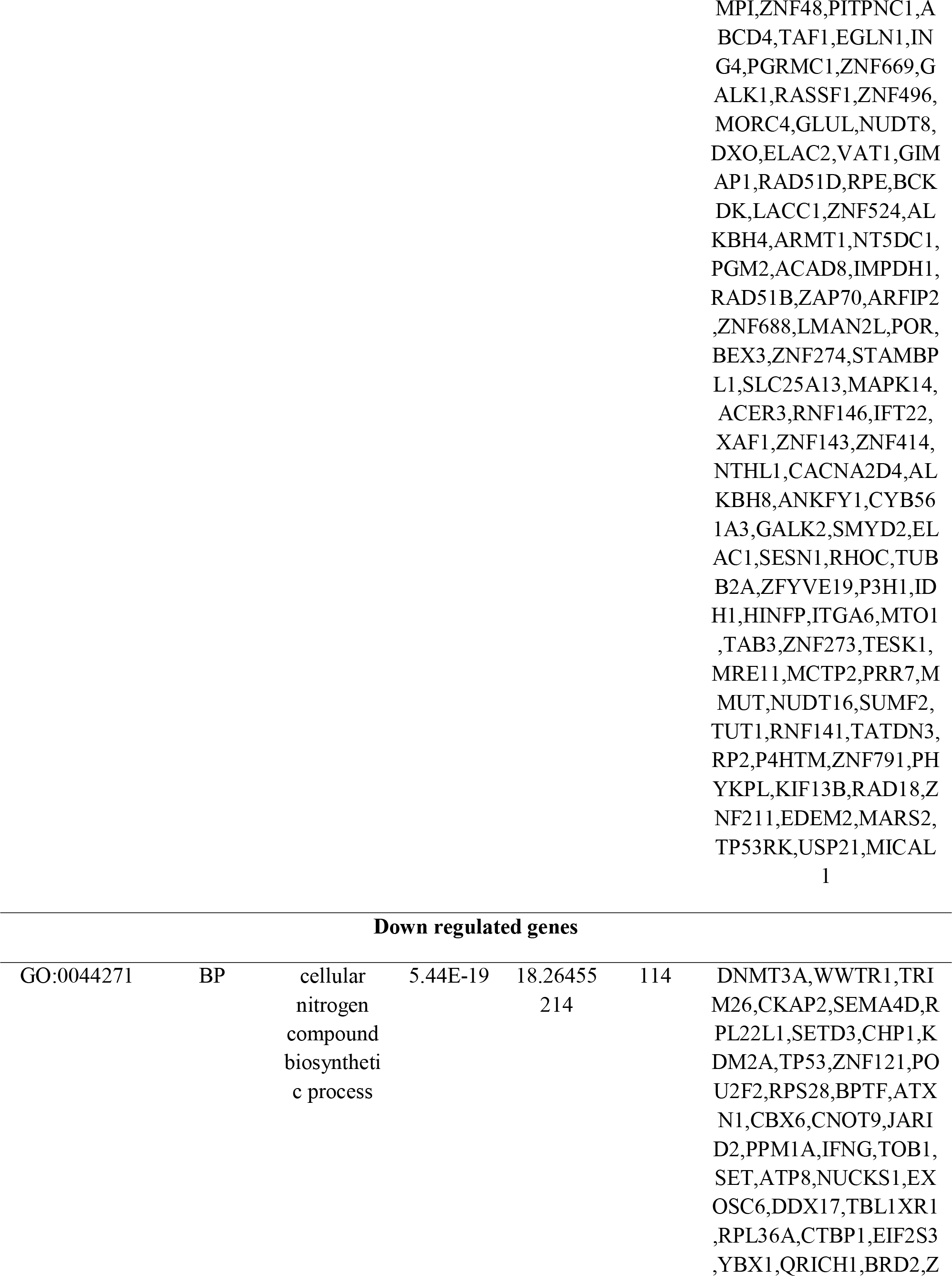
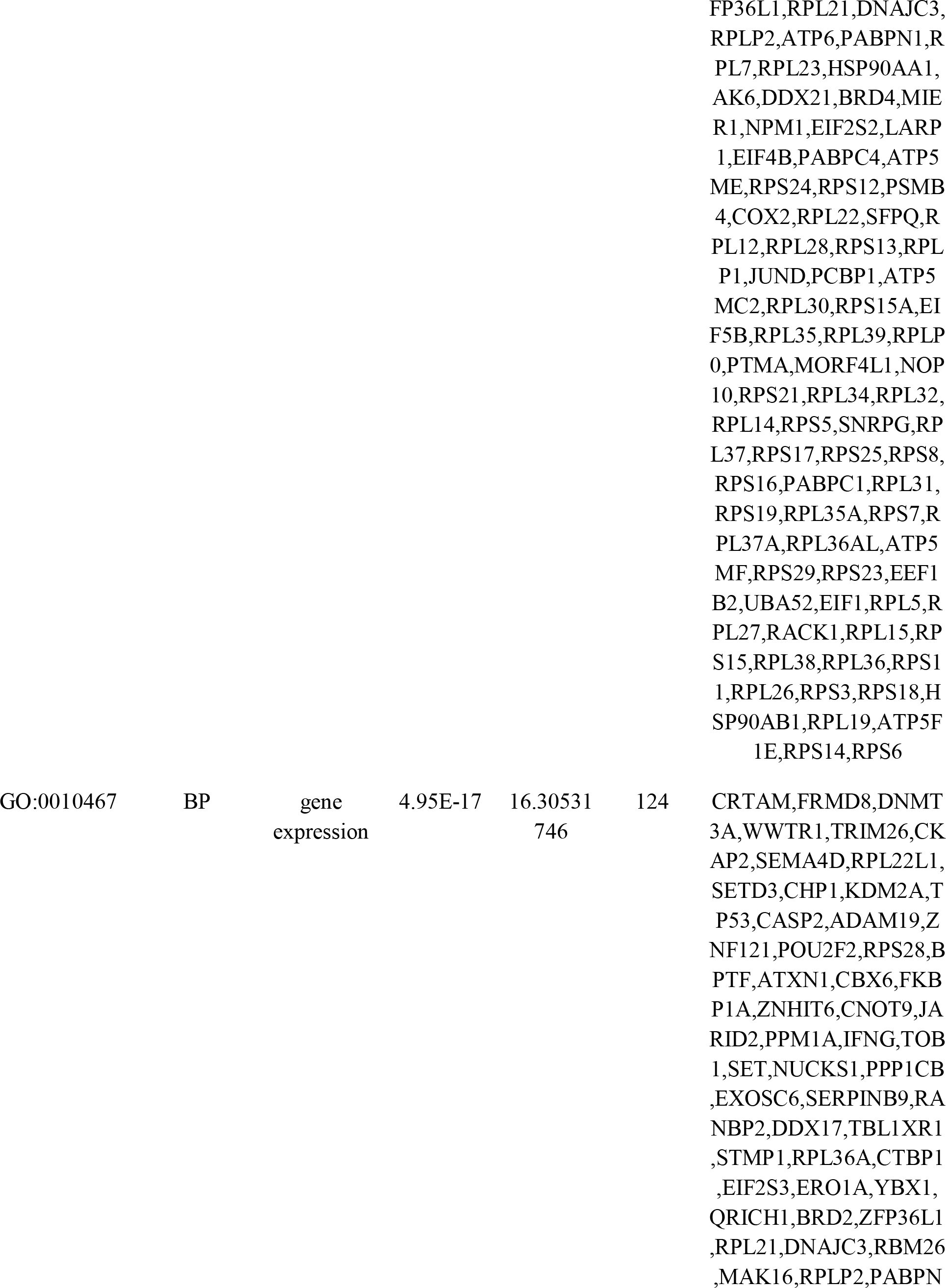
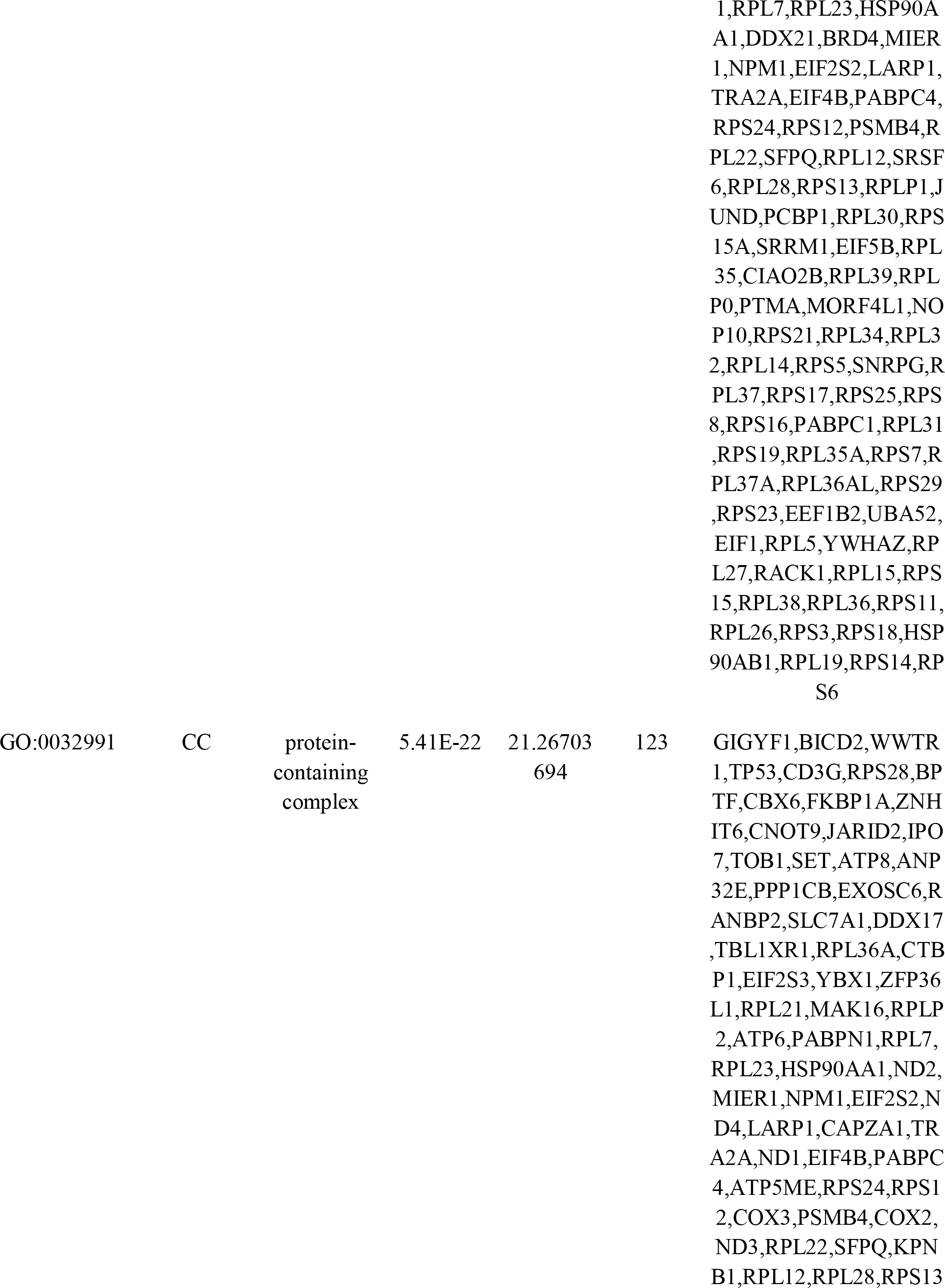
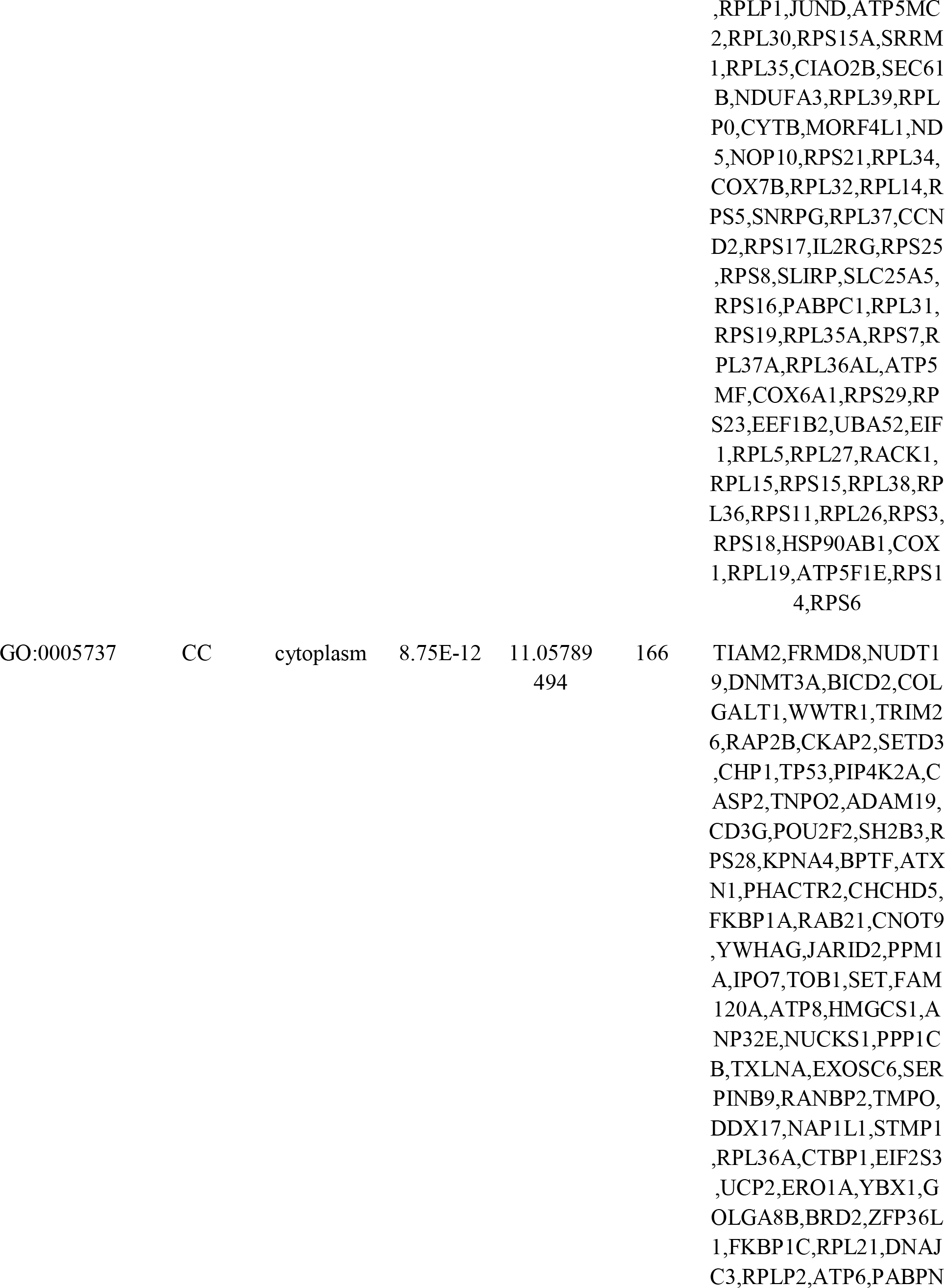
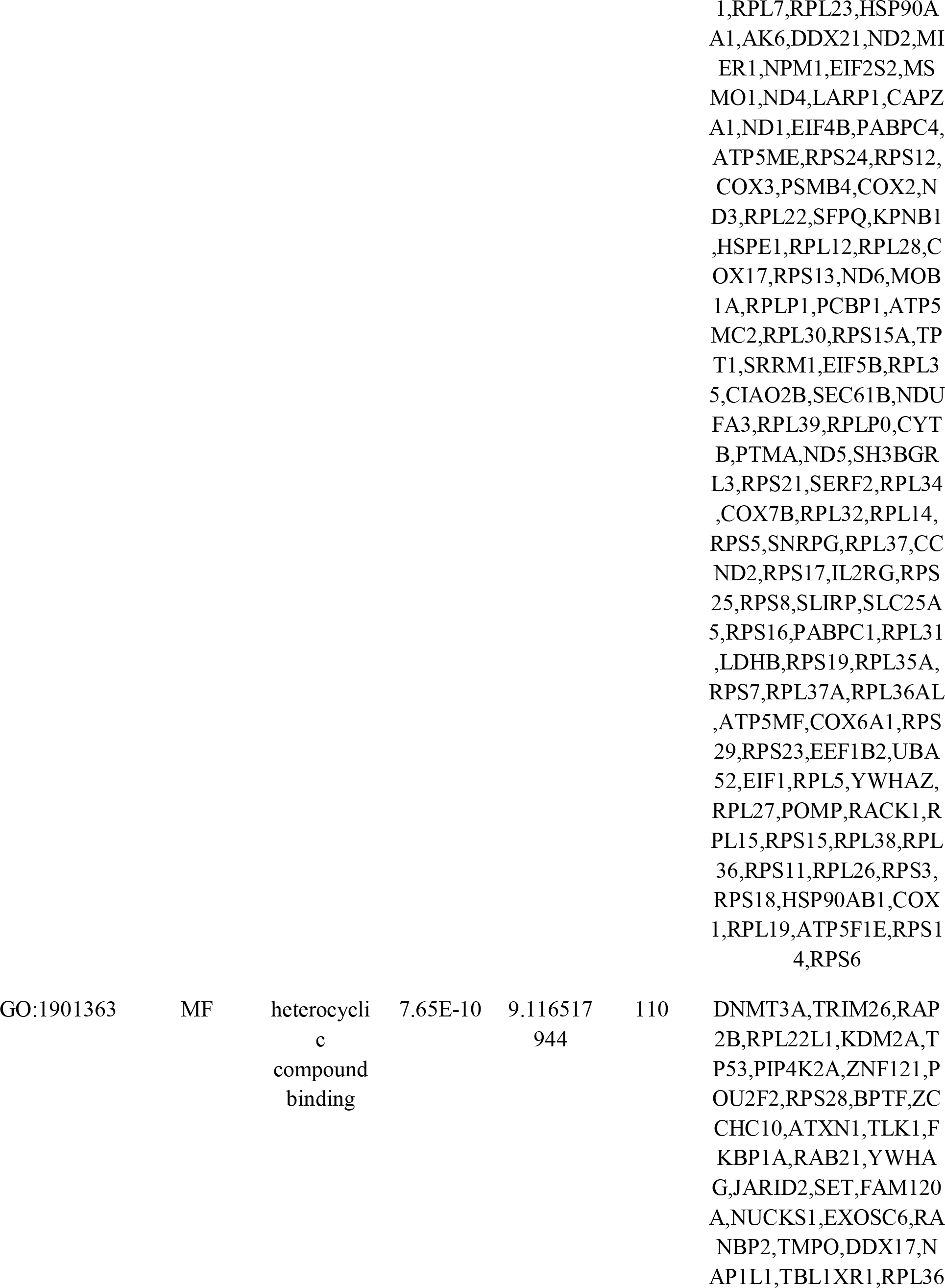
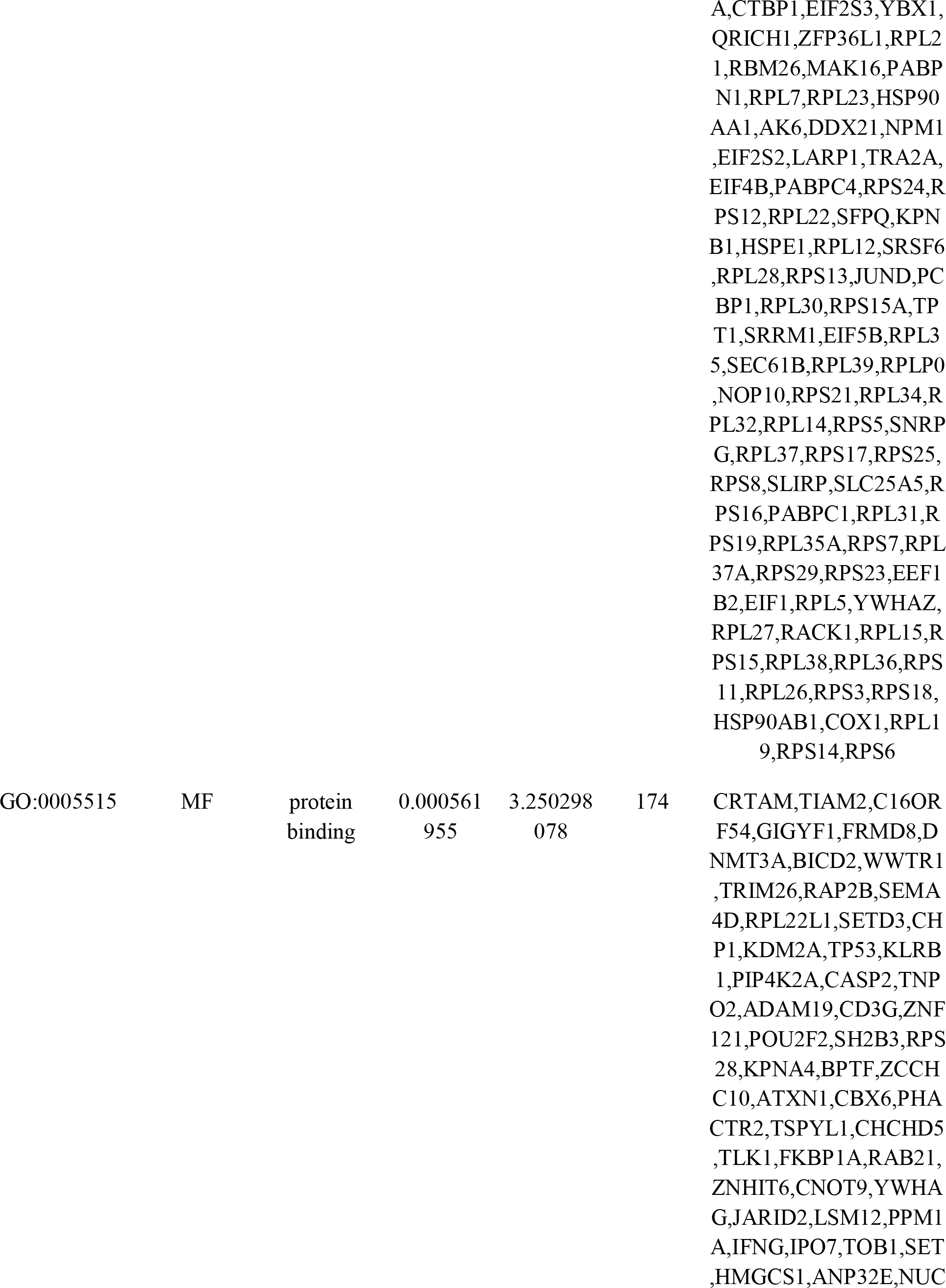
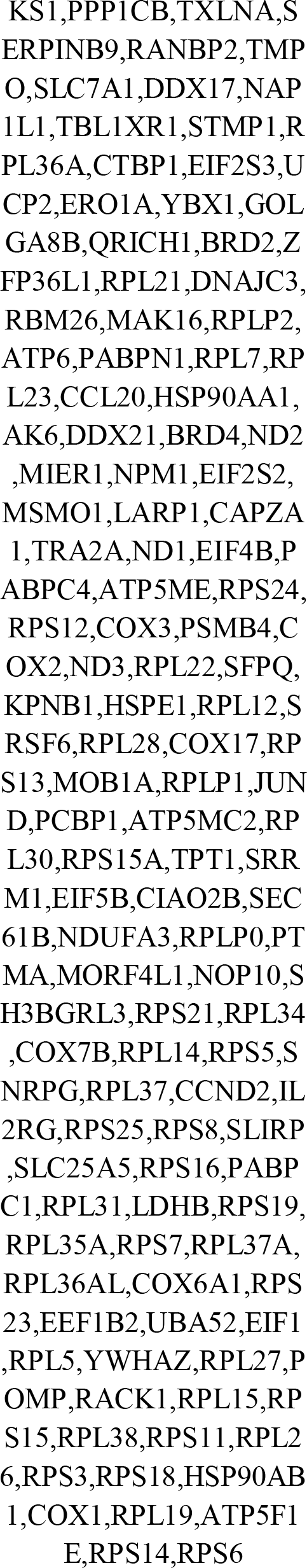
The enriched GO terms of the up and down regulated differentially expressed genes

**Table 3.** The enriched pathway terms of the up and down regulated differentially expressed genes

| Pathway ID | Pathway Name | Adjusted<br>p value | Negative<br>log10 of<br>adjusted<br>p value | Gene<br>Count | Gene |
| --- | --- | --- | --- | --- | --- |
| Up regulated genes |  |  |  |  |  |
| REAC:R-HSA-71387 | Metabolism of carbohydrates | 0.471762<br>741 | 0.326276<br>362 | 13 | DSE,ALDOC,PFKFB3,HYAL2,ENO2,FUT11,MAN2B1,PPP2R5D,GALK1,RPE,PGM2,SLC25A13,SLC25A1 |
| REAC:R-HSA-1430728 | Metabolism | 0.471762<br>741 | 0.326276<br>362 | 65 | SPHK1,DSE,ALDOC,SLC52A2,PKD3,INPP5K,GNNA15,AKR7A2,PFKFB3,FAAH2,CDIPT,CHPT1,FIG4,NEU1,HYAL2,DHRS7B,ENO2,AGPAT2,CHD9,NAPRT,FUT11,ACBD5,LPCAT3,ENTPD6,HILPDA,NAXD,MAN2B1,SMPD2,ACOX3,GSS,ACADS,BLVRB,COQ6,ACAD9,PPCDC,PPP2R5D,ACAA1,AUH,STARD10,ABCD4,GALK1,GLUL,RPE,BCKDK,PGM2,ACAD8,IMPDH1,KDSR,PDHX,POR,SLC25A13,ACER3,IL4I1,GAMT,COX15,PCYT2,IDH1,ASL,MUT,PIK3R1,NUDT16,SUMF2,PHYKPL,GBA,SLC25A1 |
| REAC:R-HSA-6798695 | Neutrophil degranulation | 0.471762<br>741 | 0.326276<br>362 | 23 | HBB,ALDOC,PRKCD,GCA,NEU1,TCIRG1,SNAIP29,AGPAT2,NAPRT,ARRHGAP45,APEH,MAN2B1,STING1,GRN,CTSD,ACAA1,PGRMC1,VAT1,FUCA1,PGM2,IMPDH1,MAPK14,IDH1 |
| REAC:R-HSA-556833 | Metabolism of lipids | 0.541235<br>611 | 0.266613<br>636 | 26 | SPHK1,INPP5K,FAAH2,CDIPT,CHPT1,FIG4,NEU1,DHRS7B,AGPAT2,CHD9,ACBD5,LPCAT3,HILPDA,SMPD2,ACOX3,ACADS,ACAA1,STARD10,KDSR,ACER3,PCYT2,MMUT,PIK3R1,SUMF2,GBA,SLC25A1 |
| REAC:R-HSA-168256 | Immune System | 0.668911<br>386 | 0.174631<br>412 | 60 | HBB,ALDOC,PRKCD,GCA,NFKBIE,ANAPC4,TRAFF2,RNF41,NEU1,TCIRG1,SNAP29,NCF4,AGPAT2,NAPRT,SIPA1,ARHGAP45,APEH,IFIT3,MAN2B1,TBK1,HERC3,STING1,BTBD6,GRN,KIF22,OASL,CTSD,PSTPIP1,PPP2R5D,TNFRSF14,IRF7,ACAA1,S1PR1,IL22,PGRMC1,VAT1,FUCA1,PGM2,IMPDH1,IL1R1,ZAP70,PDCD1,SOCS5,MAPK14,XAF1,IFIT2,TUBB2A,IDH1,GNLY,TAB3,TNFRSF12A,MRE11,CD70,IL15,PIK3R1,ASB1,ASB7,OSM,FLNB,SOCS3 |
| REAC:R-HSA-1280215 | Cytokine Signaling in Immune system | 0.681719<br>801 | 0.166394<br>091 | 23 | PRKCD,TRAF2,IFIT3,TBK1,OASL,PPP2R5D,TNFRSF14,IRF7,S1PR1,IL22,IL1R1,SOCS5,MAPK14,XAF1,IFIT2,TAB3,TNFRSF12A,CD70,IL15,PIK3R1,OSM,FLNB,SOCS3 |
| **Down regulated genes** |  |  |  |  |  |
| REAC:R-HSA-8953854 | Metabolism of RNA | 7.19E-40 | 39.14302<br>373 | 68 | RPL22L1,RPS28,CNOT9,SET,EXOSC6,RANBP2,RPL36A,YBX1,ZFP36L1 |
|  |  |  |  |  | ,RPL21,RPLP2,PABPN1,<br>RPL7,RPL23,DDX21,EI<br>F4B,RPS24,RPS12,PSM<br>B4,RPL22,RPL12,SRSF6<br>,RPL28,RPS13,RPLP1,P<br>CBP1,RPL30,RPS15A,S<br>RRM1,RPL35,RPL39,RP<br>LP0,NOP10,RPS21,RPL3<br>4,RPL32,RPL14,RPS5,S<br>NRPG,RPL37,RPS17,RP<br>S25,RPS8,RPS16,PABPC<br>1,RPL31,RPS19,RPL35A<br>,RPS7,RPL37A,RPL36A<br>L,RPS29,RPS23,UBA52,<br>RPL5,YWHAZ,RPL27,R<br>PL15,RPS15,RPL38,RPL<br>36,RPS11,RPL26,RPS3,<br>RPS18,RPL19,RPS14,RP<br>S6 |
| REAC:R-HSA-1430728 | Metabolism | 7.32E-23 | 22.13538<br>837 | 87 | TIAM2,NUDT19,RPL22<br>L1,CHP1,PIP4K2A,RPS2<br>8,ATP8,HMGCS1,PPP1C<br>B,RANBP2,TBL1XR1,R<br>PL36A,UCP2,RPL21,RP<br>LP2,ATP6,RPL7,RPL23,<br>HSP90AA1,AK6,ND2,M<br>SMO1,ND4,ND1,ATP5<br>ME,RPS24,RPS12,COX3<br>,PSMB4,COX2,ND3,RP<br>L22,KPNB1,RPL12,RPL<br>28,RPS13,ND6,RPLP1,A<br>TP5MC2,RPL30,RPS15<br>A,RPL35,CIAO2B,NDU<br>FA3,RPL39,RPLP0,CYT<br>B,ND5,RPS21,RPL34,C<br>OX7B,RPL32,RPL14,RP<br>S5,RPL37,RPS17,RPS25,<br>RPS8,RPS16,RPL31,LD<br>HB,RPS19,RPL35A,RPS<br>7,RPL37A,RPL36AL,AT<br>P5MF,COX6A1,RPS29,R<br>PS23,UBA52,RPL5,RPL<br>27,RPL15,RPS15,RPL38, |
|  |  |  |  |  | RPL36,RPS11,RPL26,RP<br>S3,RPS18,HSP90AB1,C<br>OX1,RPL19,ATP5F1E,R<br>PS14,RPS6 |
| REAC:R-HSA-392499 | Metabolism of<br>proteins | 3.13E-13 | 12.50511<br>44 | 71 | DNMT3A,RPL22L1,TP5<br>3,RPS28,RAB21,EXOSC<br>6,RANBP2,DDX17,RPL<br>36A,CTBP1,EIF2S3,ERO<br>1A,CD52,RPL21,DNAJC<br>3,RPLP2,RPL7,RPL23,N<br>PM1,EIF2S2,CAPZA1,EI<br>F4B,RPS24,RPS12,PSM<br>B4,RPL22,RPL12,RPL28<br>,RPS13,RPLP1,RPL30,R<br>PS15A,EIF5B,RPL35,SE<br>C61B,RPL39,RPLP0,RP<br>S21,RPL34,RPL32,RPL1<br>4,RPS5,RPL37,RPS17,R<br>PS25,RPS8,RPS16,PABP<br>C1,RPL31,RPS19,RPL35<br>A,RPS7,RPL37A,RPL36<br>AL,RPS29,RPS23,EEF1<br>B2,UBA52,RPL5,RPL27,<br>RPL15,RPS15,RPL38,RP<br>L36,RPS11,RPL26,RPS3,<br>RPS18,RPL19,RPS14,RP<br>S6 |
| REAC:R-HSA-1428517 | The citric acid<br>(TCA) cycle and<br>respiratory electron<br>transport | 1.40E-12 | 11.85532<br>452 | 21 | ATP8,UCP2,ATP6,ND2,<br>ND4,ND1,ATP5ME,CO<br>X3,COX2,ND3,ND6,AT<br>P5MC2,NDUFA3,CYTb,<br>ND5,COX7B,LDHB,AT<br>P5MF,COX6A1,COX1,A<br>TP5F1E |
| REAC:R-HSA-5628897 | TP53 Regulates<br>Metabolic Genes | 0.000485<br>535 | 3.313779<br>251 | 8 | TP53,YWHAG,COX3,C<br>OX2,COX7B,COX6A1,Y<br>WHAZ,COX1 |
| REAC:R-HSA-9013418 | RHOBTB2 GTPase<br>cycle | 0.032066<br>067 | 1.493954<br>3 | 2 | HSP90AA1,HSP90AB1 |

### Construction of the PPI and module analysis

PPI network of DEGs, consisting of 4200 nodes and 13663 edges (Fig. 3), was constructed by Cytoscape software, based on STRING database. The DEGs with high node degree, betweenness centrality, stress centrality and closeness centrality were selected as the hub genes of T1DM. These hub genes were MAPK14, RHOC, MAD2L1, TAF1, TRAF2, HSP90AA1, TP53, HSP90AB1, UBA52 and RACK1, which might play a critical role in T1DM progression (Table 4). Plug-ins PEWCC1 was used to carry out module analysis in Cytoscape software. According the degree of importance, we chose 2 significant modules from the PPI network complex for further analysis using PEWCC1. GO term and REACTOME pathway enrichment analysis showed that Module 1 consisted of 98 nodes and 119 edges (Fig.4A) which are mainly associated with immune system, cytokine signaling in immune system, cellular metabolic process and intracellular anatomical structure, and that Module 2 consisted of 99 nodes and 225 edges (Fig.4B), which are mainly associated with cellular nitrogen compound biosynthetic process, metabolism of RNA, metabolism, gene expression and protein-containing complex.

**Fig. 3.**
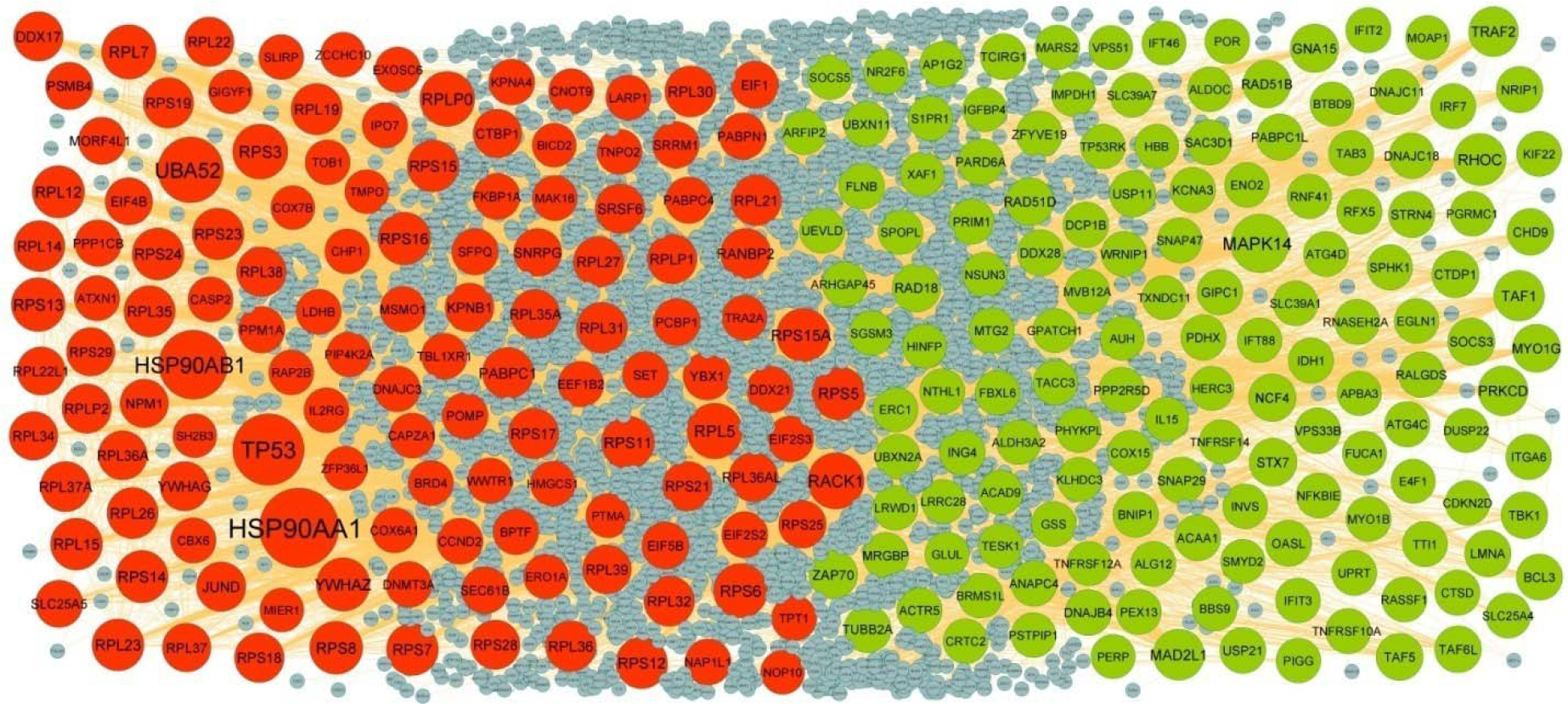
PPI network of DEGs. Up regulated genes are marked in green; down regulated genes are marked in red

**Fig. 4.**
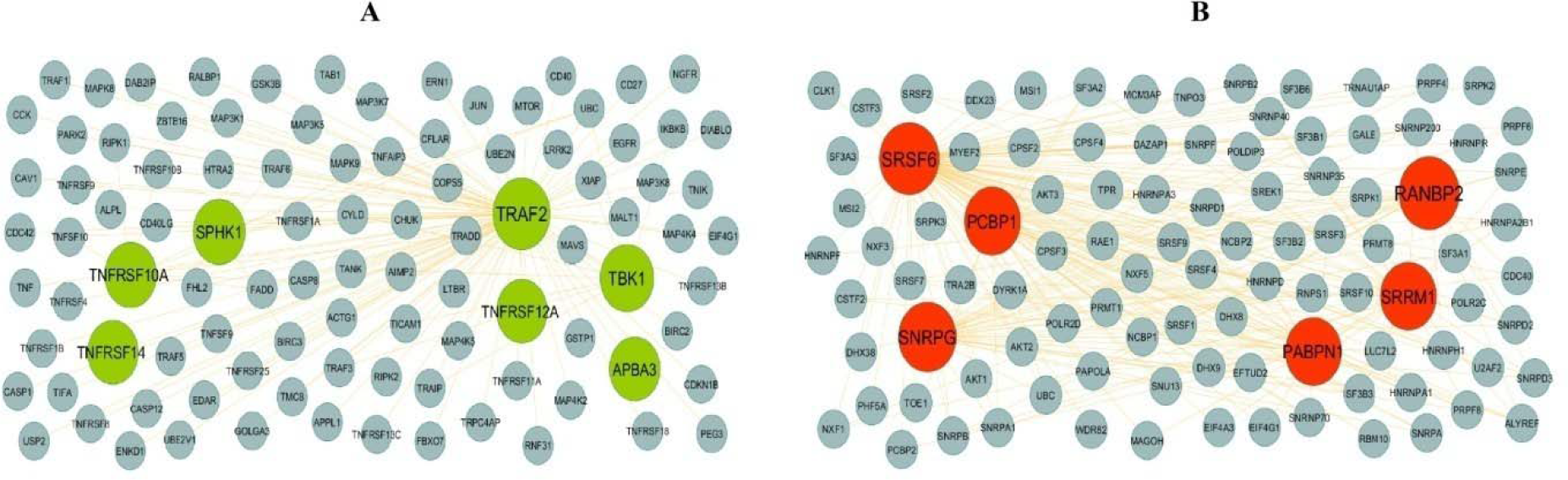
Modules of isolated form PPI of DEGs. (A) The most significant module was obtained from PPI network with 98 nodes and 119 edges for up regulated genes (B) The most significant module was obtained from PPI network with 99 nodes and 225 edges for down regulated genes. Up regulated genes are marked in green; down regulated genes are marked in red

**Table 4.** Topology table for up and down regulated genes

| **Regulation** | **Node** | **Degree** | **Betweenness** | **Stress** | **Closeness** |
| --- | --- | --- | --- | --- | --- |
| Up | MAPK14 | 248 | 0.085274 | 15728642 | 0.38312 |
| Up | RHOC | 139 | 0.043685 | 9574126 | 0.341188 |
| Up | MAD2L1 | 120 | 0.026964 | 7656784 | 0.329488 |
| Up | TAF1 | 113 | 0.024085 | 5459358 | 0.343112 |
| Up | TRAF2 | 99 | 0.026067 | 4924990 | 0.339917 |
| Up | PRKCD | 98 | 0.018476 | 3147294 | 0.368883 |
| Up | RAD18 | 76 | 0.016617 | 5141842 | 0.322975 |
| Up | ZAP70 | 76 | 0.020284 | 2506796 | 0.348957 |
| Up | GNA15 | 75 | 0.026977 | 5192140 | 0.295371 |
| Up | MYO1G | 70 | 0.01742 | 5735546 | 0.320657 |
| Up | RAD51D | 67 | 0.007022 | 3265710 | 0.291173 |
| Up | RAD51B | 64 | 0.006077 | 2970544 | 0.291274 |
| Up | STX7 | 63 | 0.01517 | 2950824 | 0.320583 |
| Up | NCF4 | 58 | 0.012981 | 2734518 | 0.292776 |
| Up | TUBB2A | 56 | 0.01216 | 2469216 | 0.333704 |
| Up | CTDP1 | 55 | 0.011258 | 7283228 | 0.276742 |
| Up | SOCS3 | 54 | 0.008802 | 3163230 | 0.320314 |
| Up | TAF5 | 51 | 0.007187 | 1831668 | 0.33978 |
| Up | TAF6L | 48 | 0.004381 | 1252990 | 0.323224 |
| Up | CHD9 | 47 | 0.008975 | 2713114 | 0.281001 |
| Up | TBK1 | 45 | 0.010143 | 1758800 | 0.340413 |
| Up | SNAP29 | 43 | 0.008774 | 2281774 | 0.320119 |
| Up | PABPC1L | 41 | 0.002339 | 773678 | 0.320461 |
| Up | NRIP1 | 40 | 0.005404 | 1502852 | 0.289227 |
| Up | ZFYVE19 | 40 | 0.007074 | 831294 | 0.299714 |
| Up | PARD6A | 39 | 0.009159 | 1882504 | 0.319486 |
| Up | USP11 | 39 | 0.011354 | 1197114 | 0.337377 |
| Up | LMNA | 39 | 0.00873 | 1524904 | 0.340717 |
| Up | STRN4 | 38 | 0.007387 | 1889442 | 0.288472 |
| Up | OASL | 37 | 0.005287 | 1062404 | 0.314132 |
| Up | DDX28 | 37 | 0.005105 | 2699210 | 0.264454 |
| Up | BCL3 | 36 | 0.003112 | 1161430 | 0.325681 |
| Up | USP21 | 34 | 0.008469 | 1803250 | 0.317937 |
| Up | TCIRG1 | 33 | 0.013659 | 1152612 | 0.320975 |
| Up | ACAA1 | 33 | 0.009413 | 1045108 | 0.259262 |
| Up | DNAJB4 | 33 | 0.001099 | 321606 | 0.337377 |
| Up | MRGBP | 32 | 0.003878 | 608560 | 0.281095 |
| Up | ING4 | 32 | 0.003996 | 936048 | 0.29884 |
| Up | PPP2R5D | 31 | 0.005832 | 1555198 | 0.318879 |
| Up | VPS33B | 30 | 0.00702 | 1230966 | 0.318661 |
| Up | BNIP1 | 29 | 0.003774 | 1198206 | 0.317553 |
| Up | ANAPC4 | 29 | 0.004601 | 1009406 | 0.320363 |
| Up | MTG2 | 29 | 7.05E-06 | 7402 | 0.280326 |
| Up | RALGDS | 29 | 0.002388 | 694622 | 0.279877 |
| Up | FLNB | 29 | 0.008802 | 1271332 | 0.318323 |
| Up | AP1G2 | 28 | 0.011992 | 1106430 | 0.3165 |
| Up | DNAJC11 | 28 | 0.001638 | 330744 | 0.337242 |
| Up | GIPC1 | 26 | 0.009418 | 1109090 | 0.318734 |
| Up | DCP1B | 25 | 0.010044 | 1042580 | 0.319218 |
| Up | DNAJC18 | 25 | 1.89E-04 | 84208 | 0.300293 |
| Up | SAC3D1 | 25 | 0.003215 | 617792 | 0.270938 |
| Up | ITGA6 | 25 | 0.006779 | 1004692 | 0.28151 |
| Up | ACTR5 | 24 | 0.003106 | 2091902 | 0.272928 |
| Up | KCNA3 | 24 | 0.008393 | 2417954 | 0.248227 |
| Up | PDHX | 22 | 0.006616 | 2110408 | 0.248007 |
| Up | ATG4C | 22 | 0.005663 | 2236998 | 0.256898 |
| Up | IRF7 | 22 | 0.002609 | 639414 | 0.320461 |
| Up | UEVLD | 22 | 0.0042 | 627058 | 0.319388 |
| Up | SPHK1 | 21 | 0.006193 | 1135846 | 0.319973 |
| Up | PRIM1 | 21 | 0.002968 | 578352 | 0.317553 |
| Up | IDH1 | 20 | 0.007106 | 661676 | 0.317745 |
| Up | RFX5 | 20 | 0.004228 | 609476 | 0.288929 |
| Up | WRNIP1 | 20 | 0.00221 | 458594 | 0.321368 |
| Up | UBXN11 | 20 | 0.001746 | 1014346 | 0.2543 |
| Up | KIF22 | 19 | 0.002522 | 474830 | 0.319826 |
| Up | TNFRSF10A | 19 | 0.003621 | 505748 | 0.32051 |
| Up | CTSD | 19 | 0.006483 | 747530 | 0.318202 |
| Up | ATG4D | 18 | 0.001153 | 548318 | 0.238471 |
| Up | TAB3 | 18 | 0.003592 | 444596 | 0.331413 |
| Up | IFIT2 | 18 | 0.00145 | 311530 | 0.304033 |
| Up | RNASEH2A | 17 | 0.003297 | 612270 | 0.316762 |
| Up | NSUN3 | 17 | 0.001913 | 2399130 | 0.241308 |
| Up | ARHGAP45 | 17 | 0.002169 | 351820 | 0.27364 |
| Up | TTI1 | 17 | 0.001537 | 999004 | 0.273586 |
| Up | NTHL1 | 16 | 0.001542 | 580194 | 0.243774 |
| Up | SLC25A4 | 16 | 0.004359 | 1429122 | 0.318154 |
| Up | PEX13 | 16 | 0.005743 | 2746072 | 0.201391 |
| Up | MYO1B | 16 | 0.002138 | 662962 | 0.318444 |
| Up | SNAP47 | 16 | 5.49E-04 | 54264 | 0.250179 |
| Up | ERC1 | 16 | 0.003935 | 677408 | 0.316428 |
| Up | UPRT | 16 | 0.003911 | 1567784 | 0.257024 |
| Up | BRMS1L | 15 | 0.001104 | 529978 | 0.265877 |
| Up | COX15 | 15 | 0.005257 | 4347496 | 0.246117 |
| Up | TACC3 | 15 | 0.003933 | 511960 | 0.316285 |
| Up | UBXN2A | 15 | 0.003198 | 626068 | 0.316691 |
| Up | RASSF1 | 15 | 0.001065 | 255298 | 0.277546 |
| Up | GSS | 14 | 0.004921 | 562258 | 0.317361 |
| Up | LRWD1 | 14 | 0.003048 | 279620 | 0.336512 |
| Up | SLC39A1 | 14 | 0.005731 | 763412 | 0.204102 |
| Up | PGRMC1 | 13 | 0.003258 | 929272 | 0.316738 |
| Up | ALDOC | 13 | 0.003983 | 526508 | 0.316071 |
| Up | SOCS5 | 13 | 2.18E-04 | 123360 | 0.258639 |
| Up | ALG12 | 13 | 0.004287 | 2097716 | 0.221396 |
| Up | CRTC2 | 13 | 4.60E-04 | 111786 | 0.323348 |
| Up | IFIT3 | 13 | 1.66E-04 | 65032 | 0.300573 |
| Up | POR | 13 | 0.005042 | 852768 | 0.31693 |
| Up | INVS | 12 | 0.003378 | 610266 | 0.229228 |
| Up | NFKBIE | 12 | 0.002056 | 1131960 | 0.257497 |
| Up | CDKN2D | 12 | 4.13E-05 | 28664 | 0.269928 |
| Up | TESK1 | 12 | 0.003862 | 1051194 | 0.248109 |
| Up | IMPDH1 | 12 | 0.002739 | 383906 | 0.317721 |
| Up | TP53RK | 12 | 0.004387 | 322116 | 0.331884 |
| Up | PSTPIP1 | 12 | 0.002918 | 971936 | 0.258974 |
| Up | ENO2 | 11 | 0.001402 | 544636 | 0.249288 |
| Up | MARS2 | 11 | 0.003351 | 852330 | 0.234006 |
| Up | ACAD9 | 11 | 0.002298 | 1498752 | 0.247437 |
| Up | S1PR1 | 11 | 0.001616 | 289100 | 0.316214 |
| Up | MVB12A | 11 | 0.001579 | 327132 | 0.316882 |
| Up | FBXL6 | 11 | 0.001762 | 246360 | 0.315762 |
| Up | ALDH3A2 | 11 | 0.004671 | 696342 | 0.317145 |
| Up | SPOPL | 11 | 0.00155 | 550770 | 0.262766 |
| Up | IFT88 | 11 | 0.004758 | 566990 | 0.218721 |
| Up | EGLN1 | 11 | 0.001927 | 1082624 | 0.258051 |
| Up | PIGG | 11 | 0.004758 | 735790 | 0.195767 |
| Up | BBS9 | 10 | 0.003337 | 1282716 | 0.231541 |
| Up | ARFIP2 | 10 | 0.001525 | 604576 | 0.262241 |
| Up | SGSM3 | 10 | 0.001528 | 165830 | 0.249051 |
| Up | PHYKPL | 10 | 0.00288 | 1111518 | 0.20453 |
| Up | NR2F6 | 10 | 0.001499 | 762172 | 0.24786 |
| Up | RNF41 | 10 | 0.001032 | 174526 | 0.316285 |
| Up | LRRC28 | 2 | 0 | 0 | 0.294708 |
| Up | MOAP1 | 2 | 0 | 0 | 0.244426 |
| Up | E4F1 | 2 | 4.62E-05 | 22816 | 0.290146 |
| Up | SMYD2 | 2 | 0 | 0 | 0.309501 |
| Up | VPS51 | 1 | 0 | 0 | 0.179475 |
| Up | IFT46 | 1 | 0 | 0 | 0.179475 |
| Up | GLUL | 1 | 0 | 0 | 0.292593 |
| Up | IL15 | 1 | 0 | 0 | 0.240327 |
| Up | TXNDC11 | 1 | 0 | 0 | 0.240451 |
| Up | GPATCH1 | 1 | 0 | 0 | 0.258098 |
| Up | HERC3 | 1 | 0 | 0 | 0.278559 |
| Up | KLHDC3 | 1 | 0 | 0 | 0.241794 |
| Up | IGFBP4 | 1 | 0 | 0 | 0.241405 |
| Up | APBA3 | 1 | 0 | 0 | 0.253701 |
| Up | HINFP | 1 | 0 | 0 | 0.289566 |
| Up | TNFRSF12A | 1 | 0 | 0 | 0.253701 |
| Up | HBB | 1 | 0 | 0 | 0.25775 |
| Up | DUSP22 | 1 | 0 | 0 | 0.277015 |
| Up | TNFRSF14 | 1 | 0 | 0 | 0.253701 |
| Up | XAF1 | 1 | 0 | 0 | 0.289566 |
| Up | FUCA1 | 1 | 0 | 0 | 0.241253 |
| Up | SLC39A7 | 1 | 0 | 0 | 0.169513 |
| Up | AUH | 1 | 0 | 0 | 0.205894 |
| Up | BTBD9 | 1 | 0 | 0 | 0.241253 |
| Up | PERP | 1 | 0 | 0 | 0.289566 |
| Down | HSP90AA1 | 620 | 0.172854 | 42572554 | 0.413572 |
| Down | TP53 | 479 | 0.165254 | 40252372 | 0.407551 |
| Down | HSP90AB1 | 437 | 0.069176 | 19212422 | 0.392871 |
| Down | UBA52 | 379 | 0.090169 | 23709030 | 0.386079 |
| Down | RACK1 | 214 | 0.02696 | 9421194 | 0.358705 |
| Down | RPL5 | 202 | 0.017462 | 4964436 | 0.380862 |
| Down | RPS3 | 195 | 0.011749 | 4196160 | 0.368172 |
| Down | RPLP0 | 191 | 0.009668 | 3521194 | 0.362201 |
| Down | RPL7 | 184 | 0.007353 | 3033322 | 0.368948 |
| Down | RPS6 | 178 | 0.009932 | 3621206 | 0.367206 |
| Down | RPS13 | 171 | 0.009257 | 3062900 | 0.35892 |
| Down | RPS5 | 167 | 0.004438 | 2164900 | 0.347341 |
| Down | YWHAZ | 167 | 0.038337 | 6099060 | 0.374911 |
| Down | RPS15A | 165 | 0.005005 | 2279708 | 0.363455 |
| Down | RPS8 | 157 | 0.004145 | 2168132 | 0.347859 |
| Down | RPS11 | 155 | 0.002739 | 1589330 | 0.347484 |
| Down | RPS16 | 151 | 0.005525 | 2041378 | 0.362139 |
| Down | RPS14 | 146 | 0.004872 | 2010096 | 0.360336 |
| Down | RPL12 | 145 | 0.003174 | 1296430 | 0.345625 |
| Down | RPS24 | 141 | 0.001737 | 1190126 | 0.345995 |
| Down | RPL15 | 140 | 0.001768 | 1208442 | 0.346824 |
| Down | RPS7 | 140 | 0.003166 | 1696606 | 0.357514 |
| Down | RPL23 | 139 | 0.003517 | 1622304 | 0.358093 |
| Down | RPS23 | 135 | 0.001977 | 1199702 | 0.356603 |
| Down | RPS15 | 135 | 0.005952 | 2032926 | 0.345824 |
| Down | RPS12 | 134 | 0.003274 | 1376082 | 0.348638 |
| Down | RPL19 | 128 | 0.004089 | 1543280 | 0.351675 |
| Down | RPL35 | 128 | 0.001404 | 872788 | 0.344887 |
| Down | PABPC1 | 127 | 0.023684 | 7393480 | 0.347226 |
| Down | RPL26 | 126 | 0.003244 | 1507600 | 0.357849 |
| Down | RPL30 | 125 | 0.001193 | 949910 | 0.344745 |
| Down | RPL14 | 121 | 9.40E-04 | 734188 | 0.34714 |
| Down | RPL31 | 120 | 0.001651 | 962208 | 0.347312 |
| Down | RPL36 | 119 | 0.001724 | 795722 | 0.344944 |
| Down | RPL21 | 116 | 4.60E-04 | 552056 | 0.345199 |
| Down | RPS19 | 116 | 0.003536 | 1021154 | 0.34554 |
| Down | RPL27 | 113 | 0.001878 | 843532 | 0.345369 |
| Down | RPS17 | 112 | 3.43E-04 | 540936 | 0.342943 |
| Down | RPL32 | 112 | 0.002687 | 1221610 | 0.360151 |
| Down | RANBP2 | 110 | 0.03413 | 5339474 | 0.375346 |
| Down | RPL37A | 109 | 8.85E-04 | 590672 | 0.344859 |
| Down | RPL38 | 100 | 0.002323 | 833852 | 0.34272 |
| Down | RPS18 | 100 | 1.19E-04 | 145130 | 0.308659 |
| Down | SRSF6 | 98 | 0.016104 | 5303426 | 0.328998 |
| Down | JUND | 97 | 0.014853 | 4410440 | 0.34069 |
| Down | RPLP1 | 97 | 0.001121 | 557658 | 0.341188 |
| Down | DDX17 | 97 | 0.016264 | 5304838 | 0.345455 |
| Down | RPL22 | 96 | 8.95E-04 | 532836 | 0.343 |
| Down | YWHAG | 95 | 0.012293 | 2256044 | 0.361515 |
| Down | RPLP2 | 95 | 2.45E-04 | 352522 | 0.340607 |
| Down | RPS21 | 95 | 0.001606 | 586560 | 0.345767 |
| Down | RPS28 | 94 | 7.27E-04 | 488270 | 0.344068 |
| Down | RPL36A | 91 | 3.93E-05 | 83604 | 0.31385 |
| Down | RPL36AL | 90 | 1.78E-04 | 287190 | 0.342105 |
| Down | RPL35A | 89 | 3.67E-04 | 450232 | 0.342356 |
| Down | RPL39 | 85 | 8.04E-04 | 570124 | 0.343364 |
| Down | KPNB1 | 83 | 0.019658 | 5199570 | 0.328355 |
| Down | RPL34 | 82 | 2.42E-04 | 364792 | 0.339945 |
| Down | RPL37 | 81 | 5.13E-04 | 406440 | 0.339313 |
| Down | CTBP1 | 81 | 0.028114 | 4378160 | 0.341521 |
| Down | RPS29 | 79 | 3.58E-04 | 317560 | 0.339258 |
| Down | NPM1 | 78 | 0.012757 | 3155324 | 0.351469 |
| Down | RPL22L1 | 76 | 0.001122 | 440010 | 0.339039 |
| Down | YBX1 | 74 | 0.011705 | 3045144 | 0.345 |
| Down | SNRPG | 70 | 0.013186 | 3815654 | 0.321985 |
| Down | PSMB4 | 62 | 0.006753 | 1735816 | 0.331125 |
| Down | RPS25 | 62 | 1.55E-04 | 225554 | 0.337975 |
| Down | EIF4B | 61 | 0.004919 | 1856768 | 0.329669 |
| Down | MORF4L1 | 61 | 0.011893 | 3532114 | 0.324473 |
| Down | SLC25A5 | 59 | 0.010774 | 2239390 | 0.351322 |
| Down | EIF5B | 58 | 0.003783 | 975898 | 0.347859 |
| Down | EIF1 | 58 | 0.001784 | 538584 | 0.332252 |
| Down | PCBP1 | 55 | 0.00897 | 1866076 | 0.327305 |
| Down | SEC61B | 52 | 0.012585 | 3292532 | 0.323797 |
| Down | PABPN1 | 48 | 0.005608 | 1194108 | 0.327842 |
| Down | DDX21 | 48 | 0.004886 | 1206590 | 0.33063 |
| Down | PPP1CB | 47 | 0.010348 | 2416792 | 0.320608 |
| Down | PPM1A | 47 | 0.008289 | 1791912 | 0.3305 |
| Down | SRRM1 | 47 | 0.003736 | 1423870 | 0.284485 |
| Down | PABPC4 | 45 | 0.002845 | 959888 | 0.321221 |
| Down | SFPQ | 44 | 0.00625 | 1834782 | 0.32243 |
| Down | FKBP1A | 42 | 0.010368 | 1512446 | 0.343252 |
| Down | NAP1L1 | 40 | 0.00809 | 1625742 | 0.333307 |
| Down | CCND2 | 39 | 0.006869 | 1999478 | 0.318734 |
| Down | EIF2S2 | 39 | 0.002146 | 612642 | 0.333148 |
| Down | SET | 38 | 0.00902 | 2128256 | 0.325504 |
| Down | EIF2S3 | 37 | 0.003735 | 729686 | 0.332226 |
| Down | POMP | 37 | 0.009677 | 1871584 | 0.320657 |
| Down | KPNA4 | 36 | 0.006172 | 957132 | 0.337703 |
| Down | DNMT3A | 34 | 0.007674 | 4549464 | 0.270345 |
| Down | DNAJC3 | 34 | 0.00181 | 379954 | 0.337405 |
| Down | TBL1XR1 | 34 | 0.00652 | 1196036 | 0.325756 |
| Down | MSMO1 | 32 | 0.010962 | 2113600 | 0.317841 |
| Down | EEF1B2 | 30 | 0.007297 | 1308518 | 0.335813 |
| Down | BPTF | 30 | 0.006522 | 4067568 | 0.255026 |
| Down | WWTR1 | 27 | 0.003574 | 707626 | 0.291678 |
| Down | EXOSC6 | 27 | 0.008176 | 594646 | 0.254778 |
| Down | ATXN1 | 25 | 0.004093 | 805872 | 0.297907 |
| Down | IPO7 | 25 | 0.003925 | 590152 | 0.337486 |
| Down | COX6A1 | 24 | 0.008594 | 2281402 | 0.212017 |
| Down | CNOT9 | 24 | 0.004063 | 1757210 | 0.24855 |
| Down | CBX6 | 24 | 0.007097 | 968366 | 0.317962 |
| Down | TPT1 | 24 | 0.003131 | 604626 | 0.344124 |
| Down | BRD4 | 23 | 0.001753 | 1249076 | 0.280288 |
| Down | CASP2 | 22 | 0.003547 | 2071912 | 0.270554 |
| Down | LDHB | 21 | 0.007543 | 1331780 | 0.317986 |
| Down | CAPZA1 | 21 | 0.004999 | 1074858 | 0.319316 |
| Down | LARP1 | 20 | 0.001895 | 309726 | 0.324047 |
| Down | TNPO2 | 20 | 0.003444 | 525134 | 0.317433 |
| Down | TRA2A | 19 | 0.003342 | 587256 | 0.316309 |
| Down | ZCCHC10 | 19 | 0.003441 | 411010 | 0.26539 |
| Down | RAP2B | 19 | 0.0079 | 958974 | 0.319146 |
| Down | MAK16 | 19 | 0.002785 | 472748 | 0.316476 |
| Down | IL2RG | 19 | 0.005094 | 794848 | 0.316333 |
| Down | TOB1 | 18 | 0.003684 | 546506 | 0.287544 |
| Down | HMGCS1 | 18 | 0.010419 | 1189944 | 0.317986 |
| Down | CHP1 | 18 | 0.005871 | 484124 | 0.335625 |
| Down | PIP4K2A | 18 | 0.002977 | 517300 | 0.319632 |
| Down | ERO1A | 18 | 0.004085 | 664272 | 0.316547 |
| Down | SLIRP | 18 | 0.002816 | 474836 | 0.320583 |
| Down | TMPO | 1 | 0 | 0 | 0.254146 |
| Down | PTMA | 1 | 0 | 0 | 0.245584 |
| Down | GIGYF1 | 1 | 0 | 0 | 0.19908 |
| Down | NOP10 | 1 | 0 | 0 | 0.258527 |
| Down | ZFP36L1 | 1 | 0 | 0 | 0.277015 |
| Down | SH2B3 | 1 | 0 | 0 | 0.258702 |
| Down | BICD2 | 1 | 0 | 0 | 0.272928 |
| Down | MIER1 | 1 | 0 | 0 | 0.219429 |
| Down | COX7B | 1 | 0 | 0 | 0.174936 |

### miRNA-hub gene regulatory network construction

miRNA-hub gene regulatory network constructed by miRNet database was adjusted and visualized by Cytoscape. As shown in Fig 5, the miRNA-hub gene regulatory with 2574 nodes (miRNA: 2300; hub gene: 274) and 18416 edges, and multiple hub genes associated with miRNAs. TUBB2A could be regulated by 206 miRNAs (ex; hsa-mir-4471), MAPK14 could be regulated by 124 miRNAs (ex; hsa-mir-7159-3p), STX7 could be regulated by 109 miRNAs (ex; hsa-mir-4478), RHOC could be regulated by 91 miRNAs (ex; hsa-mir-6818-5p), RAD18 could be regulated by 61 miRNAs (ex; hsa-mir-3150b-3p), YWHAZ could be regulated by 279 miRNAs (ex; hsa-mir-518f-5p), HSP90AA1 could be regulated by 188 miRNAs (ex; hsa-mir-133a-3p), TP53 could be regulated by 174 miRNAs (ex; hsa-mir-940), HSP90AB1 could be regulated by 162 miRNAs (ex; hsa-mir-411- 5p) and RPS15A could be regulated by 95 miRNAs (ex; hsa-mir-3666) (Table 5).

**Fig. 5.**
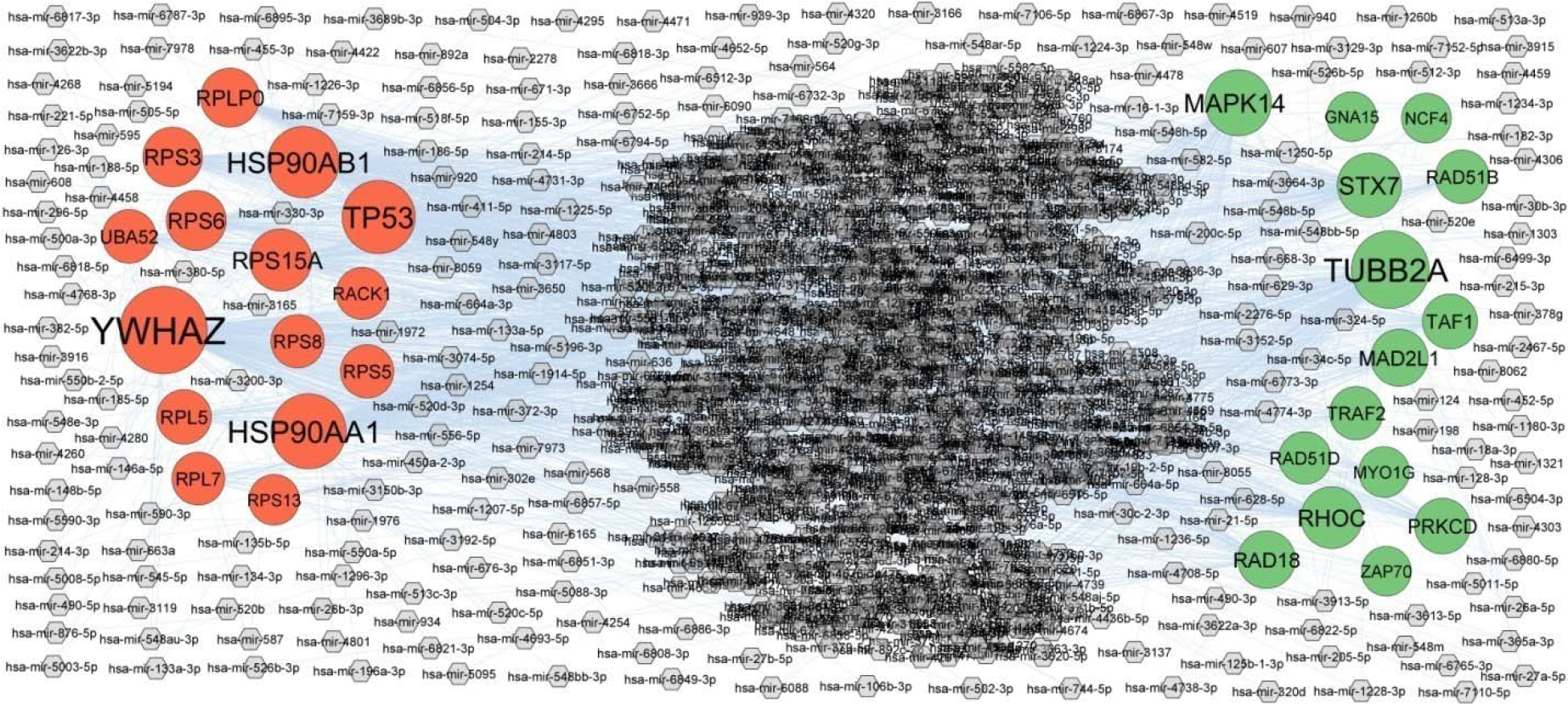
Target gene - miRNA regulatory network between target genes. The purpule color diamond nodes represent the key miRNAs; up regulated genes are marked in green; down regulated genes are marked in red.

**Table 5.**
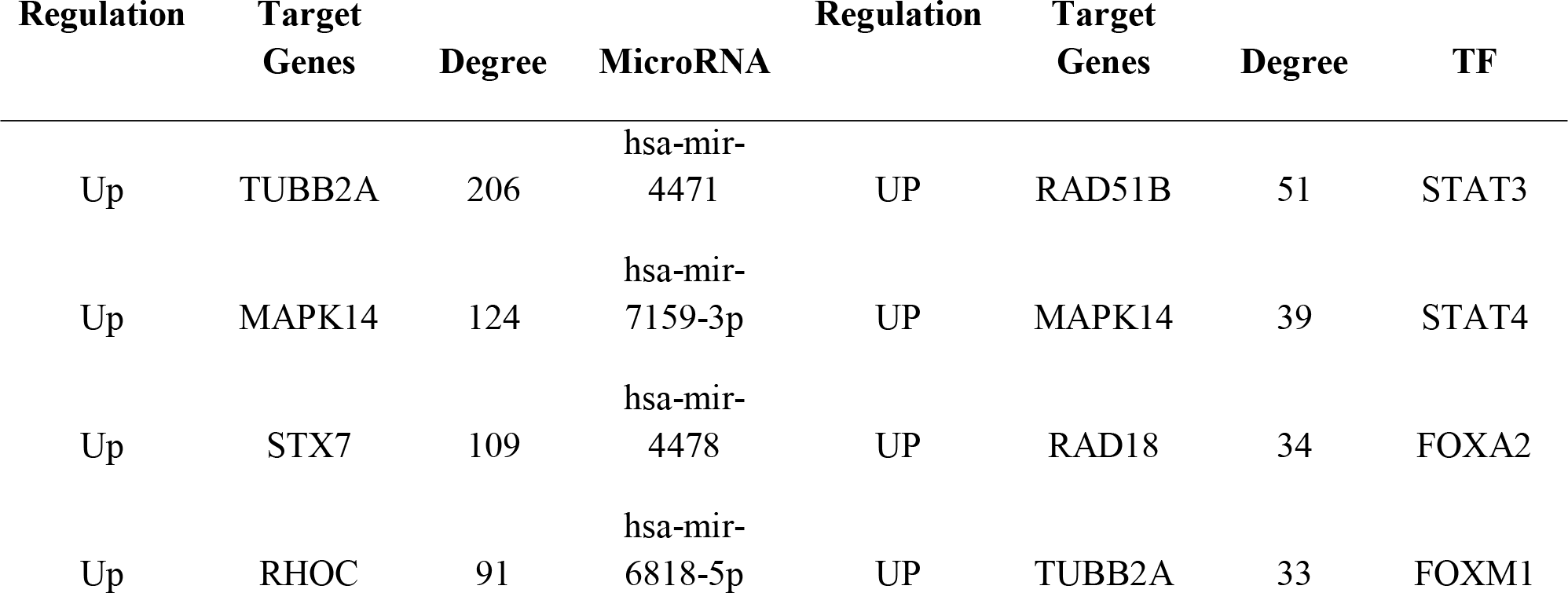
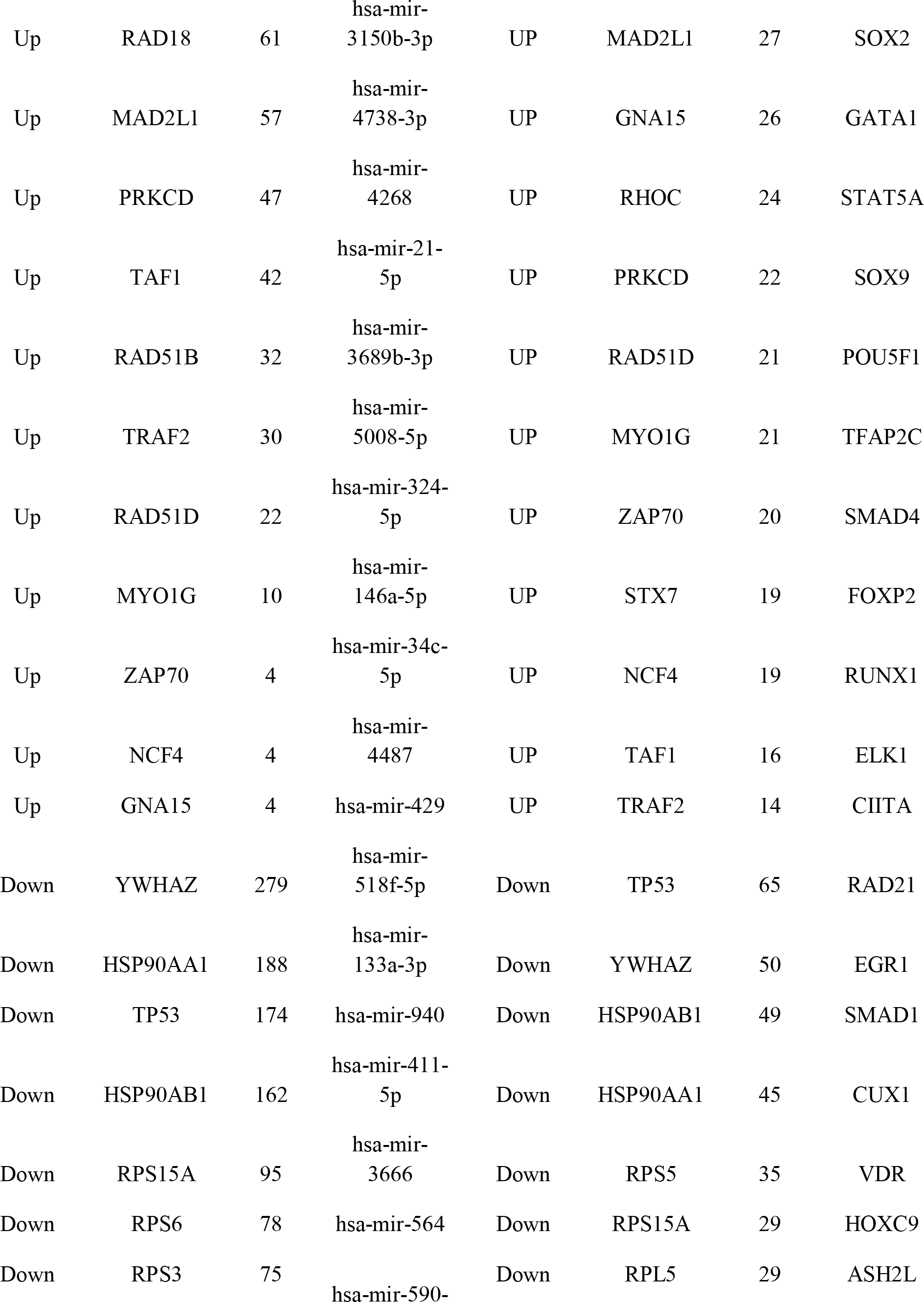
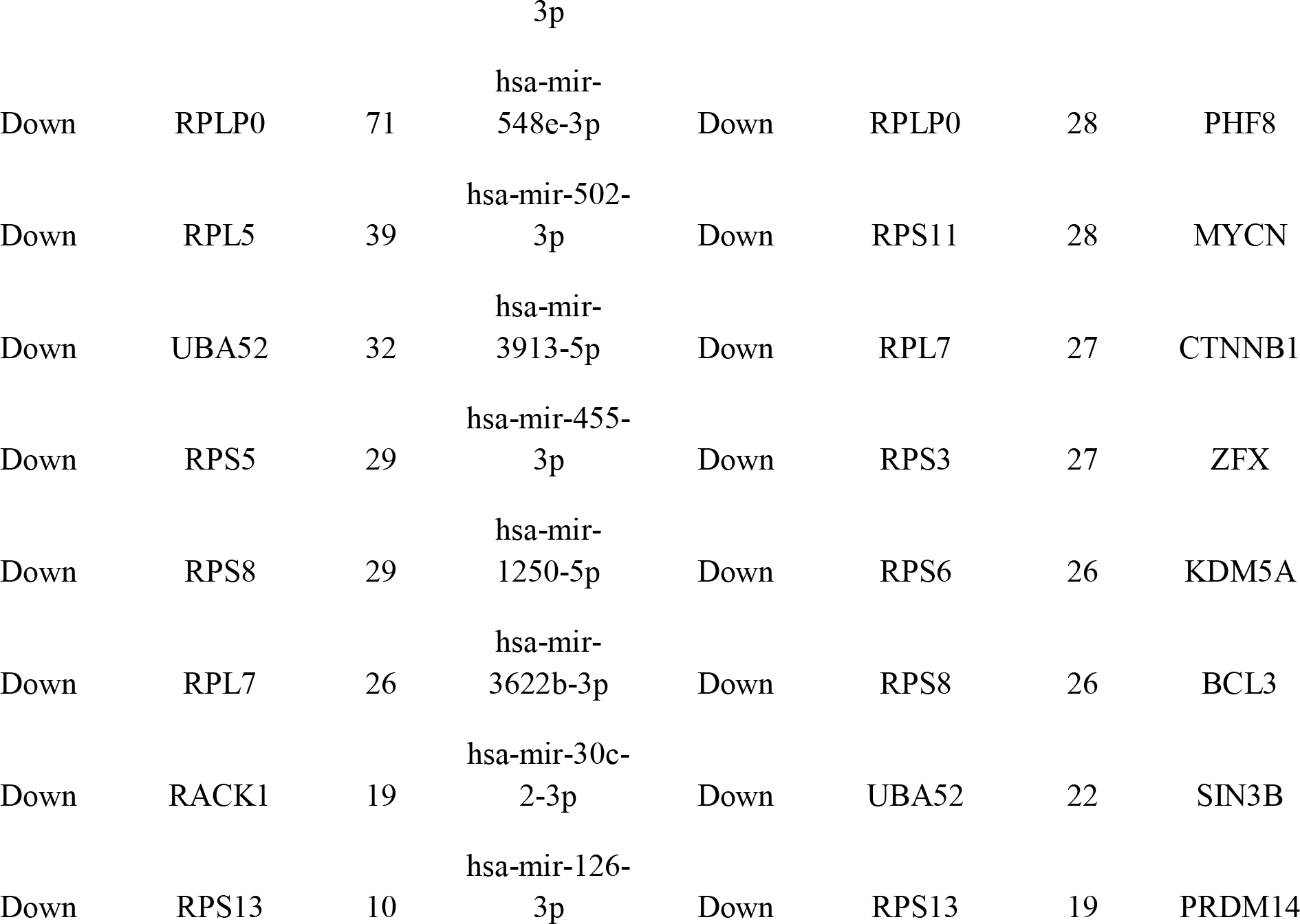
miRNA - target gene and TF - target gene interaction

### TF-hub gene regulatory network construction

TF-hub gene regulatory network constructed by NetworkAnalyst databasewas adjusted and visualized by Cytoscape. As shown in Fig 6, the TF-hub gene regulatory with 463 nodes (TF: 193; hub Gene: 270) and 7730 edges, and multiple hub genes associated with TFs. RAD51B could be regulated by 51 TFs (ex; STAT3), MAPK14 could be regulated by 39 TFs (ex; STAT4), RAD18 could be regulated by 34 TFs (ex; FOXA2), TUBB2A could be regulated by 33 TFs (ex; FOXM1), MAD2L1 could be regulated by 27 TFs (ex; SOX2), YWHAZ could be regulated by 50 TFs (ex; EGR1), HSP90AB1 could be regulated by 49 TFs (ex; SMAD1), HSP90AA1 could be regulated by 45 TFs (ex; CUX1), RPS5 could be regulated by 35 TFs (ex; VDR) and RPS15A could be regulated by 29 TFs (ex; HOXC9) (Table 5).

**Fig. 6.**
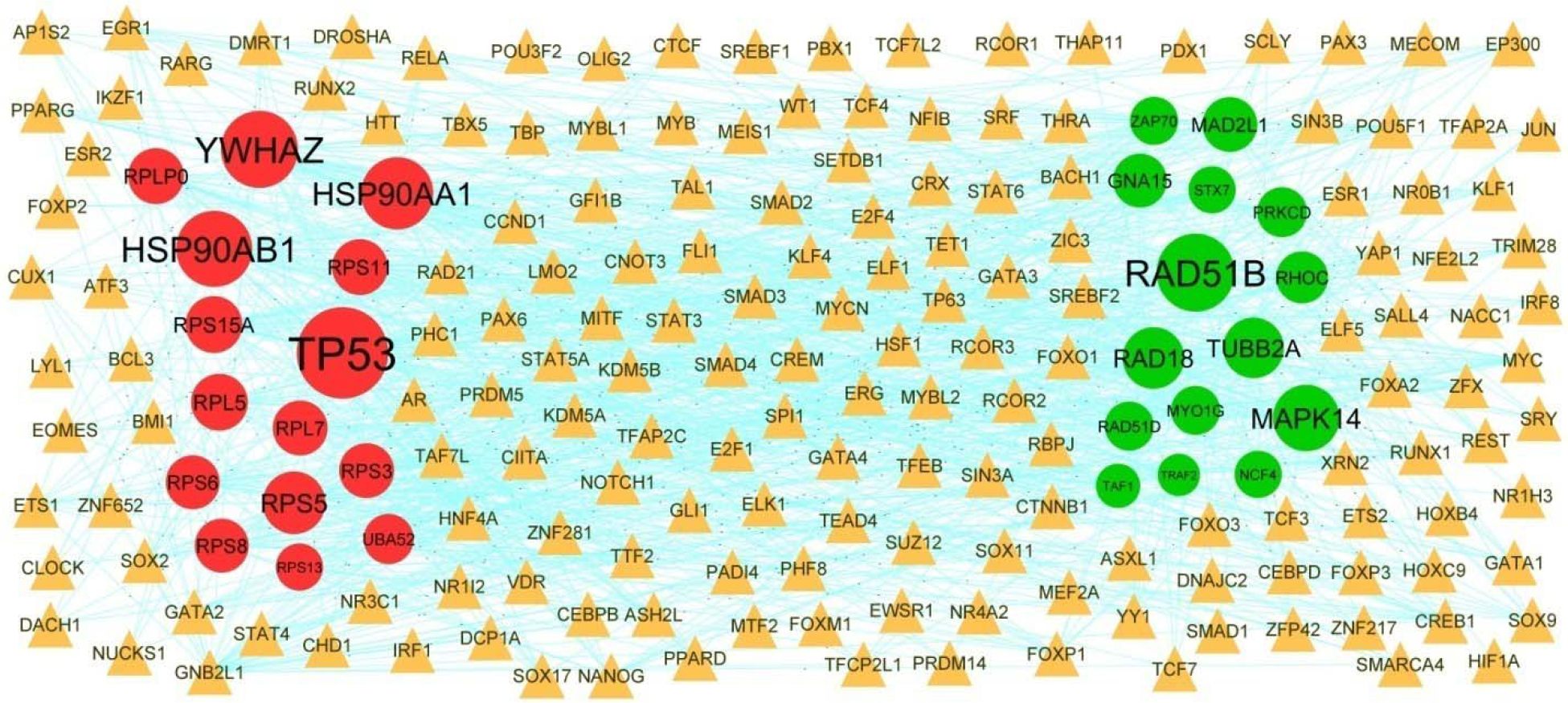
Target gene - TF regulatory network between target genes. The orange color triangle nodes represent the key TFs; up regulated genes are marked in green; down regulated genes are marked in red.

### Receiver operating characteristic curve (ROC) analysis

The ROC analysis of the ten hub genes expression of MAPK14, RHOC, MAD2L1, TAF1, TRAF2, HSP90AA1, TP53, HSP90AB1, UBA52 and RACK1 from PPI network, miRNA-hub gene regulatory network and TF-hub gene regulatory was performed. MAPK14, RHOC, MAD2L1, TAF1, TRAF2, HSP90AA1, TP53, HSP90AB1, UBA52 and RACK1 with AUC more than 0.80 were considered as hub genes (Fig 7), indicating that they have the capability to diagnose T1DM patients with excellent specificity and sensitivity.

**Fig. 7.**
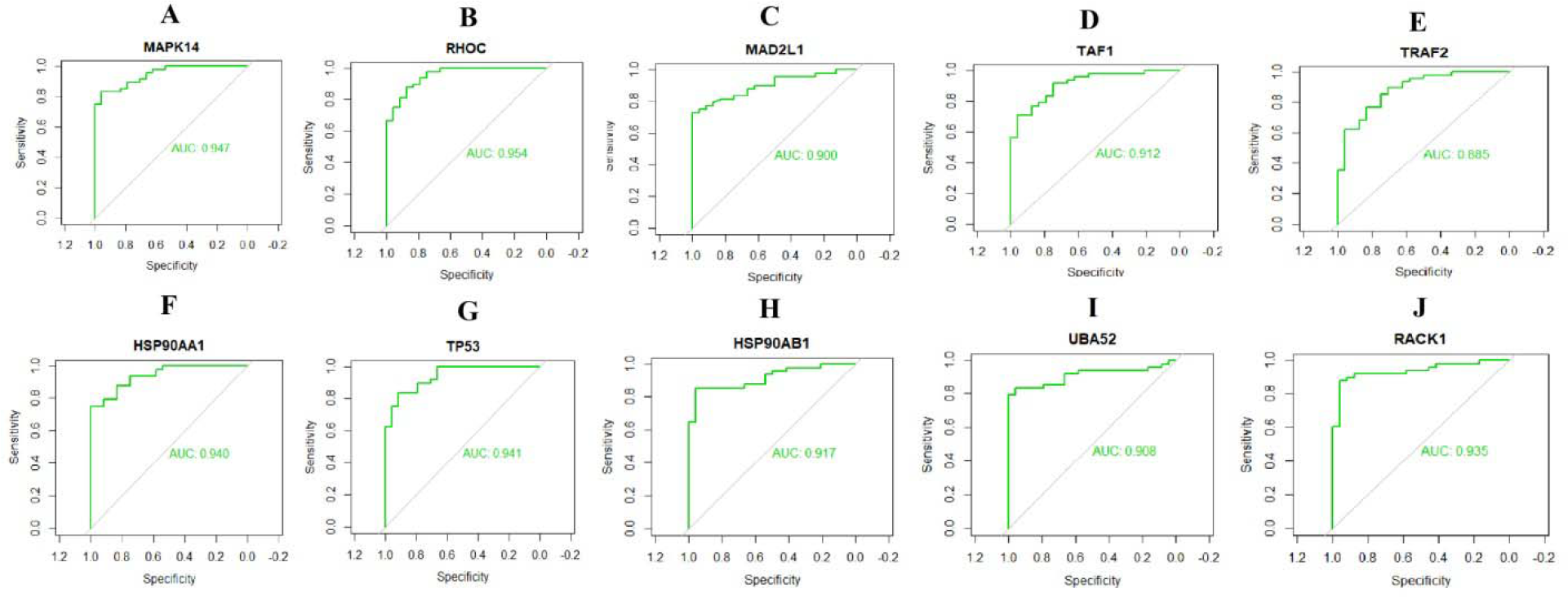
ROC curve analyses of hub genes. A) MAPK14 B) RHOC C) MAD2L1 D) TAF1 E) TRAF2 F) HSP90AA1 G) TP53 H) HSP90AB1 I) UBA52 J) RACK1

## Discussion

T1DM is a heterogeneous auto immune endocrine and metabolic disease with dismal prognosis [39]. If not treated promptly and effectively, T1DM can seriously reduce the quality of life and even cause serious other complication. There is no confusion that insight of diseases at the molecular level will help to better their diagnosis and treatment [40]. Up to now, various biomarkers have been identified to be associated with T1DM and might be selected as therapeutic targets, but the detailed molecular mechanism of gene regulation leading to disease advancement remains elusive [41].

In our investigation, we aimed to identify biomarkers of T1DM and uncover their biological functions through bioinformatics analysis. NGS dataset GSE182870 was selected in our analysis. As a result, 477 up regulated and 383 down regulated genes, at least 4-fold change between T1DM and normal control samples. Wang et al. [42], Mishra et al. [43], Deaton et al. [44] and You et al. [45] demonstrated that altered expression of SPHK1, WDR13, GIGYF1 and DNMT3A were found to be substantially related to type 2 diabetes mellitus. Altered expression of SPHK1 [46] was easily found in hypertension. Recently, increasing evidence demonstrated that SPHK1 [47] and DNMT3A [48] were altered expression in cognitive diseases. Gabriel et al. [49] reported that altered expression of SPHK1 could be an index for obesity. The involvement of SPHK1 [50] and WWTR1 [51] with renal diseases was demonstrated previously. Several studies have shown that altered expression of SPHK1 [52], ADTRP (androgen dependent TFPI regulating protein) [53] and DNMT3A [54] can be a strong prognosis biomarkers to predict relapse in patients with cardiovascular complications.

Enrichment analysis of GO and REACTOME pathway was carried out to explore the interaction between DEGs. Pathways include metabolism [55], neutrophil degranulation [56], metabolism of lipids [57], immune system [58], metabolism of proteins [59] and TP53 [60] regulates metabolic genes were linked with progression of T1DM. KDM7A [61], LMNA (lamin A/C) [62], PFKFB3 [63], NEU1 [64], DUSP22 [65], ADORA2B [66], CRTC2 [67], GRN (granulin precursor) [68], CTSD (cathepsin D) [69], IGFBP4 [70], SMYD2 [71], OSM (oncostatin M) [72], INVS (inversin) [73], ANKFY1 [74], SEMA4D [75], PPM1A [76], IFNG (interferon gamma) [77], CTBP1 [78], ATP6 [79], COX2 [80], RPS19 [81], COX1 [82] and TLK1 [83] are potential biomarkers for the detection and prognosis of renal diseases. A previous study reported that LXN (latexin) [84], LMNA (lamin A/C) [85], PFKFB3 [86], NEU1 [87], TBK1 [88], GRN (granulin precursor) [89], CTSD (cathepsin D) [90], ACADS (acyl-CoA dehydrogenase short chain) [91], IRF7 [92], S1PR1 [93], ZAP70 [94], IDH1 [95], IL15 [96], PIK3R1 [97], OSM (oncostatin M) [98], SOCS3 [99], USP21 [100], CEP19 [101], KDM2A [102], TP53 [103], BRD2 [104], ATP6 [105], BRD4 [106], COX2 [107], RPS6 [108], ND2 [109], CYTB (cytochrome b) [110] and COX1 [111] are altered expressed in obesity. Altered expression of BCL3 [112], TRAF2 [113], NEU1 [114], SNAP29 [115], AGPAT2 [116], LPCAT3 [117], ADORA2B [118], CTSD (cathepsin D) [119], ACADS (acyl-CoA dehydrogenase short chain) [120], ACAD9 [121], E4F1 [122], IRF7 [123], TAF1 [124], S1PR1 [125], RASSF1 [126], ELAC2 [127], RNF146 [128], COX15 [129], SMYD2 [130], IDH1 [131], MTO1 [132], IL15 [133], PIK3R1 [134], ASB1 [135], OSM (oncostatin M) [136], ZNF791 [137], GBA (glucosylceramidase beta) [138], SOCS3 [139], SLC39A7 [140], AKIP1 [141], AMIGO2 [142], GLUL (glutamate-ammonia ligase) [143], SEMA4D [144], KDM2A [145], TP53 [146], JARID2 [147], CTBP1 [148], ATP6 [149], RPL7 [150], HSP90AA1 [151], BRD4 [152], PSMB4 [153], COX2 [154], JUND (JunD proto-oncogene, AP-1 transcription factor subunit) [155], RPS5 [156], RACK1 [157], ND1 [158], CCND2 [159], COX1 [160], TLK1 [161] and TMPO (thymopoietin) [162] are associated with cardiovascular complications. A previous study found that LMNA (lamin A/C) [85], SLC11A2 [163], CRTC2 [164], TBK1 [165], GRN (granulin precursor) [166], CTSD (cathepsin D) [167], STARD10 [168], PGRMC1 [169], TFE3 [170], POR (cytochrome p450 oxidoreductase) [171], SESN1 [172], IL15 [173], PIK3R1 [134], OSM (oncostatin M) [98], SOCS3 [174], USP21 [100], GLUL (glutamate-ammonia ligase) [175], IL1R1 [176], TP53 [177], PPM1A [178], CTBP1 [179], DNAJC3 [180], ATP6 [181], DDX21 [182], COX2 [183], RACK1 [184], ND1 [158], CCND2 [185] and COX1 [186] are positively correlated with type 2 diabetes mellitus, suggesting its potential as a biomarker for type 2 diabetes mellitus. LMNA (lamin A/C) [187], PFKFB3 [188], TRAF2 [189], ADORA2B [190], ACADS (acyl-CoA dehydrogenase short chain) [191], IRF7 [192], ASL (argininosuccinatelyase) [193], IL15 [194], PIK3R1 [195], OSM (oncostatin M) [196], SOCS3 [197], CHD9 [198], IFI44L [199], GNLY (granulysin) [200], FLNB (filamin B) [201], IL1R1 [202], SETD3 [203], TP53 [204], BRD4 [205], COX2 [206], HSP90AB1 [207], SLC7A1 [208], ND2 [209] and COX1 [210] have been reported to be associated with hypertension. LMNA (lamin A/C) [211], INPP5K [212], SCYL1 [213], AKR7A2 [214], TRAF2 [215], SLC11A2 [216], NEU1 [217], SNAP29 [218], DUSP22 [219], P2RX4 [220], ADORA2B [221], NAXD (NAD(P)HX dehydratase) [222], FEZ1 [223], TBK1 [224], GRN (granulin precursor) [225], ATG4D [226], CTSD (cathepsin D) [227], PPP2R5D [228], IRF7 [229], ACAA1 [230], S1PR1 [231], PGRMC1 [169], MAPK14 [232], IL15 [233], OSM (oncostatin M) [234], GBA (glucosylceramidase beta) [235], SOCS3 [236], GLUL (glutamate-ammonia ligase) [175], MON1A [237], LMAN2L [238], RHOC (ras homolog family member C) [239], TUBB2A [240], CHP1 [241], TP53 [242], ATXN1 [243], JARID2 [244], IFNG (interferon gamma) [245], NUCKS1 [246], CTBP1 [247], DNAJC3 [180], ATP6 [248], BRD4 [249], EIF4B [250], COX2 [251], SFPQ (splicing factor proline and glutamine rich) [252], PCBP1 [253], RPL5 [254], HSP90AB1 [255], TRA2A [256], PIP4K2A [257], YWHAG (tyrosine 3-monooxygenase/tryptophan 5-monooxygenase activation protein gamma) [258] and SRSF6 [259] have been shown to be activated in cognitive diseases. DEAF1 [260], PFKFB3 [261], GRN (granulin precursor) [68], IRF7 [262], ZAP70 [263], TFE3 [170], IGFBP4 [70], IL15 [264], EDEM2 [265], SOCS3 [266], PHTF1 [267], IL1R1 [268], TP53 [269], IFNG (interferon gamma) [270], EIF2S3 [271], COX2 [272], IL2RG [273] and COX1 [186] were diagnostic biomarkers of T1DM and could be used as therapeutic targets. Enriched genes in these GO terms and pathways might found to be involved in the development of T1DM.

Construction of PPI network and modules of DEGs might be helpful for understanding the relationship of developmental T1DM. Altered expression of the TNFRSF12A [274] gene plays a role in the development of cardiovascular complications. The findings from the present investigation indicated that the MAD2L1, APBA3, TNFRSF10A, TNFRSF14, SNRPG (small nuclear ribonucleoprotein polypeptide G), RANBP2, SRRM1 and PABPN1 genes might be novel biomarkers for T1DM.

In this investigation, we also constructed a miRNA-hub gene regulatory network and TF-hub gene regulatory network for the hub genes. Mirza et al [275], Warshauer et al [276], Fichna et al [277], Zhang et al [278] and Abd-Allah et al [279] reported that the altered expression of the hsa-mir-133a-3p, STAT3, STAT4, SOX2 and VDR (vitamin D receptor) are correlated with T1DM. Recent studies have proposed that the hsa-mir-133a-3p [280], STAT3 [281], STAT4 [282], FOXA2 [283], FOXM1 [284], EGR1 [285], CUX1 [286] and VDR (vitamin D receptor) [287] are associated with obesity. Xie et al [288], Liu et al [289], Peng et al [290], Sarlak and Vincent, [291], Wang et al [292], Hegarty et al [293] and Liu et al [294] reported that altered expression of hsa-mir-411-5p, STAT3, FOXA2, SOX2, EGR1, SMAD1, CUX1 and VDR (vitamin D receptor) could be an index for cognitive diseases. A previous investigation found that altered expression of STAT3 [295], STAT4 [296], FOXA2 [297], SOX2 [298], EGR1 [299], SMAD1 [300] and VDR (vitamin D receptor) [301] were associated with type 2 diabetes mellitus. Bao et al. [302], Reid et al. [303], Tian et al. [304], Liang et al. [305], Fan et al. [306], Wang et al. [307] and Gonzalez-Parra et al. [308] showed that STAT3, STAT4, FOXM1, SOX2, EGR1, SMAD1 and VDR (vitamin D receptor) might play an important role in regulating the genetic network related to the occurrence and development of cardiovascular complications. The altered expression of STAT3 [309], FOXM1 [310], SMAD1 [311] and VDR (vitamin D receptor) [312] might be related to the progression of hypertension. Viau et al. [313], Bolin et al. [314], Zheng et al. [315], Xie et al. [316], Németh et al. [317], Ji et al. [318], Livingston et al. [319] and Yang et al. [320] studied the clinical and prognostic value of STAT3, STAT4, FOXA2, FOXM1, EGR1, SMAD1, CUX1 and VDR (vitamin D receptor) in patients with renal diseases.. In this investigation, STX7, RAD18, YWHAZ, RPS15A, RAD51B, hsa-mir-4471, hsa-mir-7159-3p, hsa-mir-4478, hsa-mir-6818-5p, hsa-mir-3150b-3p, hsa-mir-518f-5p, hsa-mir-940, hsa-mir-3666 and RAD21 were might be potential novel modulators of T1DM.

In conclusion, we used a series of bioinformatics analysis methods to identify the crucial genes and pathways associated in T1DM initiation and progression from NGS dataset containing T1DM samples and normal control samples. Our results provide a more detailed molecular mechanism for the advancement of T1DM, shedding light on the novel biomarkers and therapeutic targets. However, the interacting mechanism and function of genes need to be confirmed in further experiments.

## Acknowledgement

I thank Alex Ken Hu, Benaroya Research Institute, Systems Immunology, Seattle, WA, USA, very much, the author who deposited their NGS dataset GSE182870, into the public GEO database.

## Conflict of interest

The authors declare that they have no conflict of interest.

## Ethical approval

This article does not contain any studies with human participants or animals performed by any of the authors.

## Informed consent

No informed consent because this study does not contain human or animals participants.

## Availability of data and materials

The datasets supporting the conclusions of this article are available in the GEO (Gene Expression Omnibus) (https://www.ncbi.nlm.nih.gov/geo/) repository. [(GSE182870) https://www.ncbi.nlm.nih.gov/geo/query/acc.cgi?acc=GSE182870)]

## Consent for publication

Not applicable.

## Competing interests

The authors declare that they have no competing interests.

### Author Contributions

B. V. - Writing original draft, and review and editing

C. V. - Software and investigation

## Notes

### Competing Interest Statement

The authors have declared no competing interest.

